# Beyond snapshots: A dynamic framework for continuous motor unit tracking to enhance neurophysiological assessments

**DOI:** 10.64898/2026.09.25.754489

**Authors:** Subaryani Dambawati H. Soedirdjo, Aleš Holobar, Yasin Y. Dhaher

## Abstract

Tracking individual motor units (MUs) across multiple testing visits is critical for understanding neuromuscular adaptations and disease progression. However, existing methods fail to account reliably for spatiotemporal variations in motor unit action potentials (MUAPs) and lack validation for longitudinal tracking. Here, we present a robust computational framework for continuous MU tracking that integrates automatic estimation of electrode grid displacements and compensation for MUAP shape variations. Our algorithm demonstrated superior tracking accuracy, capturing 63% and 75% of MUs in the tibialis anterior and medial gastrocnemius, respectively, substantially outperforming state-of-the-art methods. By applying this approach to investigate estradiol-driven neuromuscular plasticity as a model system, we reveal previously undetectable modulations in recruitment thresholds and firing rates, linking hormonal fluctuations to motor unit behavior. Beyond advancing fundamental neuromuscular research, our approach holds promise for enhancing biomarker sensitivity in clinical trials, particularly in conditions such as amyotrophic lateral sclerosis, where reliable MU tracking is paramount. These findings establish a new standard for non-invasive motor unit tracking, unlocking opportunities for longitudinal neurophysiological assessment and evaluation of therapeutics.

---

The neuromuscular function is profoundly influenced by pathological conditions such as motor neuron diseases,^1,2^ age-related degeneration,^3^ medical interventions,^4,5^ physical training,^6^ and environmental factors such as microgravity.^7^ These factors progressively alter properties and behavior of motor unit (MU), the fundamental functional entity of the peripheral motor system comprises of a motoneuron and its innervated muscle fibers; necessitating longitudinal assessments to monitor neuromuscular integrity and adaptation.

MU function is defined by key neurophysiological parameters, including recruitment threshold, firing rate, and the number of active units. Current methods primarily rely on population-level MU comparisons, which may obscure MU-specific adaptive or pathological responses. This limits mechanistic insight into disease progression and therapeutic response, underscoring the need for precise longitudinal MU tracking in clinical and research settings.^8^

Needle electrodes are traditionally used in clinical practice to evaluate MUs. This technique is invasive and produces MU action potential (MUAP) shapes that vary with electrode position within the muscle,^9^ making it challenging to examine the same MU across testing visits. To overcome this drawback, a noninvasive High-Density surface Electromyography (HDsEMG) electrode grid has been used to infer MU characteristics during muscle contraction by exploiting the differences in MUAP distribution on the skin surface.^10^

One approach to identifying MUs across visits using HDsEMG is to electrically stimulate the motor axon and define identical MUs across visits based on their excitation thresholds.^11,12^ While this technique estimates the MU number, it does not provide information about the firing behavior or central motor drive, making it unsuitable for evaluating tasks involving voluntary contractions. Early attempts to follow individual MU during voluntary contraction are based on MUAP shape evaluation or applying the pre-calculated MU filters to new signals.^13–15^ Because MUAP shape depends on the relative distance between the source and the detection electrode, misalignment of the HDsEMG grid during multiple testing sessions can alter the recorded MUAP morphology. Consequently, these techniques are suboptimal for longitudinal MU tracking because they do not compensate for MUAP shape changes. A single study that attempted to follow individual MU by compensating MUAP shape implemented the approach on a controlled electrode grid displacement.^16^ In the experimental setting, the amount of displacement is unknown; hence, a method to automatically measure the displacement is essential, and a computer-aided processing pipeline that automatically identifies the grid displacement and compensates for its effect when tracking individual MUs is required but currently lacking. Moreover, the sensitivity of MU tracking to grid displacement has not been systematically assessed.

Here, we present an image registration-based MU tracking algorithm and its processing pipeline addressing the aforementioned issues to enable tracking of individual MUs across testing visits (Fig. 1). Unlike prior approaches, our proposed algorithm integrates automatic estimation of HDsEMG grid displacement, compensation of MU morphological changes, and validation of the tracked MU. State-of-the-art (SoA) approach^17^ uses the MU filter calculated directly from the HDsEMG signal recorded at the first testing visit and applied it directly to subsequent HDsEMG signals to extract MU characteristics (without MUAP shapes compensation). We show that the proposed algorithm outperforms SoA method in the number of accurately tracked MUs on tibialis anterior (TA) and medial gastrocnemius (MG) muscles collected at two testing visits separated by 12-16 day. Using synthetic grid displacement, we quantify the grid displacement estimation error and tracking accuracy.

**Fig. 1:**
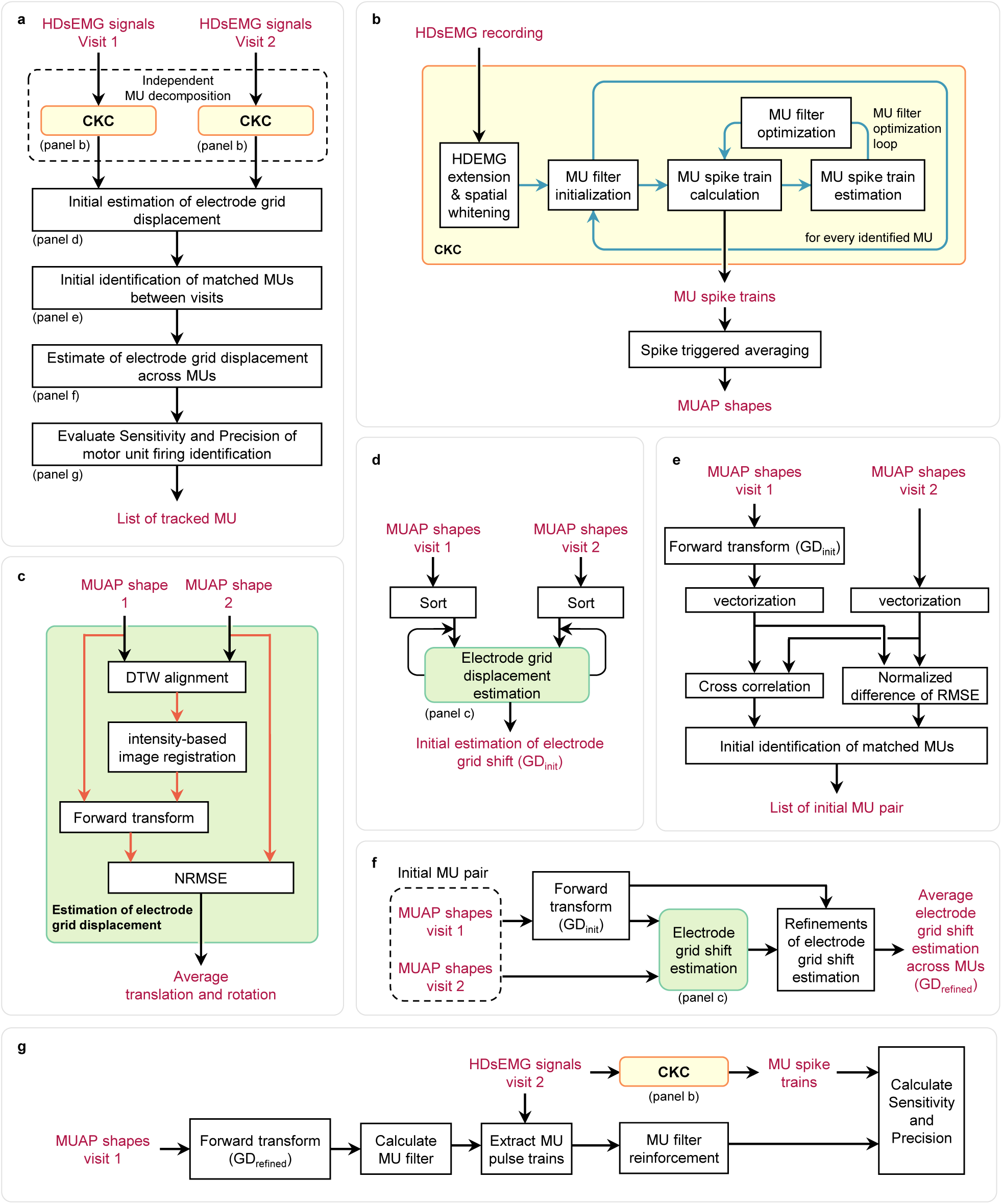
Pipeline of the MU tracking across visits. **a**, The overall workflow to track motor unit (MU) across testing visits. **b**, Detailed process of the decomposition of HDsEMG signal using Convolution Kernel Compensation (CKC) algorithm. **c**, Estimation of HDsEMG grid displacement using intensity-based image registration on dynamic time-warped (DTW) aligned samples. Normalized root mean square error (NRMSE) assessed the electrode grid displacement estimation quality. **d**, Detailed process of initial estimation of electrode grid displacement. The initial estimation of the electrode grid displacement was performed starting from the MU with the largest peak-to-peak MUAP amplitude. The estimation process was iterated until the estimation was obtained. **e**, Detailed process of identifying the initial match of MU between visits. **f**, The average electrode grid displacement across MU was obtained by calculating the average of the translation and rotation of each MU. **g**, Detailed process of calculating sensitivity and precision of motor unit firing identification.

Finally, we demonstrate the performance of the proposed algorithm in a case study of evaluating the effect of estradiol fluctuation on MU characteristics, showing its potential to clinical use.

## RESULTS

We developed an algorithm and its processing pipeline that accurately tracked MUs across testing visits (Supplementary Note 3). To understand the workflow of the proposed algorithm, we examined the intermediate and final outputs generated at each step. The recorded signals (Fig. 2a) were independently decomposed into their MUs constituent and MUAP images were formed (Fig. 2b). Then, the algorithm estimated the translations and rotation of the dynamic time warping aligned MUAP images (Fig. 2c), vectorized the compensated MUAP shape to evaluate shape similarity (Fig. 2d), and ultimately identified the matched pairs of MUs. During initial identification of matched MUs (Fig. 1a), each MU may have multiple matches which were identified and resolved accordingly. Our proposed algorithm transformed the MUAP spatiotemporal shape at Visit 1, resulting in a shape similar to that of Visit 2 (Fig. 2e).

**Fig. 2:**
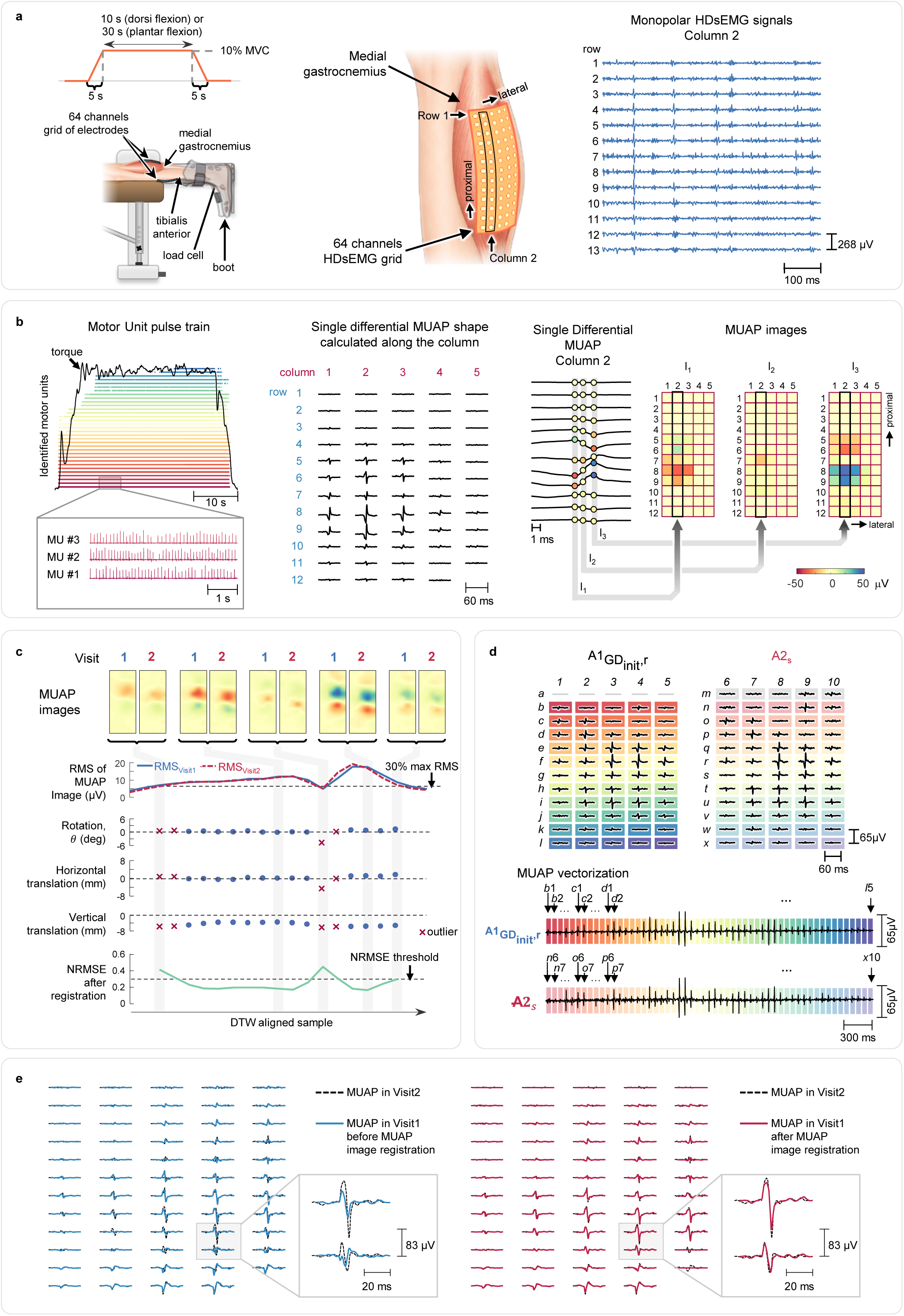
Process overview of the algorithm to track MU. **a**, MG and TA monopolar HDsEMG signals were collected following a trapezoidal torque trajectory during isometric dorsi and plantar flexion. **b**, The Convolution Kernel Compensation algorithm was employed to estimate MU pulse trains (Supplementary Note 1). Spike-triggered averaging was used for each identified MU to calculate the MUAP spatiotemporal shapes. MUAP images of each time sample were obtained by taking the instantaneous MUAP amplitude of each electrode and arranging them in a matrix according to the location of the corresponding electrode on the HDsEMG grid. **c**, Examples of electrode grid displacement estimation of two pairs of MUs using the MUAP image registration process (Supplementary Note 3). After time series alignment using the dynamic time warping (DTW) algorithm, intensity-based image registration was used to estimate the electrode grid displacement. Translation and rotation values that yielded a normalized root mean square error (NRMSE) less than 0.3 were averaged. The closed blue circles and red crosses indicate a DTW-aligned sample with NRMSE < 0.3 and > 0.3 (outliers), respectively. **d**, Vectorized MUAP obtained by concatenating non-empty electrodes row-wise. Vectorized MUAP was used to evaluate the similarities of the MUAP shape using cross-correlation. Gray lines in A1_GD_*_init_*_,*r*_ and gray shades in A2*_s_* indicate missing electrodes of Visit 1 and discarded electrodes of Visit 2 after the initial estimation of electrode grid displacement, respectively. **e**, MUAP shape transformation after application of the proposed algorithm. The dashed lines are the MUAP spatiotemporal shapes of the MU in Visit 2. Solid lines on the left and right side of the panel denote the MUAP spatiotemporal shape obtained using SoA algorithm in Visit 1 and after MUAP image registration, respectively. The proposed algorithm improves the similarity of MUAP spatiotemporal shapes between Visit 1 and Visit 2.

### Dataset for performance evaluation

To evaluate and compare the performance of our algorithm with SoA method,^17^ we utilized MUs identified in each of two testing visits for TA (14 female participants) and MG (11 female participants) muscles during 10% maximal voluntary contraction (MVC) isometric dorsiflexion and plantarflexion, respectively. The independent decomposition (Fig. 1b) identified in TA muscle (mean ± SD) 14 ± 7 MUs in Visit 1 and 13 ± 5 MUs in Visit 2 (*p* = 0.439). Similarly, 14 ± 8 and 13 ± 8 MUs were identified independently from the MG muscle (11 participants, p = 0.457) at Visits 1 and 2, respectively. Between the two visits, we observed no significant change in the TA MVC (31.7 ± 7.9 Nm and 32.5 ± 8.2 Nm, p = 0.456) and pulse to noise ratio (PNR)^18^ of the identified MUs of TA muscle (34.5 ± 2.9 and 34.8 ± 3.4 dB, p = 0.756). The MG MVC (68.0 ± 16.2 Nm and 68.6 ± 15.5 Nm, p = 0.833) and PNR of the identified MUs of MG muscle (36.2 ± 4.2 dB and 36.4 ± 4.3 dB, p = 0.833) were also not significantly different. These results indicated that the quality of the identified MUs was similarly high in both visits and consistent muscle activation level across visits.

### Electrode grid displacement estimation errors

Synthetic displacement of MUAPs provided a controlled benchmark for evaluating the accuracy of grid displacement estimation. We synthetically displaced the identified MUAPs for one inter-electrode distance (8 mm) vertically and quantified the absolute error between the grid displacement estimated by the proposed algorithm and ground-truth displacement. In the TA muscle, our proposed algorithm showed a minimal estimation error of horizontal and vertical translations (0.35 ± 0.29 mm and 0.26 ± 0.12 mm), and rotation (0.15 ± 0.14 degrees). Similarly, in the case of the MG muscle, the average horizontal and vertical translations and rotation estimation errors were 0.42 ± 0.17 mm, 0.31 ± 0.10 mm, and 0.08 ± 0.09 degrees.

### Improvement of MU tracking performance on synthetically displaced MUAP

To evaluate the performance on tracking MUs, we applied our proposed algorithm and the SoA method on the synthetically displaced MUAP described above. Subsequently, we compared the number of tracked MU, quantified the quality of resulting firing rate using PNR, and compared MU firing identification sensitivity and precision.^17^

During the initial identification of matched MUs, 40% of the MUs in the TA and 35% of MG MUs had multiple matches. After resolving these matches, results show that our proposed algorithm accurately tracks, on average, 99% of the MUs in the TA and MG muscles and outperforms the SoA method (2% and 30% of the MUs in TA and MG muscles, respectively).

Bayesian analysis (Fig. 3c-d) revealed that, when tracking 28 sets of TA MUs, the proposed algorithm demonstrated superior performance in enhancing the MU pulse train PNR (median 34.1 dB [95% credible interval, 32.6-35.6 dB]) compared to the SoA method (17.8 dB [17.5-18.2 dB]). Additionally, relative to the SoA method, it exhibited higher sensitivity (98% [97-98%] vs 21% [18-24%]) and precision (95% [94-96%] vs 30% [26-33%]).

**Fig. 3:**
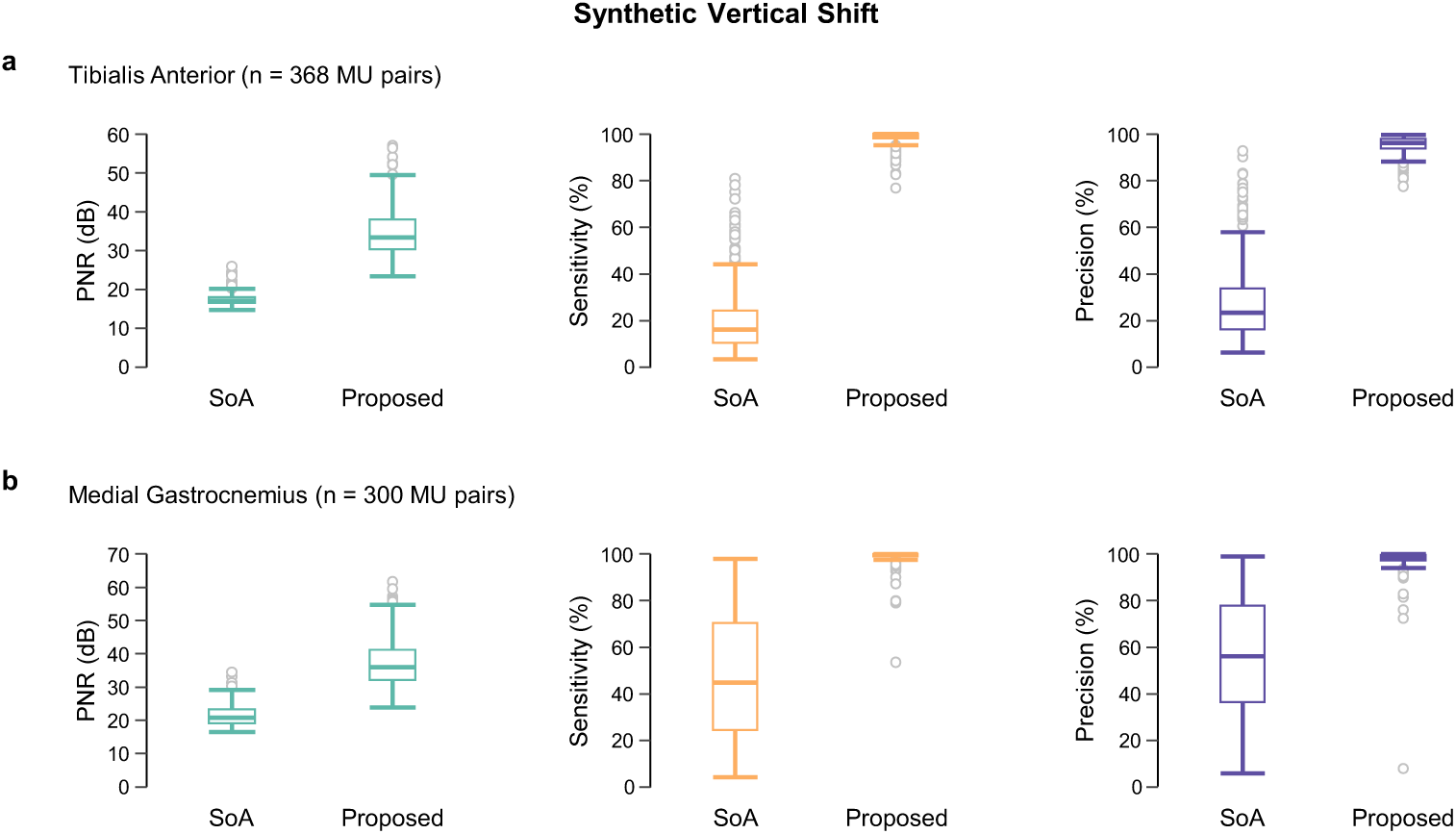
Synthetic 1 inter-electrode distance (8 mm) vertical electrode grid displacement. Comparison of Pulse-to-Noise Ratio (PNR), MU firing sensitivity, and MU firing precision of identified MUs are shown for each **a,** tibialis anterior and **b,** medial gastrocnemius muscle obtained by the proposed algorithm and the state-of-the-art (SoA) algorithm. Open circles indicate outliers.

Similarly, when tracking 22 sets of MG MUs, the proposed algorithm outperformed the SoA method in PNR (35.9 dB [33.6-38.2 dB] vs 22.3 dB [21.4-23.2 dB]), and sensitivity (98% [97-99%] vs 53% [47-60%]). The proposed algorithm also exceeded the precision of the SoA method (96% [95-97%] vs 60% [55-66%]).

Results show that, in both muscles, the proposed algorithm credibly improved PNR, sensitivity, and precision over SoA method: the posterior distribution excluded zero difference, and less than 2.5% fell within the region of practical equivalence (ROPE).

### Comparison to SoA method on across visits MU tracking

To assess MU tracking performance under a practical condition, we applied the proposed algorithm and the SoA method to track MUs between two visits. The HDsEMG grid location was measured and marked at the first testing visit. During MU tracking of the TA muscle, the proposed algorithm was unable to obtain an initial estimation of the electrode grid displacement in one participant and failed to accurately track all MUs in another. Similarly, during MU tracking of the MG muscle, the proposed algorithm failed to obtain an initial estimation of the electrode grid displacement in one participant. Data from these participants was excluded from subsequent analyses.

The proposed algorithm estimated absolute inter-visit grid displacement on TA of 4.45 ± 3.51 mm horizontally, 10.05 ± 4.98 mm vertically, and 2.77 ± 1.92 degrees in rotation, and 9.89 ± 7.87 mm, 11.73 ± 9.08 mm, and 1.03 ± 0.68 degrees, respectively, for inter-visit grid displacement on MG.

During the initial identification of matched MUs, on average, 37% of TA MUs and 46% of MG MUs had multiple matches, which were resolved accordingly. On average, the proposed algorithm accurately tracked 63% of MUs in the TA muscle of 12 participants, significantly outperforming the SoA method (7%). For the MG muscle of 10 participants, the proposed algorithm accurately tracked 75% of MUs on average, compared to 32% achieved by the SoA method.

In tracking TA MUs across visits (Fig. 4), our proposed algorithm achieved higher PNR (33.2 dB [31.0-35.0 dB]) compared to the SoA method (18.3 dB [17.2-19.6 dB]) and exceeded the sensitivity (91% [88-94%] vs 23% [15-33%]). Similarly, the precision of our proposed algorithm was significantly higher than the SoA method (91% [88-93%] vs 31% [22-41%]).

**Fig. 4:**
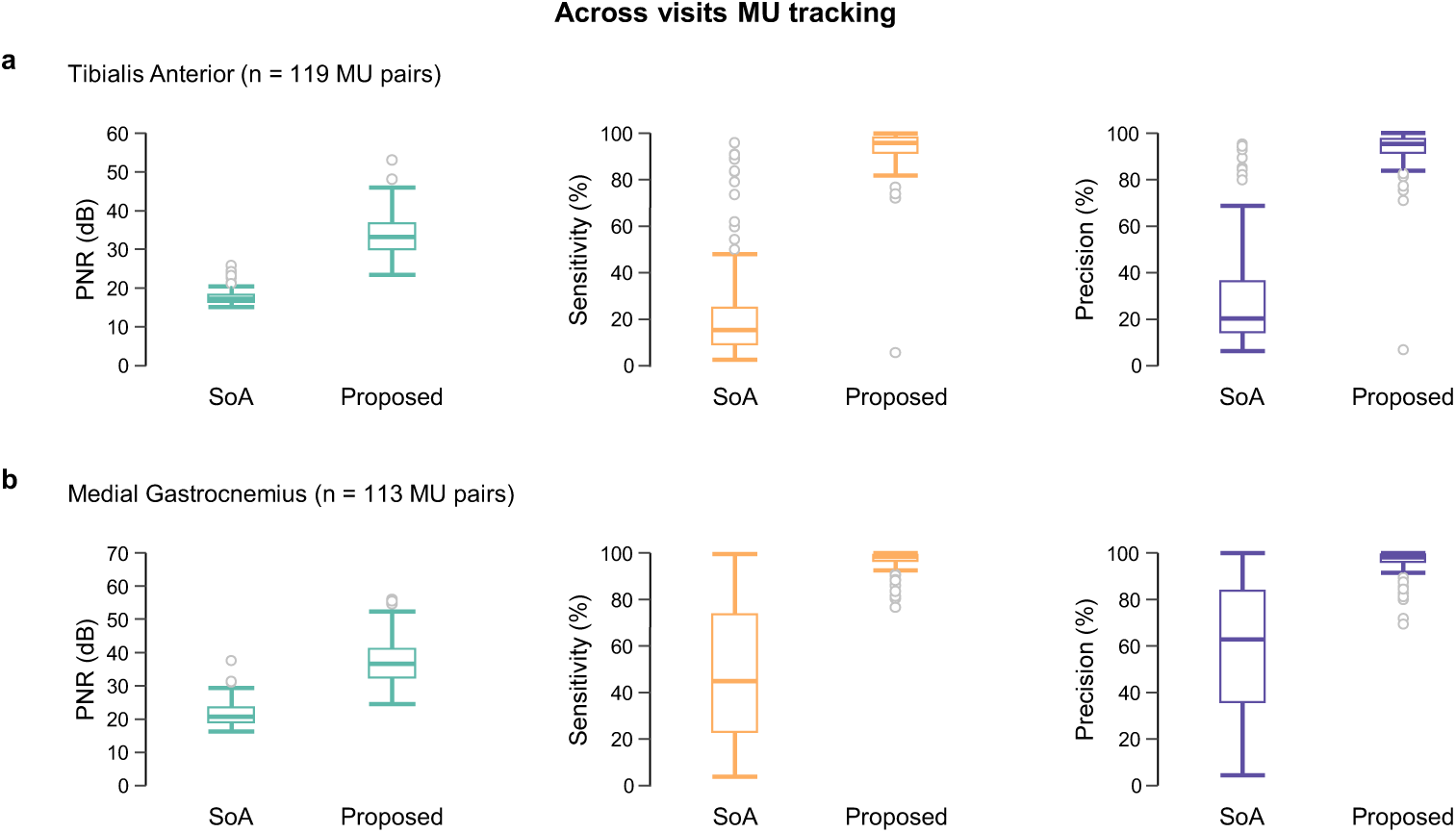
PNR, sensitivity, and precision of identified MU firing in MU tracking across visits. Comparison of Pulse-to-Noise Ratio (PNR), MU firing sensitivity, and MU firing precision of MUs on **a**, tibialis anterior and **b**, medial gastrocnemius muscles obtained by the proposed algorithm and the state-of-the-art (SoA) algorithm.

Similarly, when tracking MUs across visits on MG muscle, our proposed algorithm achieved higher PNR (35.3 dB [30.6-39.6 dB]) compared to the SoA method (22.6 dB [20.7- 25.0 dB]) and higher sensitivity than the SoA method (96% [94-97%] vs 53% [40-65%]).

Likewise, the precision obtained with the proposed algorithm (95% [93-97%]) was higher than the SoA method (58% [49-67%]).

In tracking MUs across visits of TA and MG muscles, our proposed algorithm demonstrated a superior performance compared to the SoA method on PNR, precision, and sensitivity: the posterior distribution excluded zero difference, and < 2.5% within ROPE.

### Algorithm runtime performance

We quantified the runtime of the proposed algorithm to track MUs across testing visits processed on a workstation equipped with a central processing unit (AMD Ryzen 5 3600, 32 GB of RAM). The proposed algorithm achieved an average running time ranging from 2.73 to 59.17 seconds with a median of 10.14 seconds. On average, 41.8% of the processing time was spent on the initial estimation of the electrode grid displacement, 52.1% on the electrode grid displacement across MUs, and 6.1% on the identification of the matched MU across two visits and other processes. These results show that the proposed algorithm can process data within a timeframe compatible with clinical and research use.

### Case study: MU characteristic modulation with increase in estradiol

We examined a representative case to demonstrate the capability of the proposed algorithm in a research setting. Using performance-evaluation’s protocol, we recorded HDsEMG signals from TA of nine female participants in three randomized testing visits: two during early follicular phase (EF1 and EF2) and one at peri-ovulatory (OV) and compared MU firing rate (FR) measured as pulse per second (pps) and recruitment threshold (RT) measured as percentage to MVC (%MVC).

The menstrual phases of eight participants were confirmed using the serum estradiol (EF1: 41 ± 19 pg/ml; EF2: 56 ± 20 pg/ml; OV: 314 ± 103 pg/ml), and progesterone concentrations (EF1: 0.5 ± 0.2 ng/ml; EF2: 0.4 ± 0.2 ng/ml; OV: 0.7 ± 0.3 ng/ml). Holm-adjusted paired t-test showed that estradiol concentrations were comparable between EF1 and EF2 (p = 0.239) but significantly lower at both than at OV (p < 0.001), and estradiol concentrations at EF2 was significantly lower than OV (p < 0.001). The progesterone concentrations were substantially lower than the reported maximum concentration in the luteal phase.^19^ Although a Friedman test showed a significant difference (χ²(2) = 6.30, p = 0.043), post-hoc Wilcoxon signed-rank test with Holm correction did not identify any statistically significant pairwise differences between sessions (p > 0.05).

The MVC were not significantly different EF1 33.6 ± 12.8 Nm, EF2 34.8 ± 11.1 Nm, and OV 33.7 ± 11.6 Nm (F(1.11, 8.91) = 0.63, p = .464, generalized η² = .002); confirming consistent muscle activation level across three visits.

The independent HDsEMG decomposition identified 130 MUs, 145 MUs, and 124 MUs at EF1, EF2, and OV visits, respectively. Total number of MUs between visits were not significantly different (F(2, 16) = 1.44, p = 0.265, generalized η² = .03). The average PNR were also similar between EF1 35.9 ± 3.6 dB, EF2 34.6 ± 2.7 dB, and OV 34.8 ± 2.0 dB (χ²(2) = 1.56, p = 0.459).

Our proposed algorithm accurately tracked 64 MUs across the three testing visits (on average, 56% of the lowest total number of the identified MUs across visits). A linear mixed-effects model (LMM) analysis of variance showed no evidence that FR differed significantly across EF1, EF2, and OV (F(2, 124) = 1.738, p = 0.180). Between EF1 and EF2, 71% of MUs showed invariant FR (±1 pps), 8% of MUs showed a decrease in FR (< -1 pps), and 21% of MUs showed an increase in FR (> 1 pps). Similarly, between EF2 and OV, 62% of MUs have invariant FR, 24% of MUs showed a decrease in FR, and 14% of MUs showed an increase in FR (Fig. 5a).

**Fig. 5:**
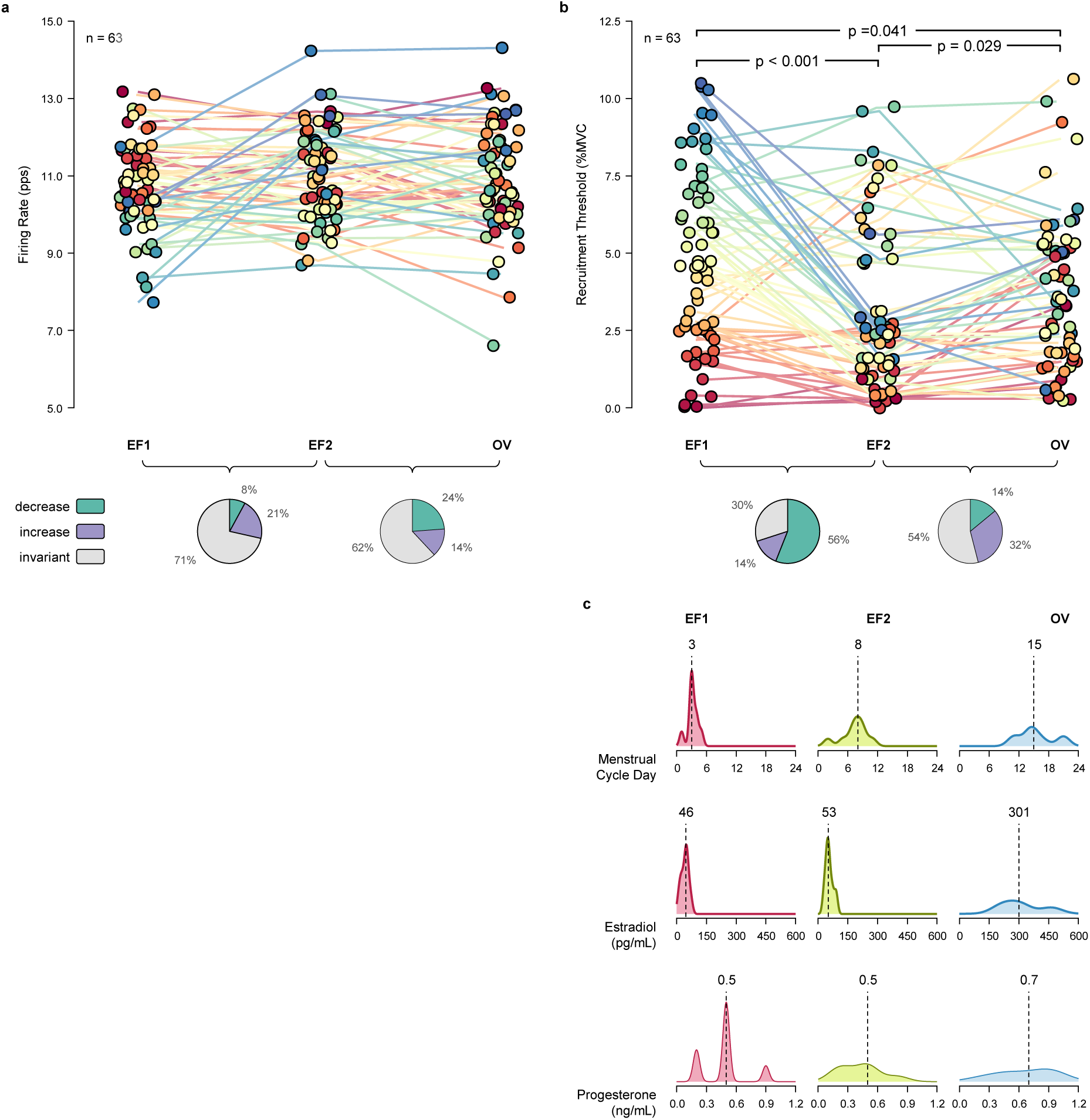
Effect of acute increase in estradiol concentration on mean firing rate and recruitment threshold of tracked Mus. **a**, The mean firing rates across three visits (n = 63) did not change significantly across three testing visits (EF1, EF2, and OV). A different color indicates each tracked MU. **b**, Recruitment thresholds in EF1 were significantly different from those in EF2 (p < 0.001) and those in OV (p = 0.041). A significant difference was also observed between RT in EF2 and OV (p = 0.029). **c**, The vertical dashed line indicates the median value of the distribution. Estradiol and progesterone concentrations were measured from 8 participants. The estradiol concentrations were significantly higher at OV compared to EF1 and EF2. Data is shown on the original outcome scale. Statistical comparison on firing rate was performed on the non-transformed. Statistical comparison on recruitment threshold was performed on square-root-transformed model scale.

Interestingly, the LMM fitted to square-root-transformed RT showed a significant effect of visits (Fig. 5b, F(2, 124) = 12.68, p < 0.001). For interpretability, estimated marginal means are presented on the original outcome scale after back-transformation. Pairwise tests on the model scale revealed that RT at EF1 was significantly higher than EF2 (Δ√RT = 0.451, t(df) = 5.036, adjusted p-value < 0.001, Cohen’s d = 0.897) and OV (Δ√RT = 0.219, t(df) = 2.448, adjusted p-value 0.041, Cohen’s d = 0.436). However, RT at EF2 was lower than at OV, Δ√RT = -0.232, t(df) = -2.588, adjusted p-value 0.029, Cohen’s d = -0.461. Between EF1 and EF2, 30% of MUs showed invariant RT (±1 %MVC), 56% of MUs showed a decrease in RT (< -1 %MVC), and 14% of MUs showed an increase in RT (> 1 %MVC). Between EF2 and OV, 54% of MUs have invariant RT, 14% of MUs showed a decrease in RT, and 32% of MUs showed an increase in RT.

We repeated the analyses using all identified MUs in three testing visits. Because MU were not tracked across visits, repeated observations from the same MU could not be identified. A MU-level random effect was therefore not included. The LMM analysis revealed that there was no evidence that FR differ significantly across Visits (Fig. 6a, F(2, 384.29) = 2.494, p = 0.084), but a significant fixed effect of Visits on square-root-transformed RT was observed (Fig. 6b, F(2, 383.28) = 4.367, p = 0.013). On the model scale, Tukey-adjusted pairwise comparisons showed that RT was significantly lower at EF2 (Δ√RT = 0.228, t(df) = 2.413, p = 0.043, Cohen’s d = 0.294) and OV (Δ√RT = 0.264, t(df) = 2.688, p = 0.020, Cohen’s d = 0.341) than at EF1, with no difference between EF2 and OV (Δ√RT = 0.036, t(df) = 0.374, p = 0.926, Cohen’s d = 0.046) unlike the analysis using tracked MU.

**Fig. 6:**
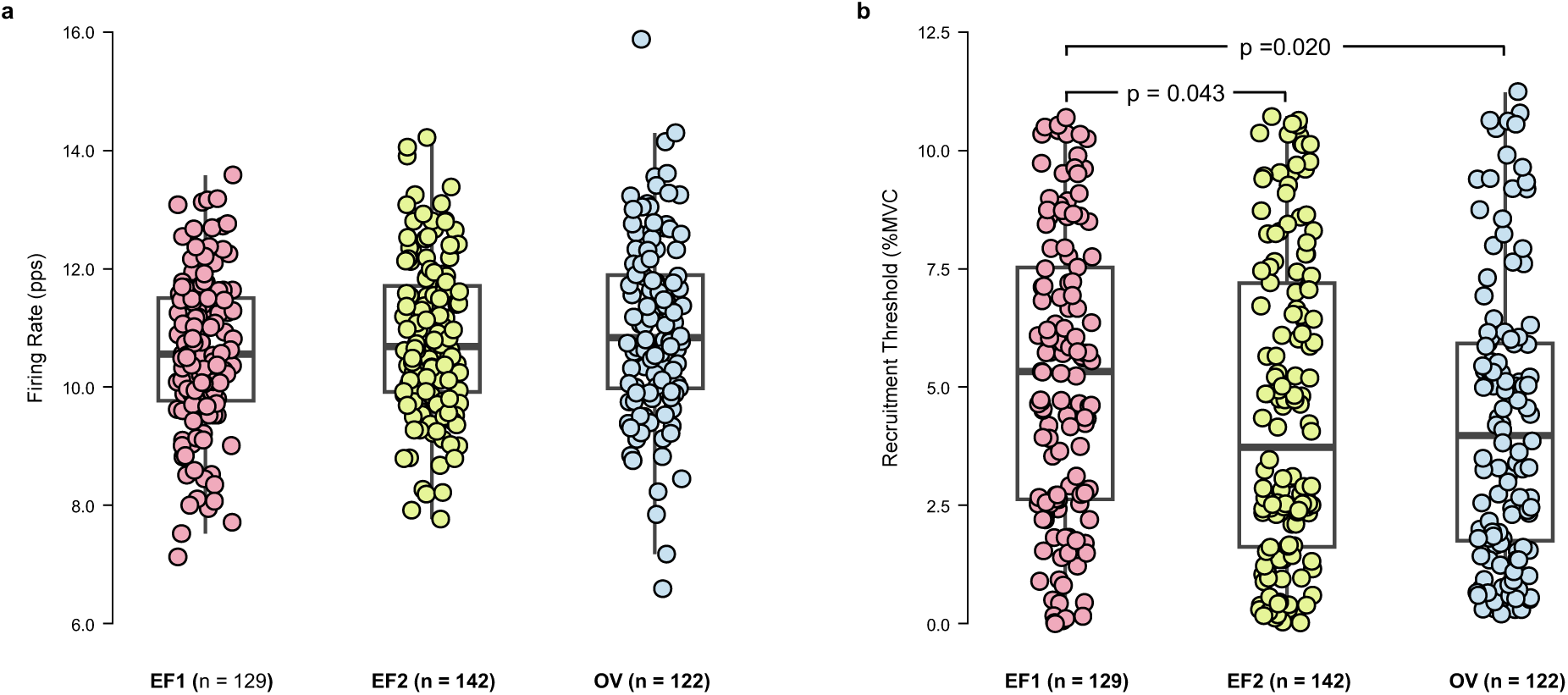
Effect of acute increase in estradiol concentration on MUs characteristics without tracking. The **a**, mean firing rates remained constant across testing visits: 2 visits at early follicular (EF1 and EF2) and 1 visit at peri-ovulatory (OV). **b**, The recruitment threshold at EF1 is significantly higher than EF2 (p = 0.043) and OV (0.020). Data is shown on the original outcome scale. Statistical comparison on firing rate was performed on the non-transformed. Statistical comparison on recruitment threshold was performed on square-root-transformed model scale.

These results highlight the practical use of the algorithm to reveal individual MU characteristic patterns that may be obscured if population-level analysis is used.

## DISCUSSION

Although previous studies have attempted to track individual MU across testing visits, current solutions remain limited by the absence of processing pipeline that automatically identifies the change of grid positioning, insufficient consideration of changes in spatiotemporal characteristics of MUAPs due to relative positioning of the detection electrodes, and constraints in rigorous accuracy validation. These limitations hinder precise classification, reduce the number of reliably tracked MUs, and preclude direct performance comparisons with the proposed algorithm. To address these challenges, we developed a robust technique to accurately track individual MU across two testing visits.

Our experimental results show that the proposed algorithm could, on average, accurately track 63% and 75% of the identified MUs in the TA and MG across testing visits, respectively, while maintaining sensitivity and precision ≥ 70 %, thereby substantially outperforming the SoA method (TA: 7% and MG: 32%). These improvements likely reflect the ability of the proposed algorithm to estimate the translations and rotations of the MUAP spatiotemporal shape between the two testing visits. Compared with the SoA method, the proposed algorithm also achieved higher average PNR, sensitivity, and precision, suggesting greater accuracy for MU firing estimation^18^ and higher validity of MU tracking.

We further demonstrated the practical utility of the proposed algorithm through a case study investigating the effect of acute increase in estradiol concentration on RT and mean FR of MU identified from TA muscle. Motivated by prior evidence that estrogen receptors in skeletal muscle^20,21^ and motoneurons^22^ facilitate estrogen signaling, we expected changes in estradiol levels to be accompanied by changes in MU characteristics. The proposed algorithm enabled tracking of 56% of the identified MUs across EF1, EF2, and OV visits and revealed that RT significantly increased at OV (high estradiol levels) compared with EF2 (low estradiol levels), an association that was not evident when all identified MUs were analyzed. These findings underscore the importance of MU tracking algorithms across testing visits.

Although the underlying mechanism remains to be established, the increase in RT, which represents motoneuron pool’s excitation level,^23^ may reflect changes in muscle fiber size. This interpretation is consistent with prior evidence that post-menopausal women on hormone replacement therapy exhibit greater lean muscle cross-sectional area,^24^ suggesting a potential link between higher estradiol levels and larger muscle fiber size that could increase RT.^25^ By contrast, the decrease in RT detected specifically in EF2 may reflect a delayed effect of low estradiol levels or requirement of prolonged exposure to low circulating estradiol concentrations, warranting further investigation.

Beyond this case study, the proposed algorithm may offer broader methodological advantages. To the best of our knowledge, the proposed algorithm is the first to accurately track a relatively large number of MUs across two or more testing visits, enabling repeated measurement analysis and facilitating paired MU comparisons between testing visits. The use of tracked MUs increased the effect size in statistical analyses, suggesting that smaller sample sizes may be sufficient to detect changes in MU characteristics across multiple testing visits and thereby reduce experiment cost. In addition, the integration of automatic estimation of electrode grid displacement and compensation of MUAP shapes reduces the need and time required for manual editing of motor unit identification results and enable analysis of MU firing estimation accuracy.

These advantages may be relevant to clinical settings, where the proposed algorithm could improve neuromuscular assessments by enhancing the reliability and efficiency of clinical trials. They may also be important in ALS disease, where current clinical assessments remain inadequate.^26^ Although HDsEMG utilization has improved the reproducibility and accuracy of MU number estimation, a precise metric of neuromuscular deterioration, over traditional techniques,^27^ the inability to reliably track the same population of MUs across repeated assessments remains a key limitation, restricting longitudinal comparisons and diminishing sensitivity of biomarker-based evaluations. The proposed algorithm addresses this limitation through consistent MU tracking across clinical visits. In doing so, it may reduce variability and enhance the precision of neuromuscular biomarkers, support robust monitoring of disease progression and therapeutic efficacy, and inform the optimization of patient stratification in ALS and other neuromuscular disorders.

Although these advantages may support broader applications of the proposed algorithm, several findings and limitations warrant further consideration. One unanticipated result was that not all the MUs can be tracked across testing visits. This may be explained by different range of RT between visits, or by the PNR value that is below 28 dB and hence not included in the analysis; however, these possibilities require further examination. Second, the proposed algorithm did not consider the change in load-sharing strategy between agonist muscles and potential differences in muscle excitation across testing visits. Third, although the proposed algorithm employs strict criteria for selecting a pair of MUs to estimate HDsEMG grid displacement, the firing occurrences of MUs with similar spatiotemporal shapes (merging of MUs) in an MU pulse train may still occur. Finally, further validation is needed under conditions that may alter the shape of the action potential, such as fatiguing contraction,^28^ and to assess the sensitivity of the proposed algorithm to electrode grid configuration (inter electrode distance and number of electrodes) and the amount of displacement. Despite these limitations, the proposed algorithm provides a robust tool for MU tracking across testing visits.

## METHODS

### Research participants

We recorded HDsEMG signals from healthy neurologically intact participants. All the participants were engaged in leisure to moderate physical activity and had no history of lower limb injuries or neurological disorders. Each participant provided written informed consent approved by the Institutional Review Boards of the University of Texas Southwestern Medical Center (STU2018-0192 and STU2022-0186), and the experimental procedure was conducted following the Declaration of Helsinki. All testing visits were performed at the same time of the day chosen by the participant to minimize any influence of circadian variance. They were instructed to abstain from caffeinated and alcoholic beverages and refrain from strenuous exercise 24 hours before the testing visit.

### Dataset for performance evaluation

We recorded HDsEMG signals from the TA muscles of 14 female participants (25.9 ± 3.7 yr., BMI 24.1 ± 3.3 kg/m^2^, mean ± SD) and from the MG muscles of 11 female participants (26.1 ± 3.0 yr., BMI 25.7 ± 4.5 kg/m^2^, mean ± SD), in two testing visits separated by 12 – 16 days. HDsEMG signals from the TA and MG were recorded while the participants performing 10% MVC isometric dorsiflexion at and plantar flexion, respectively.

### Dataset for case study

We recorded HDsEMG signals from the TA muscles of nine healthy eumenorrheic female participants (age: 27.4 ± 5.2 years.; BMI: 23.0 ± 2.3 kg/m^2^) while performing isometric dorsiflexion at 10% MVC. To capture changes in estradiol concentration while progesterone remained low, participants completed two testing visits during the early follicular phase (EF1 and EF2) and one during the peri-ovulatory phase (OV), with OV timing determined using urinary ovulation prediction kits. The order of the visits was randomized. During each testing visit, blood was drawn from the antecubital area and analyzed for serum levels of estradiol and progesterone.

After identifying MUs in each visit, we applied the proposed algorithm to identify MUs that were present in EF1 and EF2 visits, as well as in EF2 and OV. MUs that present across all testing visits were identified by intersecting MUs from the EF1-EF2 and EF2-OV matched sets. Subsequently we calculated the mean FR and RT from each MU. The mean FR was determined from the most stable 5-second segment during the plateau phase of the torque trajectory. The RT was assessed based on the torque production normalized to the participant’s MVC at each testing visit and was calculated by averaging the normalized torque over 4 samples beginning at the first firing time of the motor unit.

### Experimental setup and electrode placement

Monopolar HDsEMG signals were collected using a 64-channel electrodes grid positioned over muscle belly while participants were placed in the pronate position on a padded examination table with ankle joint secured in a boot (Fig. 2a). The boot was affixed to a load cell with six degrees of freedom (JR3®, CA, United States) such that the ankle joint was aligned with the rotational axis of the load cell. Surface EMG signals were recorded from MG and TA muscles using a 64-channel HDsEMG grid (13 × 5 electrode grid, 8-mm inter-electrode distance, ELSCH064NM2, OT Bioelettronica, Torino, Italy). The HDsEMG grid was fixed with a disposable bi-adhesive foam layer (Spes Medica, Battipaglia, Italy) with holes filled with conductive paste (Ten20, Weaver and Company, CO, USA). During the placement of the HDsEMG grid, care was taken to ensure that the columns were parallel to the proximal-distal muscle axis. Before electrode placement, the skin was cleaned with alcohol, shaved, and lightly rubbed using a skin abrasive paste (Nuprep®, Weaver and Company, CO, United States), and cleansed with water to remove any residue. The HDsEMG grid site was measured and marked with a permanent marker to ensure electrode placement at subsequent testing visits. Participants were encouraged not to erase the marker location.

HDsEMG signals were recorded in a monopolar configuration, digitized at 2,000 samples/s using a wireless EMG data acquisition system (Sessantaquattro, OT Bioelettronica, Torino, Italy), and acquired using OT Biolab+ software (OT Bioelettronica, Torino, Italy).

The load cell data were sampled at 2,000 samples/s using a Micro1401 Data Acquisition Unit (Cambridge Electronic Design, United Kingdom) and acquired using Spike2 software (version 7.19, Cambridge Electronic Design).

### Isometric dorsiflexion and plantarflexion

Participants were asked to perform isometric dorsiflexion and plantarflexion contractions following a torque trajectory displayed on a monitor (Fig. 2a). The trajectory was trapezoidal: torque increased linearly within 5 seconds, was sustained at 10% MVC, and decreased linearly within 5 seconds. Torque was sustained for 10 seconds during the dorsiflexion task and for 30 seconds during the plantarflexion task. The MVC torque was measured three times and was defined as the highest value of the three trials. At least two minutes of rest were provided between the two subsequent tasks.

### Decomposition of high-density electromyography

We employed a Convolution Kernel Compensation algorithm (CKC) to identify MUs from HDsEMG signals. The mathematical approach of the CKC algorithm is explained in detail elsewhere^18,29^ and summarized in Supplementary Note 1. The resulting firing instances were visually inspected and segmented by an experienced expert operator. Specifically, for each MU, the operator selected the portions of the pulse train that were distinguishable from the background pulse train noise.^30^ The quality of the identified MU firing was then assessed using PNR.^18^ MUs with a PNR < 28 dB were excluded from further analysis.

Subsequently, the MUAP spatiotemporal shapes are calculated using the spike-triggered averaging method, which utilizes the information of the identified MU firings obtained from the decomposition,^31^ and can be written in matrix form as:

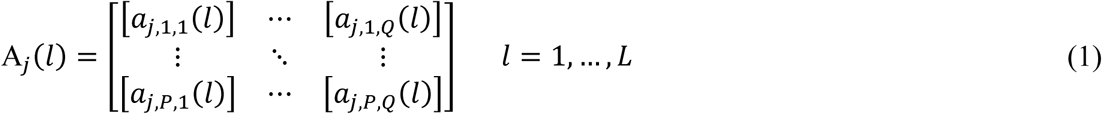

where *a_j_*_,*p*,*q*_(*l*) is the MUAP of the *j*-th MU as recorded by an electrode located at *p*-th row and *q*-th column at the time sample *l*, A*_j_*(*l*) is the MUAP matrix.

### Synthetic displacement of electrodes grid

To benchmark the proposed algorithm under conditions with known ground truth, we simulated controlled HDsEMG grid displacement. Specifically, each HDsEMG recording was synthetically displaced in vertical directions implemented by duplicating the MUAP spatiotemporal shape and removing 1 row from each MUAP matrix.

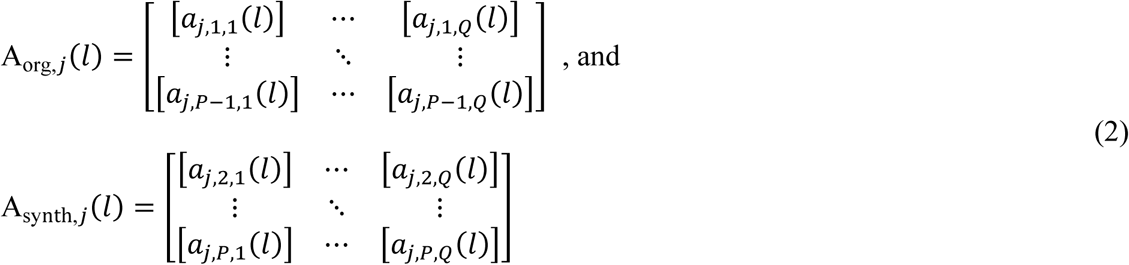

This action is equivalent to removing the HDsEMG channels corresponding to the first and the last row of electrodes, creating a synthetic dataset that contains the same MU captured with HDsEMG grid from its original position and a vertically shifted position. The proposed MU tracking algorithm was then applied to the synthetically generated HDsEMG data.

### MU tracking algorithm

The workflow of the proposed MU tracking algorithm is summarized in Fig. 1a. Briefly, each MUAP matrix was considered as a series of images with spatial resolution that satisfy the Nyquist criterion.^32^ After increasing the spatial resolution of the MUAP, intensity-based image registration with rigid transformation was used to estimate the affine transformation of the MUAP images (Fig. 1c) which reflects the displacement of the HDsEMG electrode grid. In this study we performed this step twice. First, the initial estimation was obtained using MUAP with the largest temporal peak-to-peak in each visit (Fig. 1d). The estimated displacement was then used to transform all MU identified in Visit 1. Subsequently, the similarity of the MUAP spatiotemporal shapes was evaluated and initial list of matched MUs between visits was determined (Fig. 1e). Second, using these initial matches, the grid displacement was re-estimated and averaged across MU (Fig. 1f).

Then the refined estimation of grid displacement was used to transform all MUAP spatiotemporal shape of Visit 1. Following MU filters calculation from the transformed MUAP spatiotemporal shape, MU pulse trains were extracted from HDsEMG signals in Visit 2 (Fig. 1g) as described in Supplementary Note 1. Finally, to determine tracked MUs, the resulting pulse trains were compared with the pulse trains in Visit 2 obtained from independent decomposition of HDsEMG signals in Visit 2 and performance metric (PNR, sensitivity, and precision) was calculated after automatic segmentation^29^ of the pulse trains as described in Supplementary Note 2. Accurately tracked MU was defined as MU with sensitivity and precision that satisfy a threshold. In this study we set the threshold at 70%.

Further details and pseudocode are provided in Supplementary Note 3.

### Performance evaluation

The performance of the proposed algorithm was compared to the SoA method by evaluating the PNR, sensitivity, and precision of the MU firing identification. In summary, the SoA method extracted the MU pulse trains in subsequent HDsEMG signals by applying the MUAP filters obtained from identified MU in Visit 1 directly to HDsEMG signals in Visit 2 (Supplementary Note 1).^17^ Following automatic segmentation^29^ of the pulse train, the PNR values, precision, and sensitivity were calculated (Supplementary Note 2).

We compared the PNR, precision, and sensitivity of MU filters that yielded maximum sensitivity and precision ≥ 70% across the two methods. Additionally, we also assessed the number of tracked MU obtained from each method by counting the MUs with sensitivity ≥ 70%.

### Processing time analysis

We developed the algorithm in MATLAB R2022b (The MathWorks Inc., Natick, MA, USA) using the System Identification Toolbox version 10.0, Signal Processing Toolbox version 9.1, and Image Processing Toolbox version 11.6. The processing time was evaluated by executing the algorithm on a desktop processor (AMD Ryzen 5 3600) with 32 GB of RAM and measuring the time-lapse between the initial estimation of electrode grid displacement and the average translations and rotation across MU (Fig. 1d-f). The time-lapse measurement was repeated 100 times, and the median values of the mean time-lapse across participants and muscles of interest were analyzed. To evaluate the distribution of the execution time among algorithm components, we employed *Profiler app* supplied by MATLAB software.

### Statistical analysis

Statistical analyses were performed in R (version 4.5.3, R Foundation for Statistical Computing, Vienna, Austria). Assumption checks were performed for variables related to the base condition, including MVC and PNR of the MU identified by the decomposition algorithm. We assess the normality and sphericity using the Shapiro-Wilk test and Mauchly’s test, respectively. Sphericity was assessed only on the case study of the effect of estradiol acute increase on MU characteristics.

### Statistical analysis on the performance of the proposed algorithm

The MVC and PNR of the identified MU between testing visits were evaluated using paired t-test because the data followed a normal distribution. Sphericity was not applicable because only two repeated levels were compared. The significance was set to p < 0.05.

To compare the performance of the proposed algorithms and SoA method on MU tracking, Bayesian statistical analyses were used for the primary inference because it enabled direct estimation of the uncertainty and posterior probability of performance differences between the proposed algorithm and SoA method by a practically meaningful margin.

Analyses were implemented using the *brms* package (version 2.23.0).^33^ We compared converged candidate models representing alternative random-effect structures using leave-one-out cross-validation (LOO-CV) method.^34^ The details of convergence analysis model and LOO-CV implementation are provided in Supplementary Note 4.

The PNR was assumed to be independently sampled from a lognormal distribution, whereas the sensitivity and precision were assumed to be independently sampled from a zero-one inflated beta distribution. Weakly informative priors were used in the model. Priors for the intercept, subject-level random-effect standard deviation, and beta precision parameter were chosen using prior predictive check to ensure plausible simulated predicted variable value after back-transformation to the original scale (Supplementary Note 5).

A Bayesian statistical analysis was performed for each predicted variable (PNR, sensitivity, or precision). Three models were fitted and compared to obtain model that best reflects the data. Full candidate-model specifications, model-comparison results, convergence diagnostics, and posterior predictive checks are provided in Supplementary Notes 5-11 in accordance with recent guidelines.^35^ To evaluate the performance on tracking MU across synthetic vertical grid displacement on TA muscle, the selected model was specified as:

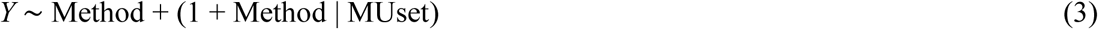

where *Y* was predicted variable (PNR, sensitivity, or precision), Method was set as categorical variable (SoA or proposed algorithm), and MUset denotes the MU set identified for each recording. Similarly, to evaluate the performance on MG muscle across the synthetic displacement, the model above was selected except for analyzing PNR. In analyzing the performance on PNR, following model was selected:

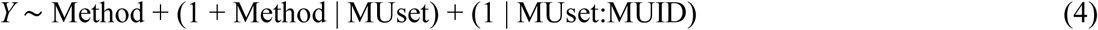

where MUID was the MU identification number. For models describing PNR, we additionally modeled the lognormal-scale parameter as function of Method. For models describing sensitivity and precision, we further model the precision parameter *phi*, zero-one inflation probability *zoi*, and the conditional one-inflation probability *coi* as function of Method.

The posterior inference was based on 4 chains with 10,000 iterations, including 5,000 warm-up iterations. Effects were summarized using posterior medians and 95% credible intervals. Practical equivalence was assessed using region of practical equivalence (ROPE). The consensus in the area of this study has not been established. Therefore, the reference value of ROPE limits for PNR, sensitivity, and precision did not exist. In this study, the ROPE limits for PNR were set to -3 to +3 dB; for sensitivity and precision, the ROPE limits were set to -10% to +10%. The null value of zero difference was rejected if the percentage of the posterior distribution inside the ROPE was less than 2.5% and accepted for practical purposes if the rate of the posterior distribution inside the ROPE was greater than 95%.^36^

Utilizing the Bayesian model specification and inference settings described above, we compared the performance of the proposed algorithm with the SoA method on tracking MU of TA and MG muscles across two testing visits. Similarly, three models were compared, and the model-comparison results, convergence diagnostics, and posterior predictive checks are reported in Supplementary Note 12-17. The selected model for each outcome (PNR, sensitivity, and precision) was described in equation (3). We report the details of the R session info in Supplementary Note 18.

### Statistical analysis on the case study

The data on average PNR of MU firing in three testing sessions were not normally distributed but satisfied the assumption of sphericity, therefore Friedman test was used. A repeated measure ANOVA with Greenhouse-Geisser correction (*rstatix* package^37^) was used to evaluate the estradiol across testing visits, as the data followed a normal distribution but the assumption of sphericity was violated. Post-hoc Holm-adjusted paired t-tests were then used to compare sessions pairwise. To evaluate the progesterone concentrations across testing visits, Friedman test was also used because the data violates both normality and sphericity assumptions. We evaluated the total number of MUs and MVC across testing session using one-way repeated-measures ANOVA because the data satisfied the assumption of normality and sphericity.

To evaluate the effect of acute increase in estradiol concentration on FR and RT of the tracked MUs, a linear mixed effect model (LMM) implemented using *lmerTest* package (version 3.2-1)^38^ was used. The FR and RT were assessed separately using data obtained from accurately tracked MUs across the three testing visits. The tracked MUs are individually labeled by a number *MUID*. The fixed effect set to *Visit*. Model used in the analyses is defined below.

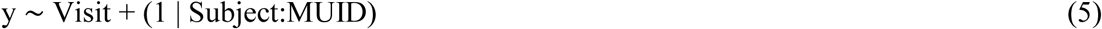

where y is the predicted variable (FR or RT). Details on model diagnostics are provided in Supplementary Note 19.

This analysis was repeated using data from all identified MUs to explore whether untracked MUs would yield a distinct trend across follicular phase. Because MUs were not tracked across testing visits, the observation from EF1, EF2, and OV could not be paired at the MU level. Therefore, this dataset was treated as a repeated cross-sectional comparison of

Visit-level rather than a longitudinal within-MU analysis. Model specification for this dataset was

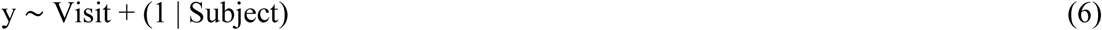

Details on model diagnostics are provided in Supplementary Note 20.

In all models, the covariance parameters were estimated using residual maximum likelihood estimation to obtain an unbiased estimate. The estimated marginal means were used to facilitate post hoc analysis. The significance level of p < 0.05 was used for all statistical analyses. Additionally, the effect size of the statistical analyses on FR and RT of tracked MUs and all identified MU were also compared. The Cohen’s D effect size was calculated using *emmeans* package.^39^

## DATA AVAILABILITY

HDsEMG signals analyzed during the current study are not publicly available due to institutional data-sharing restrictions but are available from the corresponding author upon reasonable requests.

## CODE AVAILABILITY

The code related to the Bayesian statistical analyses and linear mixed effect model are available to editors and reviewers through a private Zenodo record during peer review. Access information has been provided confidentially to the editors. The code will be made publicly available with a persistent DOI upon publication. The algorithm and pipeline of MU tracking described in this study is the subject of a pending patent application. A detailed mathematical description, pseudocode, and a processing workflow are provided in the Methods and Supplementary Information. The source code generated during this study related to the algorithm and pipeline of MU tracking are not publicly available due to proprietary reasons.

## ACKNOWLEDGMENTS

The authors would like to thank all the participants who volunteered in this study. Figure 2a was partially created with illustrae.co.

## AUTHOR CONTRIBUTIONS

A.H. and Y.Y.D. conceived the idea and designed the experiments. S.D.H.S. implemented and performed the experiments. S.D.H.S. and A.H. analyzed the data and experimental results. S.D.H.S., A.H., and Y.Y.D. prepared the manuscript. A.H. and Y.Y.D. jointly supervised the work. S.D.H.S., A.H., and Y.Y.D. approved final version of the manuscript.

