## Supplementary Information for "Beyond snapshots: A dynamic framework for continuous motor unit tracking to enhance neurophysiological assessments"

Supplementary Note 1. Motor unit decomposition and firing pattern extraction 3

Supplementary Note 2. Calculations of pulse to noise ratio, sensitivity, and precision 6

Supplementary Note 3. Implementation 7

Supplementary Note 4. Model convergence and selection 14

Supplementary Note 5. Model implementation, likelihood, priors used in Bayesian analysis on performance comparison between proposed algorithm and SoA 15

Supplementary Note 6. Comparison of firing identification PNR for tracked TA MUs across synthetic grid displacement. 24

Supplementary Note 7. Comparison of firing identification sensitivity in tracked TA MUs across synthetic grid displacement. 32

Supplementary Note 8. Comparison of firing identification precision in tracked TA MUs across synthetic grid displacement. 42

Supplementary Note 9. Comparison of firing identification PNR for tracked MG MUs across synthetic grid displacement. 52

Supplementary Note 10. Comparison of firing identification sensitivity in tracked MG MUs across synthetic grid displacement. 60

Supplementary Note 11. Comparison of firing identification precision in tracked MG MUs across synthetic grid displacement. 70

Supplementary Note 12. Comparison of firing identification PNR for tracked TA MUs across testing visits. 80

Supplementary Note 13. Comparison of firing identification sensitivity in tracked TA MUs across testing visits. 88

Supplementary Note 14. Comparison of firing identification precision in tracked TA MUs across testing visits. 98

Supplementary Note 15. Comparison of firing identification PNR for tracked MG MUs across testing visits. 108
Supplementary Note 16. Comparison of firing identification sensitivity in tracked MG MUs across testing visits. 116
Supplementary Note 17. Comparison of firing identification precision in tracked MG MUs across testing visits. 126
Supplementary Note 18. Bayesian analysis R session info 136
Supplementary Note 19. Linear mixed effect model for the rate and recruitment threshold of tracked MUs across follicular phase 138
Supplementary Note 20. Linear mixed effect model of firing rate and recruitment threshold of all identified motor units across follicular phase 143
Supplementary References 148

### Supplementary Note 1. Motor unit decomposition and firing pattern extraction

#### Decomposition of high-density electromyography

Surface EMG signals at each channel in the High-Density surface EMG (HDsEMG) grid ($y_i(n)$) can be modeled as the summation of all motor unit (MU) activities detected by the corresponding electrode, which can be written as

$$y_i(n) = \sum_{j=1}^{J} \sum_{l=0}^{L-1} a_{i,j}(l) t_j(n-l) \tag{S1}$$

where $a_{i,j}$ is the $j$-th MU action potential (MUAP) with duration of $L$ samples recorded by the $i$-th electrode of the HDsEMG grid, $n$ is time sample, and pulse train $t_j(n)$ indicates the time instances of the MUAP occurrences. The MU pulse train is defined as

$$t_j(n) = \sum_{k} \delta\left(n - \tau_j(k)\right) \tag{S2}$$

where $\delta$ denotes the unit-sample pulse and $\tau_j(k)$ denotes the time instances of the $k$-th MU firing.

The CKC algorithm estimates MU firing instances by inverting the EMG mixing model. It generates MU pulse train for each MU.  The MUAP spatiotemporal shapes ($a_{i,j}(l)$) are calculated using the spike-triggered averaging method, which utilizes the information of the identified MU firings.[1]

Assume that the electrodes on the HDsEMG grid are arranged in $P$ rows and $Q$ columns. For each motor unit $j$, the MUAP spatiotemporal shape can then be written in matrix form as follows:

$$\mathrm{A}_j(l) = \begin{bmatrix} [a_{j,1,1}(l)] & \cdots & [a_{j,1,Q}(l)] \\ \vdots & \ddots & \vdots \\ [a_{j,P,1}(l)] & \cdots & [a_{j,P,Q}(l)] \end{bmatrix} \quad l = 1, \ldots, L \tag{S3}$$

where $a_{j,p,q}(l)$ is the MUAP of the $j$-th MU as recorded by an electrode located at $p$-th row and $q$-th column at the time sample $l$. The matrix $\mathrm{A}_j(l)$ will be defined herein as the MUAP matrix.

**Extracting MU firing pattern using MUAP filters**

Once the MUAP spatiotemporal shapes are identified ($\mathrm{A}_j(l)$), the firing pattern of each MU in the subsequent recording can be extracted by applying the corresponding MU filter ($\mathrm{G}_j$) on the HDsEMG signals. To improve the filter efficiency, both the set of HDsEMG channels and the MUAP matrix $\mathrm{A}_j(l)$ are extended by adding delayed repetitions of recorded HDsEMG channels and MUAP estimates.[2] In this study, we added 14 delayed repetitions of each HDsEMG channel and MUAP. This resulted in extended HDsEMG channels Y and extended MUAP matrix $\mathrm{B}_j$. The MU filter is constructed by multiplying the transpose of $\mathrm{B}_j$ matrix by the inverse of the correlation matrix of the extended HDsEMG signals (${\mathrm{C}_Y}^{-1}$).

$$\mathrm{G}_j = {\mathrm{B}_j}^T {\mathrm{C}_Y}^{-1} \qquad \text{(S4)}$$

The pulse train of each MU is then calculated by multiplying the MU filter by the extended HDsEMG signals (Y).

$$t'_j(n) = \mathrm{G}_j \mathrm{Y}(n) \qquad \text{(S5)}$$

**Automatic MU filter reinforcement**

The MUAP spatiotemporal shapes $a_{i,j}(l)$ and consequently matrix $\mathrm{A}_j(l)$ are calculated by spike-triggered averaging of HDsEMG signals. Thus, they depend on the segmentation of the MU pulses in $t_j(n)$ into MU firings. Improving the segmentation of MU firings enhances the quality of $a_{i,j}(l)$ and $\mathrm{A}_j(l)$ estimation and, thus, the quality of MU filter $\mathrm{G}_j$. In this study, we used an automatic threshold-based MU pulse segmentation technique. First, pulses of $t_j(n)$ in equation (S2) were normalized by their maximal value. Afterward, the segmentation threshold was decremented from 0.9 to 0.05 in steps of 0.05. After each decrement, MU pulses exciding the threshold were segmented into MU firings. The coefficient of variation (CoV) of the interspike interval of segmented MU firings was calculated for each segmentation threshold, along with the median MU firing rate (MU_FR). The cost of MU firing segmentation included the CoV, and the MU segmentations with MU_FR < 5 Hz or MU_FR > 35 Hz were penalized by adding the cost of 100. The segmentation threshold with the minimal cost value was then selected and used for the final segmentation of MU firings. Afterward, the MUAP spatiotemporal shapes $a_{i,j}(l)$, the matrix $\mathrm{A}_j(l)$ and the MU filter $\mathrm{G}_j$ were recalculated and reapplied to the extended HDsEMG signals Y (equation (S5)).

**Manual inspection and editing of decomposition results**

The obtained MU pulse trains were visually inspected and segmented by an expert operator. Specifically, for each MU, the operator selected the portions of the pulse train that were distinguishable from the background pulse train noise.[3] Once all pulse trains were selected, the spatiotemporal shape of the MUAP (and thus the MU filter) was recalculated using the segmented MU firing instances. Afterwards, the pulse train of the MU was recalculated. The segmentation process was repeated until the optimum decomposition quality of the identified MU was reached.

**Supplementary Note 2. Calculations of pulse to noise ratio, sensitivity, and precision**

The identified motor unit's decomposition quality was quantified using the pulse-to-noise ratio (PNR). PNR indicates the level of noise in the MU pulse train[4] and is defined as follows:

$$\mathrm{PNR}\left(t_j(n)\right) = 20 \times \log\left(\frac{E\left(t_j^2(n)\big|_{t_j(n)=1}\right)}{E\left(t_j^2(n)\big|_{t_j(n)=0}\right)}\right) \tag{S6}$$

where $E$ represents the mathematical expectation, $t_j^2(n)\big|_{t_j(n)=1}$ denotes the segmented MU pulses denoting MU firing and $t_j^2(t)\big|_{t_j(t)=0}$ denotes the baseline noise at the $n$-th time sample. Subsequently, MUs with a PNR < 28 dB were excluded from further analysis.[4]

Assume two HDsEMG signals were decomposed independently. MU filter transfer from the first to the second HDsEMG signals was defined as a process of applying MU filters calculated from the identified MU in the first HDsEMG signals to the second HDsEMG signals (Supplementary equations (S4) and (S5)), and segmentation of the MU firings was subsequently repeated. For MUs that were identified in both HDsEMG signals, the similarity between the firing patterns of MUs identified from the independent decomposition of the second HDsEMG signals and MUs filter transfer from the first HDsEMG signals was assessed by calculating the precision and sensitivity of MU firing identification (precision and sensitivity, respectively)[5]

$$\mathrm{precision}_j = \frac{\mathrm{TP}}{\mathrm{TP+FP}} \quad , \quad \mathrm{sensitivity}_j = \frac{\mathrm{TP}}{\mathrm{TP+FN}} \tag{S7}$$

where TP, FP, and FN denote the number of true positive, false positive, and false negative firings, respectively. The tolerance limit of the MU firing match was set to 1 ms, and the results of the subsequent HDsEMG signals independent decompositions were used as ground truth reference.

### Supplementary Note 3. Implementation

This Supplementary Note provides additional details for the algorithm depicted in Fig. 1 and Fig. 2. Vectors are denoted by bold letters.

The HDsEMG signals recorded at two testing visits (Visit 1 and Visit 2) are denoted as $y1(n)$ and $y2(n)$, respectively. Each HDsEMG signal was independently decomposed in a monopolar configuration into its constituent MUs pulse trains using the CKC algorithm (Fig.1b).[2] The spatiotemporal shape of the MUAP was extracted from the single differential HDsEMG signal computed along the longitudinal axis of the muscle fibers. In this study, the single differential HDsEMG signal was computed along the column of electrode grid. MUAP matrices for datasets $y1(n)$ and $y2(n)$ are denoted as $\mathrm{A1}_r(l)$ and $\mathrm{A2}_s(l)$ where $r = 1, \ldots, R$ and $s = 1, \ldots, S$ denotes the identified MUs in each dataset, respectively.

The first step of the proposed algorithm was to calculate an initial estimate of the electrode grid displacement between visits (Fig. 1d). To begin this process, the maximum temporal peak-to-peak value across channels for each MUAP matrix was calculated. The MUs in each testing visit were sorted in descending order based on their maximum temporal peak-to-peak value across channels, resulting in new MUs orders $r_{\mathrm{srt}}$ and $s_{\mathrm{srt}}$. The initial electrode grid displacement was then estimated starting with MUs with the largest temporal peak-to-peak MUAPs across channels in datasets $y1(n)$ and $y2(n)$.

Each MUAP matrix was considered as a series of images. Specifically, each electrode of the grid may be conceived as a pixel in an image with coordinates $(p, q)$ given by the rows and columns in the HDsEMG grid of electrodes with a magnitude that evolves across time sample $l$ (Fig. 2b). The temporal root mean square (RMS) envelope of each MUAP image in the datasets was calculated and denoted as $U1$ and $U2$, respectively.

$$U1_{r_{\mathrm{srt}}}(l) = \sqrt{\frac{1}{PQ}\sum_{p=1}^{P}\sum_{q=1}^{Q}\mathrm{A1}_{r_{\mathrm{srt}},p,q}{}^{2}(l)} \quad \text{and} \quad U2_{s_{\mathrm{srt}}}(l) = \sqrt{\frac{1}{PQ}\sum_{p=1}^{P}\sum_{q=1}^{Q}\mathrm{A2}_{s_{\mathrm{srt}},p,q}{}^{2}(l)} \qquad \text{(S8)}$$

The temporal profiles of the MUAP images' RMS envelopes ($U1_{r_{\mathrm{srt}}}(l)$ and $U2_{s_{\mathrm{srt}}}(l)$) were observed to vary in time. Therefore, to align them, dynamic time warping (DTW)[6] was employed and DTW-aligned time samples are denoted as $l_{\mathrm{DTW}}$. DTW aligns the time series between two out-of-phase datasets by calculating their minimum cost path. In this study, DTW-aligned time samples ($l_{\mathrm{DTW}}$) for which the RMS of the MUAP images in both datasets ($U1_{r_{\mathrm{srt}}}(l)$ and $U2_{s_{\mathrm{srt}}}(l)$) reached at least 30% of their respective maximum value ($l_{\mathrm{DTW},\geq 30\mathrm{p}}$) were used in further processing and were also used to temporarily align MUAP images from both visits.

The inter-electrode distance of the HDsEMG grid was assumed to satisfy the Nyquist criterion for HDsEMG spatial sampling;[7] therefore, the continuous spatial distribution of the MUAP can be obtained by spatially interpolating the DTW-aligned MUAP images using a two-dimensional sinc function.[8] In this study, the MUAP images were interpolated to obtain a new spatial resolution of 0.8 mm (denoted as $\mathrm{A1}_{\mathrm{HiRes},r_{\mathrm{srt}}}(l_{\mathrm{DTW},\geq 30\mathrm{p}})$ and $\mathrm{A2}_{\mathrm{HiRes},s_{\mathrm{srt}}}(l_{\mathrm{DTW},\geq 30\mathrm{p}})$ for the first and the second testing visit, respectively). Subsequently, intensity-based image registration with rigid transformation supplied within MATLAB Image Processing Toolbox was performed on the interpolated MUAP images ($\mathrm{A1}_{\mathrm{HiRes},r_{\mathrm{srt}}}(l_{\mathrm{DTW},\geq 30\mathrm{p}})$ and $\mathrm{A2}_{\mathrm{HiRes},s_{\mathrm{srt}}}(l_{\mathrm{DTW},\geq 30\mathrm{p}})$) to estimate the affine transformation (vertical and horizontal translations and rotation) with interpolated MUAP image $\mathrm{A2}_{\mathrm{HiRes},s_{\mathrm{srt}}}(l_{\mathrm{DTW},\geq 30\mathrm{p}})$ as reference. The optimizing method of the image registration was set to regular step gradient descent, and the quantitative measure of similarity was set to mean square error. The spatially interpolated MUAP images $\mathrm{A1}_{\mathrm{HiRes},r_{\mathrm{srt}}}(l_{\mathrm{DTW},\geq 30\mathrm{p}})$ were then transformed using the estimated affine transformation and resampled to restore the spatial resolution to 8 mm ($\mathrm{A1}_{\mathrm{resampled},r_{\mathrm{srt}}}\left(l_{\mathrm{DTW},\geq 30\mathrm{p}}\right)$).

The normalized root mean square error (NRMSE) between MUAP images $\mathrm{A1}_{\mathrm{resampled.},r_{\mathrm{srt}}}\left(l_{\mathrm{DTW},\geq 30\mathrm{p}}\right)$ and $\mathrm{A2}_{s_{\mathrm{srt}}}\left(l_{\mathrm{DTW},\geq 30\mathrm{p}}\right)$ was calculated as follows.

$$X_{r_{\mathrm{srt}},s_{\mathrm{srt}}}\left(l_{\mathrm{DTW},\geq 30\mathrm{p}}\right)=\frac{\left\|\mathrm{A2}_{s_{\mathrm{srt}}}\left(l_{\mathrm{DTW},\geq 30\mathrm{p}}\right)-\mathrm{A1}_{\mathrm{resampled},r_{\mathrm{srt}}}\left(l_{\mathrm{DTW},\geq 30\mathrm{p}}\right)\right\|}{\left\|\mathrm{A2}_{s_{\mathrm{srt}}}\left(l_{\mathrm{DTW},\geq 30\mathrm{p}}\right)-\mathrm{mean}\left(\mathrm{A2}_{s_{\mathrm{srt}}}\left(l_{\mathrm{DTW},\geq 30\mathrm{p}}\right)\right)\right\|} \tag{S9}$$

where || indicates the 2-norm. If all the NRMSE values across the DTW-aligned time samples $\left(l_{\mathrm{DTW},\geq 30\mathrm{p}}\right)$ were greater than 30%, the initial electrode grid displacement estimation was repeated using the subsequent MU pair. The affine transformations that yielded an NRSME of less than 30% were then averaged across the time sample $l$ and were considered as the initial estimate of the electrode grid displacement ($\mathrm{GD_{init}}$, Fig. 1d and Fig. 2c).

The second step was to find matched MUs by evaluating the MUAP shape and amplitude (Fig. 1e). All MUAPs in Visit 1 were forward transformed using $\mathrm{GD_{init}}$, yielding a transformed MUAP spatiotemporal shape $\mathrm{A1}_{\mathrm{GD_{init}},r}(l)$. The MUAP spatiotemporal shapes ($\mathrm{A1}_{\mathrm{GD_{init}},r}(l)$ and $\mathrm{A2}_s(l)$) were vectorized by row-wise concatenating the MUAP profile of each non-empty HDsEMG channel (Fig. 2d), yielding $\mathbf{A1}_{\mathrm{GD_{init}},r}$ and $\mathbf{A2}_s$. To evaluate the similarity of the MUAP shape, the cross-correlation coefficient ($\rho_{r,s}$) between $\mathbf{A1}_{\mathrm{GD_{init}},r}$ and $\mathbf{A2}_s$ were evaluated for all possible MU pairs.

$$\rho_{r,s}=\frac{E\left\{\mathbf{A1}_{\mathrm{GD_{init}},r}\ \mathbf{A2}_s\right\}-m_{\mathbf{A1}_{\mathrm{D_{init}},r}}\,m_{\mathbf{A2}_s}}{\sigma_{\mathbf{A1}_{\mathrm{GD_{init}},r}}\sigma_{\mathbf{A2}_{\mathrm{s}}}} \tag{S10}$$

where $E$ denotes the mathematical expectation, $m_{\mathbf{A1}_{\mathrm{D_{init}},r}}$ and $m_{\mathbf{A2}_s}$ are the average values of $\mathbf{A1}_{\mathrm{GD_{init}},r}$ and $\mathbf{A2}_s$ respectively, $\sigma_{\mathbf{A1}_{\mathrm{GD_{init}},r}}$ and $\sigma_{\mathbf{A2}_{\mathrm{s}}}$ are the standard deviations. MU pairs with $\rho_{r,s}$ greater than 0.70 were selected as the initial matched MU pairs.

The MUAP amplitude detected by each electrode depends on the distance between the MU fibers and the electrode.[9] Hence, a pair of MUs with similar shapes but significantly different amplitudes are unlikely to be generated by the same source. To evaluate the differences in the amplitude, the time-wise RMS envelopes were recalculated excluding the missing and discarded electrodes data resulting from the initial affine transformation.

$$W1_r(l) = \sqrt{\frac{1}{P_{\text{adj}}Q_{\text{adj}}}\sum_{p=1}^{P_{\text{adj}}}\sum_{q=1}^{Q_{\text{adj}}} \text{A1}_{\text{GD}_{\text{init}},r,p,q}{}^2(l)} \quad \text{and} \quad W2_s(l) = \sqrt{\frac{1}{P_{\text{adj}}Q_{\text{adj}}}\sum_{p=1}^{P_{\text{adj}}}\sum_{q=1}^{Q_{\text{adj}}} \text{A2}_{s,p,q}{}^2(l)} \tag{S11}$$

where $P_{\text{adj}}$ and $Q_{\text{adj}}$ were the total number of rows and columns of the MUAP spatiotemporal shape with missing and discarded electrodes excluded. Afterward, following DTW time sample alignment, the normalized difference of the RMSE values were calculated.

$$Z_{r,s} = \frac{\| W2_s(l_{\text{DTW}}) - W1_r(l_{\text{DTW}}) \|}{\left\| W2_s(l_{\text{DTW}}) - \text{mean}\big(W2_s(l_{\text{DTW}})\big) \right\|} \tag{S12}$$

where || indicates the 2-norm and $l_{\text{DTW}}$ is DTW aligned time sample. MU pairs with $Z$ values greater than 0.3 were considered false positive pairs and were discarded. Subsequently, MUs in Visit 1 with multiple possible matches with MUs in Visit 2 were identified. For each MU with multiple possible matches, MU pair with highest $\rho_{r,s}$ was considered as a true pair. If there were more than one MU pair with equaly high $\rho_{r,s}$, the $Z_{r,s}$ values of these MU pairs were compared and MU pair with lowest $Z_{r,s}$ was used in further analysis.

The third step was to refine the candidates of the matched MU by evaluating the affine transformation of each MU pair. For each MU pair, the transformed MUAP images of Visit 1 ($\text{A1}_{\text{GD}_{\text{init}},r}(l)$) and Visit 2 ($\text{A2}_s(l)$) were aligned in time using DTW algorithms using the temporal RMS envelope (equation 11). The MUAP images were then interpolated using two-dimensional sinc function ($A1_{\mathrm{GDHiRes}.r}(l_{\mathrm{DTW}})$ and $A2_{\mathrm{HiRes}.s}(l_{\mathrm{DTW}})$). Similar to the first step of the proposed algorithm, the affine transformation between $A1_{\mathrm{GDHiRes}.r}(l_{\mathrm{DTW}})$ and $A2_{\mathrm{HiRes}.s}(l_{\mathrm{DTW}})$ was estimated using intensity-based registration with $A2_{\mathrm{HiRes}.s}(l_{\mathrm{DTW}})$ used as reference (Fig. 1c). The estimated translation and rotation values were averaged across all the MUs in Visit 1 and defined as the refined compensation of the HDsEMG grid displacement (Fig. 1f, $GD_{\mathrm{refined}}$). Finally, $GD_{\mathrm{refined}}$ was applied to all the MU filters in Visit 1, MU filters were calculated using Supplementary equation (S4), and the algorithm's performance was evaluated. The overall algorithm is presented in pseudocode below.

---

**Algorithm 1. Motor unit tracking across testing visits**

---

1. Calculate the maximum MUAP peak-to-peak value across the electrodes of all the identified MUs in Visit 1 and Visit 2
2. Sort in descending order all the MUs in Visit 1 and Visit 2 based on the maximum MUAP temporal peak-to-peak value across channels, $A1_{r_{\mathrm{srt}}}$ and $A2_{s_{\mathrm{srt}}}$
3. found initial estimation of the grid displacement ($GD_{\mathrm{init}}$) ← FALSE
4. while found $GD_{\mathrm{init}}$ = FALSE do
5.     for all possible pairs between $A1_{r_{\mathrm{srt}}}$ and $A2_{s_{\mathrm{srt}}}$
6.         Calculate the temporal RMS of the MUAP images (Supplementary equation (S8)), $U1_{r_{\mathrm{srt}}}(l)$ and $U2_{s_{\mathrm{srt}}}(l)$
7.         Align time sample using DTW algorithm, $U1_{r_{\mathrm{srt}}}(l_{\mathrm{DTW}})$ and $U2_{s_{\mathrm{srt}}}(l_{\mathrm{DTW}})$
8.         Find DTW time sample where $U1_{r_{\mathrm{srt}}}(l_{\mathrm{DTW}})/\max(U1_{r_{\mathrm{srt}}})$ and $U2_{s_{\mathrm{srt}}}(l_{\mathrm{DTW}})/\max(U2_{s_{\mathrm{srt}}})$ are greater than 30%, $l_{\mathrm{DTW},\geq 30\mathrm{p}}$
9.         for all DTW time samples $l_{\mathrm{DTW},\geq 30\mathrm{p}}$

10. Spatially up-sample of MUAP images, $\mathrm{A1}_{\mathrm{HiRes},r_{\mathrm{srt}}}\left(l_{\mathrm{DTW},\geq 30\mathrm{p}}\right)$ and $\mathrm{A2}_{\mathrm{HiRes},s_{\mathrm{srt}}}\left(l_{\mathrm{DTW},\geq 30\mathrm{p}}\right)$

11. Estimate translation and rotation using intensity-based registration

12. Apply translation and rotation to $\mathrm{A1}_{\mathrm{HiRes},r_{\mathrm{srt}}}\left(l_{\mathrm{DTW},\geq 30\mathrm{p}}\right)$

13. Resample $\mathrm{A1}_{\mathrm{HiRes},r_{\mathrm{srt}}}\left(l_{\mathrm{DTW},\geq 30\mathrm{p}}\right)$ to original spatial resolution, $\mathrm{A1}_{\mathrm{resampled},r_{\mathrm{srt}}}\left(l_{\mathrm{DTW},\geq 30\mathrm{p}}\right)$

14. Calculate the NRMSE between $\mathrm{A1}_{\mathrm{resamp.},r_{\mathrm{srt}}}\left(l_{\mathrm{DTW},\geq 30\mathrm{p}}\right)$ and $\mathrm{A2}_{s_{\mathrm{srt}}}\left(l_{\mathrm{DTW},\geq 30\mathrm{p}}\right)$ (Supplementary equation (S9)), $X_{r_{\mathrm{srt}},s_{\mathrm{srt}}}\left(l_{\mathrm{DTW},\geq 30\mathrm{p}}\right)$

15. if $X_{r_{\mathrm{srt}},s_{\mathrm{srt}}}\left(l_{\mathrm{DTW},\geq 30\mathrm{p}}\right) > 0.3$ then

16. Discard the estimated translation and rotation (Fig. 2c)

17. if $\mathrm{length}\left(X_{r_{\mathrm{srt}},s_{\mathrm{srt}}}\left(l_{\mathrm{DTW},\geq 30\mathrm{p}}\right) \leq 0.3\right) > 0$ then

18. $\mathrm{GD_{init}} \leftarrow$ mean translations and rotation across $l_{\mathrm{DTW},\geq 30\mathrm{p}}$

19. found $\mathrm{GD_{init}}$ = TRUE

20. Apply $\mathrm{GD_{init}}$ to all the MUAP matrices in Visit 1, $\mathrm{A1}_{\mathrm{GD_{init}},r}(l)$

21. for all possible pairs of MUs between Visit 1 and Visit 2

22. Vectorize each MUAP matrix without missing electrodes (Fig. 2d), $\mathrm{A1}_{\mathrm{GD_{init}},r}$ and $\mathrm{A2}_s$

23. Calculate the correlation coefficient of the vectorized MUAPs (Supplementary equation (S10)), $\rho_{r,s}$

24. if $\rho_{r,s} \geq 0.70$ then

25. MU list $\leftarrow$ MU pair

26. for all MU pairs in MU list

27. Calculate the time-wise RMS envelope of the MUAP images without missing/discarded electrode (Supplementary equation (S11)), $W1_r(l)$ and $W2_s(l)$

28. Align time sample using DTW algorithm, $W1_r(l_{\mathrm{DTW}})$ and $W2_s(l_{\mathrm{DTW}})$

29. Calculate the normalized difference of the RMS values between $W1_r(l_{\mathrm{DTW}})$ and $W2_s(l_{\mathrm{DTW}})$ (Supplementary equation (S12)), $Z_{r,s}$

30. if $Z_{r,s} > 0.3$ then

31. remove MU pair from MU list

32. for all MUs in MU list with multiple matches

33. Find the highest correlation coefficient, $\rho_{max}$

34. if $\rho_{r,s} < \rho_{max}$ then

35. remove MU pair from MU list

36. if number of matched MU >1 then

37. Find lowest normalized difference of the RMSE values, $Z_{min}$

38. if $Z_{r,s} > Z_{min}$ then

39. remove MU pair from MU list

40. for all MU pairs in MU list

41. Repeat steps 6 to 18 to obtain an estimation on electrode grid translation and rotation

42. $\mathrm{GD}_{\mathrm{refined}} \leftarrow$ average translation and rotation across MU pairs in MU list

43. Apply $\mathrm{GD}_{\mathrm{refined}}$ to all MUAPs in Visit 1

44. Calculate MU filters (Supplementary equation (S4)) and reassess the MU filter for transfer to HDsEMG signals from Visit 2

---

**Supplementary Note 4. Model convergence and selection**

The convergence of each model's Markov Chain Monte Carlo (MCMC) chains was evaluated using the potential scale reduction factor (PSRF) and effective sample size (ESS) calculated using Gelman and Rubin’s convergence diagnostic and *effectiveSize* function supplied in the *coda* package, respectively.[10,11] Convergence of the MCMC chains is achieved if the PSRF <1.05 and ESS >400.[12,13] In the case where only PSRF satisfies the criteria, the model was not included in further analysis.

The leave-one-out cross-validation (LOO-CV) method[14] was then used to evaluate the goodness of fit of each model. The model with the best goodness of fit was defined as one with the highest Bayesian leave-one-out estimate of the expected log pointwise predictive density. If 0.7 < Pareto $k$ < 1.0 of any observation and the posterior predictive check was consistent with the observed data, the *reloo* function supplied by the *loo* package was used to compute exact cross-validation for these problematic observations. If Pareto $k$ estimated after refit was >1 for at least one observation, the model was not included in further analysis.

### Supplementary Note 5. Model implementation, likelihood, priors used in Bayesian analysis on performance comparison between proposed algorithm and SoA

#### Performance on pulse-to-noise ratio (PNR)

The state-of-the-art (SoA) method calculated MU filters from MUs identified in HDsEMG signal recorded at the first testing visit and applied it directly (without MUAP shapes compensation) to subsequent HDsEMG signals to extract MU characteristics.[5] We compared the performance of our proposed algorithm to SoA approach on the PNR of the motor unit firing identification. The predicted variable was PNR (continuous values) and the predictor variable was Method (SoA or Proposed, categorical variable). Three models were fitted and compared to obtain model that best reflects the data. In all models, for performance comparison of tracked MUs across testing visits, Subject as participant identification was used instead of MUset.

Model 1

The conditional mean of the log-transformed response was modeled as a function of Method with MUset-level random intercept. To allow residual variation to differ between Method, the lognormal-scale parameter was modeled as a function of Method. The model was implemented using the formula bf(PNR ~ Method + (1 | MUset), sigma ~ Method). The complete specification was given by the likelihood and prior distributions below.

Based on visual inspection of the data distribution, PNR was strictly positive and right skewed. Therefore, the PNR observations were assumed to be independently sampled from a lognormal distribution.

$$y_i|\mu_i, \sigma_i \sim \text{LogNormal}(\mu_i, \sigma_i) \quad \text{(S13)}$$

with

$$\mu_i = \alpha_\mu + \mathrm{x}_i{}^T \beta_\mu + b_{s_i} \quad \text{(S14)}$$

and $$\log \sigma_i = \alpha_\sigma + \mathrm{z}_i{}^T \beta_\sigma \tag{S15}$$

Here $y_i$ is PNR for motor unit $i$, $s_i$ denotes the MU set for motor unit $i$, $\mu_i$ is conditional mean of the log-transformed response ($\log(y_i)$), $\sigma_i$ is conditional residual standard deviation of $\log(y_i)$, $\alpha_\mu$ denotes population-level intercept for the mean of the $\log(y_i)$, $\mathrm{x}_i$ encodes the method of calculating MU filter (SoA or proposed algorithm), $\beta_\mu$ are method effects on the mean of $\log(y_i)$, and $b_{s_i}$ is MU set specific random intercept. Additionally, $\alpha_\sigma$ is the population-level intercept for the log residual standard deviation, $\mathrm{z}_i$ denotes method in the sigma model, and $\beta_\sigma$ are method effects on $\log \sigma_i$.

Weakly informative priors were used in the model and described as follows.

$$\alpha_\mu \sim \mathrm{Normal}(\log(30), 0.25^2) \tag{S16}$$

$$\beta_{\mu,k} \sim \mathrm{Normal}(0, 0.20^2),\ k = 1, \dots, K_\mu \tag{S17}$$

$$b_j | \tau_b \sim \mathrm{Normal}(0, \tau_b{}^2),\ j = 1, \dots, J \tag{S18}$$

$$\tau_b \sim \mathrm{Exponential}(2) \tag{S19}$$

$$\alpha_\sigma \sim \mathrm{Normal}(\log(0.15), 0.35^2) \tag{S20}$$

$$\beta_{\sigma,k} \sim \mathrm{Normal}(0, 0.25^2),\ k = 1, \dots, K_\sigma \tag{S21}$$

Here, $K_\mu$ is number of non-intercept method coefficients in the model for $\mu_i$, $b_j$ is MU set-specific random intercept for MU set $j$, $\tau_b$ is between-MU set standard deviation of the random intercepts $b_j$, $J$ is number of MU set, $K_\sigma$ is number of non-intercept method coefficients in the sigma model.

Priors were specified to be weakly informative on the scale of the observed PNR values. For each analysis, these priors were evaluated using prior predictive simulation before posterior inference to confirm that the priors generated PNR in a plausible range (Supplementary Fig. 1a).

Model 2

In this model we additionally allowed the Method effect to vary by MUset by adding MUset-specific Method slopes. The model was implemented using the formula bf(PNR ~ Method + (1 + Method | MUset), sigma ~ Method). The model specification was similar to Model 1, except

$$\mu_i = \alpha_\mu + \mathbf{x}_i{}^T \beta_\mu + \mathbf{r}_i{}^T \mathbf{b}_{s_i} \tag{S22}$$

where $\mathbf{r}_i$ is group-level design vector for observation $i$.

The likelihood and priors were set according to Model 1, except for $b_j | \tau_b \sim \text{Normal}(0, \tau_b{}^2)$ which was replaced with random intercept and random method effects defined as

$$\mathbf{b}_j \big| \boldsymbol{\tau}_b, \mathbf{R} \sim \text{MultivariateNormal}\big(0, \text{diag}(\boldsymbol{\tau}_b)\ \mathbf{R}\ \text{diag}(\boldsymbol{\tau}_b)\big) \tag{S23}$$

with

$$\mathbf{b}_j = \left(b_{0j}, b_{1j}, \ldots, b_{K_\mu j}\right)^{\mathrm{T}} \tag{S24}$$

where $\mathbf{R}$ contains the correlation between the random intercept and the random method effects.

We additionally assigned an LKJ prior to the correlation matrix among the MUset-specific intercept and Method effects, defined as

$$\mathbf{R} \sim \text{LKJ}(2) \tag{S25}$$

For each analysis, prior predictive simulations confirmed that the priors generated plausible values of PNR (Supplementary Fig. 1b).

Model 3

Compared to Model 2, this model additionally allowed observations from each MUset-MUID combination to have their own random intercept. The model was implemented using the formula bf(PNR ~ Method + (1 + Method | MUset) + (1 | MUset:MUID), sigma ~ Method). Therefore,

$$\mu_i = \alpha_\mu + \mathbf{x}_i{}^T \beta_\mu + \mathbf{r}_i{}^T \mathbf{b}_{s_i} + c_{s_i l_i} \tag{S26}$$

Here $c_{s_i l_i}$ denotes random intercept for the MUset-MUID combination associated with observation $i$.

The likelihood and priors from Model 2 were used in this model because Model 2 included a broad prior for the class-level random-effect standard deviation (class = "sd"). Then, we performed prior predictive simulations for each analysis to confirm that the priors generated PNR in a plausible range (Supplementary Fig. 1c).

**Performance on sensitivity and precision of motor unit firing identification**

We compared the performance of our proposed algorithm to SoA separately for each predicted continuous variable $Y$: sensitivity and precision. The predictor variables were Method (SoA or Proposed, categorical variable). Three models were fitted and compared to obtain model that best reflects the data. In all models, for performance comparison of tracked MUs across testing visits, Subject as participant identification was used instead of MUset.

<u>Model 1</u>

The model used zero-one-inflated beta response distribution that includes Method as a population-level predictor and MUset-level random intercept. To capture Method specific effects, the mean, precision parameter *phi*, zero-one inflation probability *zoi*, and the conditional one-inflation probability *coi* were each modeled as functions of method. The model was implemented using the formula bf(Y ~ Method + (1 | MUset), phi ~ Method, zoi ~ Method, coi ~ Method).

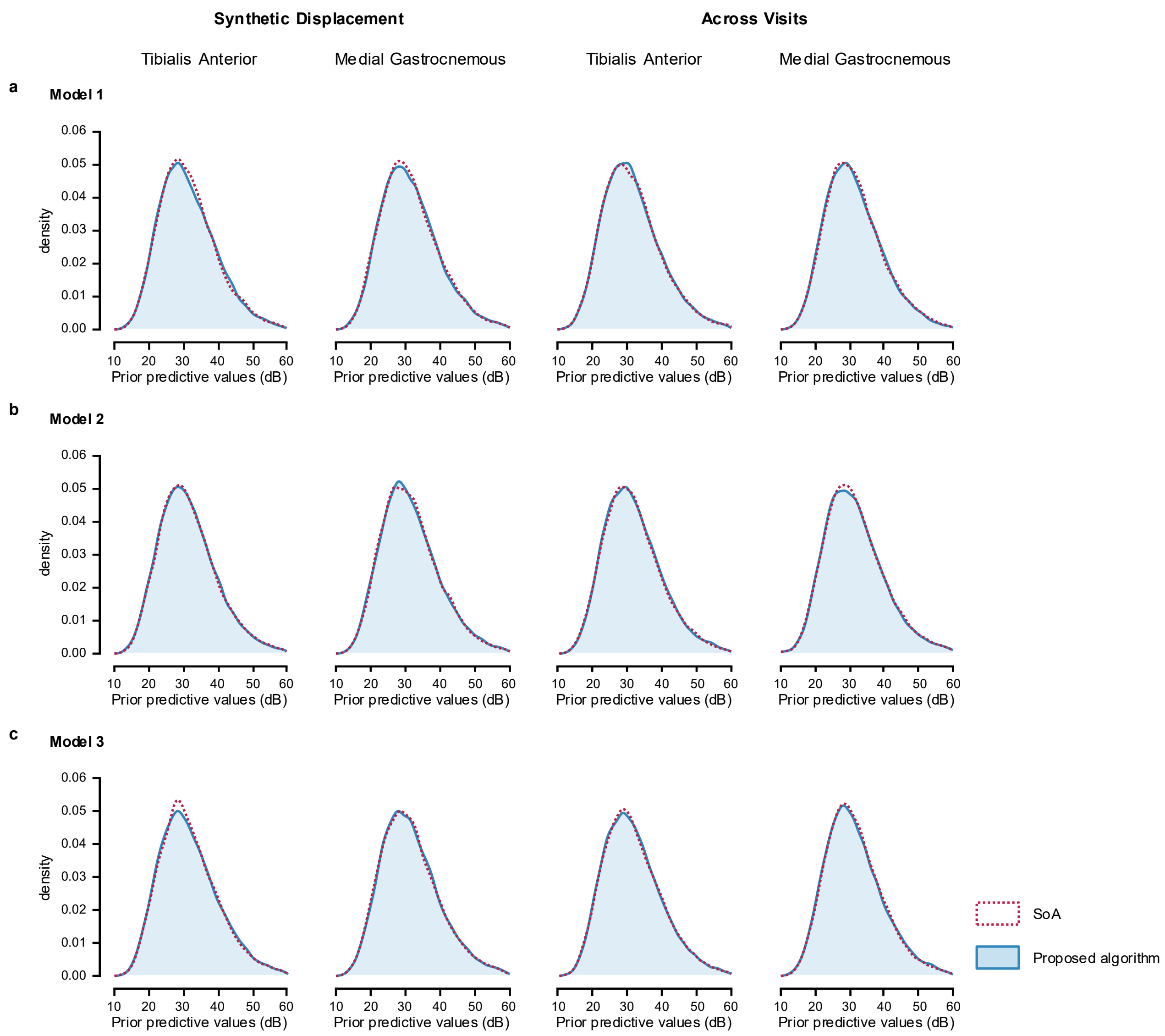


**Supplementary Fig. 1: Prior predictive check for Bayesian comparison on PNR**

Prior predictive checks for the Bayesian comparison of SoA and the proposed algorithm based on PNR of MU firing in tibialis anterior and medial gastrocnemius muscles, obtained by tracking MUs across synthetic displacement of electrode grid and two testing visits. Simulations were performed for **a**, Model 1; **b**, Model 2; and **c**, Model 3.

Based on visual inspection of data distribution, the sensitivity values were assumed to be independently sampled from a zero-one-inflated beta distribution.

$$y_i|\mu_i, \phi_i, \zeta_i, \kappa_i \sim \mathrm{ZOIB}(\mu_i, \phi_i, \zeta_i, \kappa_i) \tag{S27}$$

with

$$\mu_i, \zeta_i, \kappa_i \in (0,1) \tag{S28}$$

and

$$\phi_i > 0 \tag{S29}$$

where

$$\mathrm{ZOIB}(\mu_i, \phi_i, \zeta_i, \kappa_i) = \begin{cases} \zeta_i(1-\kappa_i) & , y_i = 0 \\ \zeta_i\kappa_i & , y_i = 1 \\ (1-\zeta_i)\ \mathrm{Beta}(\mu_i\phi_i, (1-\mu_i)\phi_i) & , 0 < y_i < 1 \end{cases} \tag{S30}$$

and the parameter models

$$\mathrm{logit}(\mu_i) = \alpha_\mu + \mathrm{x}_i{}^T\beta_\mu + b_{s_i} \tag{S31}$$

$$\log(\phi_i) = \alpha_\phi + \mathrm{x}_i{}^T\beta_\phi \tag{S32}$$

$$\mathrm{logit}(\zeta_i) = \alpha_\zeta + \mathrm{x}_i{}^T\beta_\zeta \tag{S33}$$

$$\mathrm{logit}(\kappa_i) = \alpha_\kappa + \mathrm{x}_i{}^T\beta_\kappa \tag{S34}$$

The sensitivity of MU $i$ is denoted by $y_i$, $\mu_i$ denotes the beta component mean, $\phi_i$ is the beta precision parameter, with beta shape parameter $\mu_i\phi_i$ and $(1-\mu_i)\phi_i$, the zero-one inflation probability denotes by $\zeta_i$, and $\kappa_i$ is the conditional-one probability.

Priors for the mean submodel were

$$\alpha_\mu \sim \mathrm{Normal}(0,1) \tag{S35}$$

$$\beta_{\mu,k} \sim \mathrm{Normal}(0, 3^2),\ k = 1, \dots, K_\mu \tag{S36}$$

$$b_j|\tau_b \sim \mathrm{Normal}(0, \tau_b{}^2),\ j = 1, \dots, J \tag{S37}$$

$$\tau_b \sim \mathrm{Exponential}(1),\ \tau_b > 0 \tag{S38}$$

Here $\alpha_\mu$ denotes population-level intercept for the mean submodel, $\beta_{\mu,k}$ denotes population-level method effects for coefficient $k$, $b_j|\tau_b$ denotes MUset-specific random intercepts conditional on the between-MUset standard deviation, and $\tau_b$ denotes between-MUset standard deviation.

Priors for the precision submodel were

$$\alpha_\phi \sim \text{Normal}(\log(20), 1) \quad \text{(S39)}$$

$$\beta_{\phi,k} \sim \text{Normal}(0, 3^2),\ k = 1, \dots, K_\phi \quad \text{(S40)}$$

where $\alpha_\phi$ denotes precision-model intercept and $\beta_{\phi,k}$ denotes the method effects in the precision submodel.

Priors for the zero-one inflation submodel were

$$\alpha_\zeta \sim \text{Normal}(0,1) \quad \text{(S41)}$$

$$\beta_{\zeta,k} \sim \text{Normal}(0, 3^2),\ k = 1, \dots, K_\zeta \quad \text{(S42)}$$

where $\alpha_\zeta$ denotes zero-one inflation intercept and $\beta_{\zeta,k}$ denotes the method effects in the zero-one inflation submodel.

Finally, priors for the conditional one-inflation submodel were

$$\alpha_\kappa \sim \text{Normal}(0,1) \quad \text{(S43)}$$

$$\beta_{\kappa,k} \sim \text{Normal}(0, 3^2),\ k = 1, \dots, K_\kappa \quad \text{(S44)}$$

where $\alpha_\kappa$ denotes conditional one-inflation intercept and $\beta_{\kappa,k}$ denotes the method effects in the conditional one-inflation submodel.

Priors were specified to be weakly informative on the scale of the observed sensitivity and precision of the MU firing identification. These priors were evaluated using prior predictive simulations to confirm that the priors generated values within a plausible range (Supplementary Fig. 2a).

Model 2

In this model we additionally allowed the Method effect to vary by MUset by adding MUset-specific Method slopes. The model was implemented using the formula bf(Y ~ Method + (1 + Method | MUset), phi ~ Method, zoi ~ Method, coi ~ Method). Therefore

$$\text{logit}(\mu_i) = \alpha_\mu + \mathbf{x}_i^T \beta_\mu + \mathbf{r}_i^T \mathbf{b}_{s_i} \quad \text{(S45)}$$

where $\mathbf{r}_i$ is group-level design vector for observation $i$.

The likelihood and priors of Model 1 were used, except for $b_j|\tau_b \sim \text{Normal}(0, {\tau_b}^2)$ which was replaced with random intercept and random method effects defined as

$$\mathbf{b}_j \big| \tau_b, \mathbf{R} \sim \text{MultivariateNormal}\big(0, \text{diag}(\tau_b)\ \mathbf{R}\ \text{diag}(\tau_b)\big) \tag{S46}$$

with

$$\mathbf{b}_j = \left(b_{0j}, b_{1j}, \dots, b_{K_\mu j}\right)^{\mathrm{T}} \tag{S47}$$

where $\mathbf{R}$ contains the correlation between the random intercept and the random method effects. Furthermore, we additionally assigned an LKJ prior to the correlation matrix among the MUset-specific intercept and Method effects, defined as

$$\mathbf{R} \sim \text{LKJ}(2) \tag{S48}$$

Prior predictive simulations confirmed that the priors generated sensitivity values within a plausible range (Supplementary Fig. 2b).

<u>Model 3</u>

Compared to Model 2, this model additionally allowed observations from each MUset-MUID combination to have their own random intercept. The model was implemented using the formula bf(Y ~ Method + (1 + Method | MUset) + (1 | MUset:MUID), phi ~ Method, zoi ~ Method, coi ~ Method). Therefore,

$$\text{logit}(\mu_i) = \alpha_\mu + \mathbf{x}_i{}^T \beta_\mu + \mathbf{r}_i{}^T \mathbf{b}_{s_i} + c_{s_i l_i} \tag{S49}$$

Here $c_{s_i l_i}$ denotes random intercept for the MUset-MUID combination associated with observation $i$.

The likelihood and priors of Model 2 were used in this model because Model 2 included a broad prior for the class-level random-effect standard deviation (class = “sd”). The prior predictive check is shown in (Supplementary Fig. 2c).

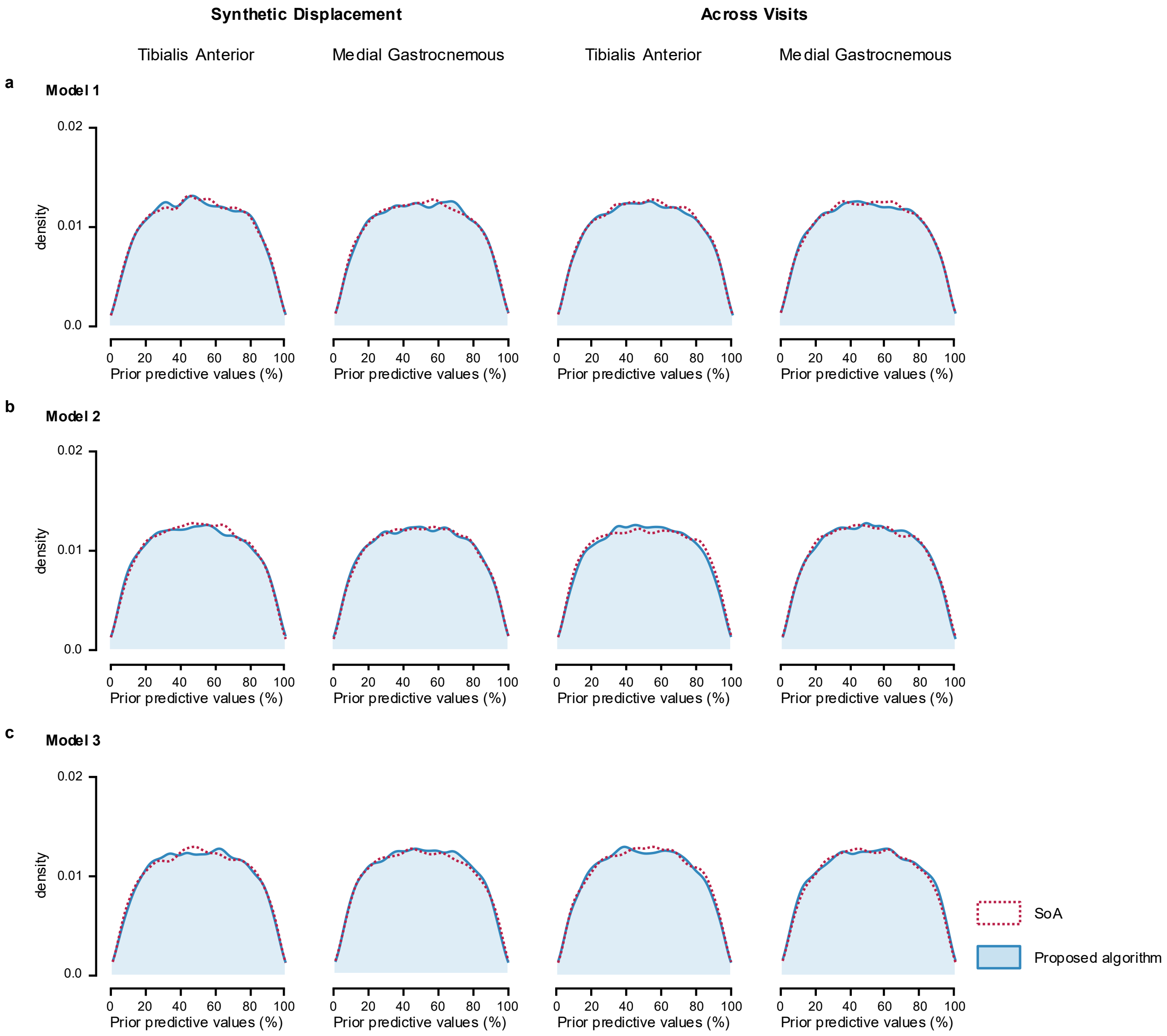


**Supplementary Fig. 2: Prior predictive check for Bayesian comparison on sensitivity and precision of the MU firing identification**

Prior predictive checks for the Bayesian comparison of SoA and the proposed algorithms on sensitivity and precision of the MU firing identification in tibialis anterior and medial gastrocnemius muscles, obtained by tracking MUs across synthetic displacement of electrode grid and two testing visits. Simulations were performed for **a**, Model 1; **b**, Model 2; and **c**, Model 3.

**Supplementary Note 6. Comparison of firing identification PNR for tracked TA MUs across synthetic grid displacement.**

Model implementation, likelihood function, and priors are detailed in Supplementary Note 5. The full dataset contains firing identification PNR values for 368 tracked MUs in the tibialis anterior (TA) muscle (Supplementary Fig. 3). The MUs firings were obtained by applying the MU filters calculated using the SoA algorithm[5] and the proposed algorithm to the HDsEMG signals generated with synthetic grid displacement.

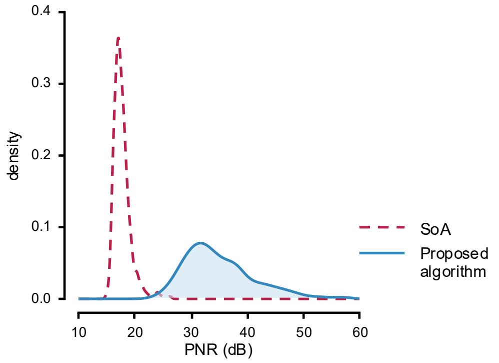


**Supplementary Fig. 3: Distribution of firing identification PNR in tracked TA MUs across synthetic grid displacement.**

PNR values were calculated from firing identification obtained by applying MU filters calculated using SoA (dashed line) and the proposed algorithms (solid line) to the HDsEMG signals generated with synthetic grid displacement.

In all models, the lognormal mean and sigma effects were exponentiated before the posterior summaries were calculated. Group-level standard deviations (SDs) are reported on their native model or link scales. Posterior diagnostics include the point estimate and upper confidence limit of the potential scale reduction factor (PSRF Pt and PSRF UpCI, respectively), as well as the effective sample size (ESS). Model selection was based on the difference in expected log pointwise predictive density (ELPD diff) and its standard error (SE diff).

Model 1

The maximum PSRF across all parameters was 1.001, indicating good convergence, while the minimum ESS was 6,327.809, indicating stable estimates of the bounds of the credible intervals (Supplementary Table 1). The medians, 95% equal-tailed interval (ETI) and 95% highest-density interval (HDI) are reported in Supplementary Table 2.

**Supplementary Table 1: Posterior MCMC diagnostics for Model 1 of firing identification PNR in tracked TA MUs across synthetic grid displacement.**

| | PSRF Pt | PSRF UpCI | ESS |
|---|---|---|---|
| $\alpha_{\mu}$ | 1.000 | 1.001 | 6,598.152 |
| $\alpha_{\sigma}$ | 1.000 | 1.000 | 2,8463.848 |
| $\beta_{\mu,\text{Proposed}}$ | 1.000 | 1.000 | 3,6178.124 |
| $\beta_{\sigma,\text{Proposed}}$ | 1.000 | 1.000 | 2,9771.300 |
| $\tau_b$ | 1.000 | 1.001 | 6,327.809 |

**Supplementary Table 2: Posterior summaries for Model 1 of firing identification PNR in tracked TA MUs across synthetic grid displacement.**

| | 50% | 95% ETI | | Mode | 95% HDI | |
|---|---|---|---|---|---|---|
| | | 2.5% | 97.5% | | low | high |
| $\alpha_{\mu}$ | 17.709 | 17.396 | 18.024 | 17.698 | 17.382 | 18.009 |
| $\alpha_{\sigma}$ | 1.952 | 1.918 | 1.987 | 1.953 | 1.918 | 1.987 |
| $\beta_{\mu,\text{Proposed}}$ | 0.073 | 0.067 | 0.079 | 0.073 | 0.067 | 0.079 |
| $\beta_{\sigma,\text{Proposed}}$ | 2.172 | 1.940 | 2.426 | 2.162 | 1.927 | 2.410 |
| $\tau_b$ | 0.041 | 0.029 | 0.058 | 0.041 | 0.028 | 0.057 |

Posterior predictive checks were performed by overlaying model-generated predictions on the observed data (Supplementary Fig. 4a). The posterior distributions (Supplementary Fig. 4b) indicated that the proposed algorithm achieved a higher PNR (35.0 dB, [34.2-35.8 dB]) than SoA algorithm (17.8 dB, [17.4-18.1 dB]). The posterior distribution of the difference excluded zero, and less than 2.5% of its distribution fell within the region of practical equivalence (ROPE).

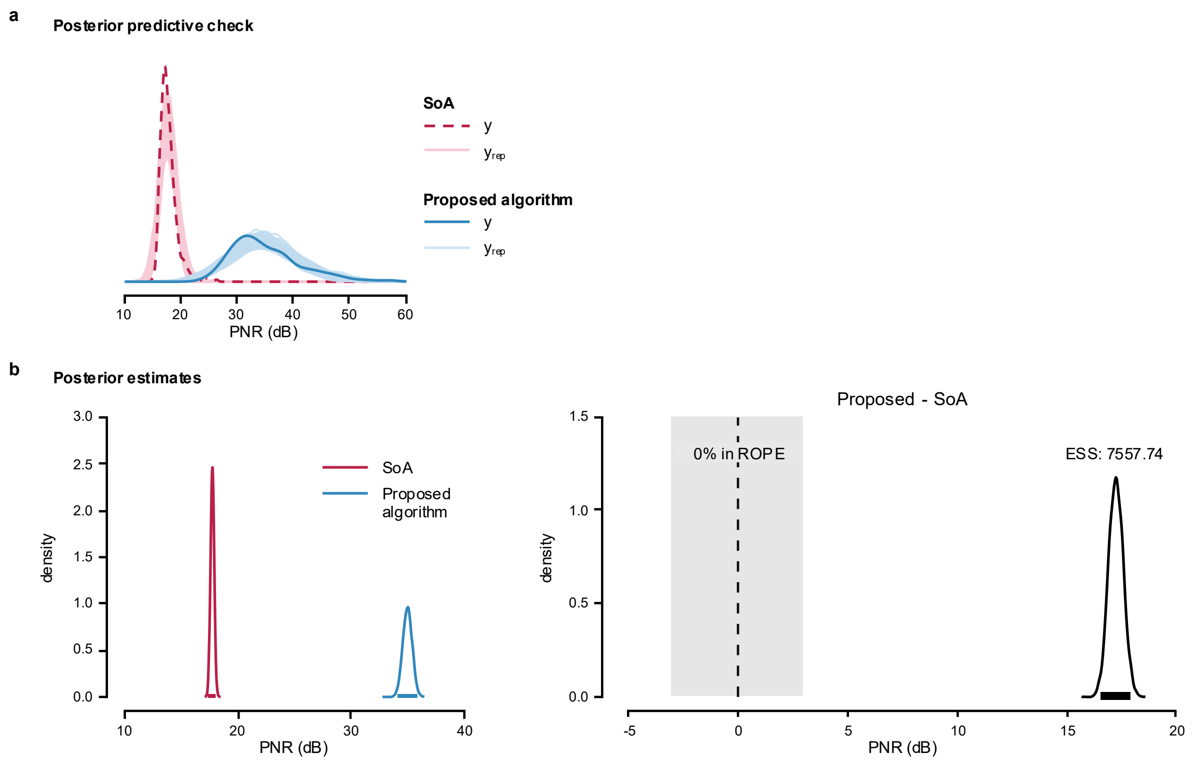


**Supplementary Fig. 4: Posterior predictive check and posterior estimates for Model 1 of firing identification PNR in tracked TA MUs across synthetic grid displacement.**

**a**, Posterior predictive check of the model; **b**, Posterior distributions and the difference between means estimated using Bayesian Model 1 to compare the performance of the proposed algorithm and SoA method.

Model 2

The maximum PSRF across all parameters was 1.005, indicating good convergence, while the minimum ESS was 3,858.575, indicating stable estimates of the bounds of the credible intervals (Supplementary Table 3). The medians, 95% ETIs, and 95% HDIs are reported in Supplementary Table 4. Group-level correlations were retained on their native scales.

Posterior predictive checks were performed by overlaying model-generated predictions on the observed data (Supplementary Fig. 5a). The posterior distributions (Supplementary Fig. 5b) indicated that the proposed algorithm achieved a higher PNR (34.1 dB, [32.6-35.6 dB]) than the SoA algorithm (17.8 dB, [17.5-18.2 dB]). The posterior distribution of the difference excluded zero, and less than 2.5% of its distribution fell within ROPE.

Model 3

The maximum PSRF across all parameters was 1.001, indicating good convergence, while the minimum ESS was 2,771.133, indicating stable estimates of the bounds of the credible intervals (Supplementary Table 5). Posterior summaries are presented in Supplementary Table 6 as medians with 95% ETIs and 95% HDIs. Correlations were summarized on their native scales.

Posterior predictive checks were performed by overlaying model-generated predictions on the observed data (Supplementary Fig. 6a). The posterior distributions (Supplementary Fig. 6b) indicated that the proposed algorithm achieved a higher PNR (34.1 dB, [32.6-35.7 dB]) than the SoA algorithm (17.8 dB, [17.5-18.2 dB]). The posterior distribution of the difference excluded zero, and less than 2.5% of its distribution fell within ROPE.

**Supplementary Table 3: Posterior MCMC diagnostics for Model 2 of firing identification PNR in tracked TA MUs across synthetic grid displacement.**

| | PSRF Pt | PSRF UpCI | ESS |
|---|---|---|---|
| $\alpha_{\mu}$ | 1.001 | 1.002 | 5,728.089 |
| $\alpha_{\sigma}$ | 1.000 | 1.000 | 2,0759.144 |
| $\beta_{\mu,\text{Proposed}}$ | 1.000 | 1.001 | 4,421.218 |
| $\beta_{\sigma,\text{Proposed}}$ | 1.000 | 1.001 | 2,1989.828 |
| $\tau_{b0}$ | 1.000 | 1.000 | 5,716.491 |
| $\tau_{b\text{Proposed}}$ | 1.000 | 1.000 | 5,140.098 |
| $R_{12}$ | 1.002 | 1.005 | 3,858.575 |

**Supplementary Table 4: Posterior summaries for Model 2 of firing identification PNR in tracked TA MUs across synthetic grid displacement.**

| | 50% | 95% ETI | | Mode | 95% HDI | |
|---|---|---|---|---|---|---|
| | | 2.5% | 97.5% | | low | high |
| $\alpha_{\mu}$ | 17.776 | 17.455 | 18.112 | 17.783 | 17.451 | 18.107 |
| $\alpha_{\sigma}$ | 0.071 | 0.066 | 0.076 | 0.070 | 0.065 | 0.076 |
| $\beta_{\mu,\text{Proposed}}$ | 1.903 | 1.813 | 1.989 | 1.907 | 1.814 | 1.989 |
| $\beta_{\sigma,\text{Proposed}}$ | 1.908 | 1.719 | 2.123 | 1.896 | 1.707 | 2.110 |
| $\tau_{b0}$ | 0.043 | 0.030 | 0.063 | 0.041 | 0.029 | 0.061 |
| $\tau_{b\text{Proposed}}$ | 0.112 | 0.081 | 0.158 | 0.109 | 0.079 | 0.154 |
| $R_{12}$ | -0.251 | -0.606 | 0.177 | -0.305 | -0.621 | 0.158 |

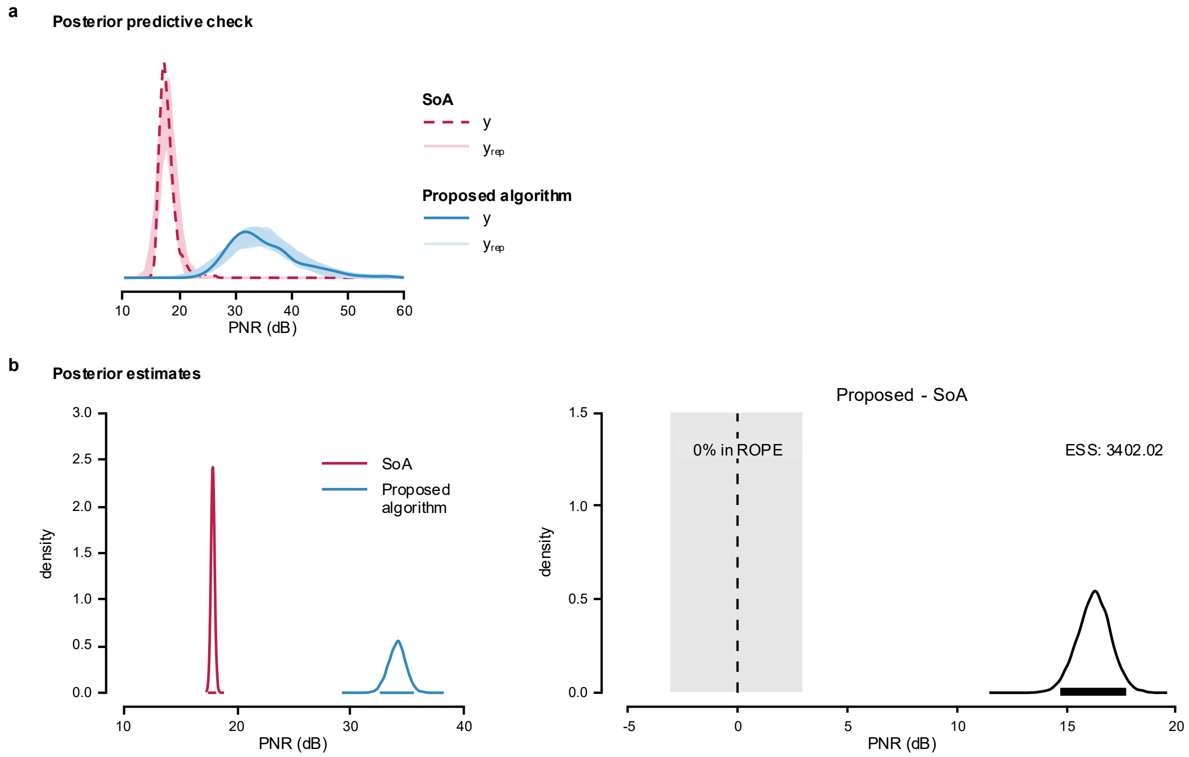


**Supplementary Fig. 5: Posterior predictive check and posterior estimates for Model 2 of firing identification PNR in tracked TA MUs across synthetic grid displacement.**

**a**, Posterior predictive check of the model; **b**, Posterior distributions and the difference in means from Bayesian Model 2, used to compare the performance of the proposed algorithm and SoA method.

**Supplementary Table 5: Posterior MCMC diagnostics for Model 3 of firing identification PNR in tracked TA MUs across synthetic grid displacement.**

| | PSRF Pt | PSRF UpCI | ESS |
|---|---|---|---|
| $\alpha_\mu$ | 1.000 | 1.000 | 8,436.545 |
| $\alpha_\sigma$ | 1.000 | 1.001 | 6,749.176 |
| $\beta_{\mu,\text{Proposed}}$ | 1.000 | 1.000 | 5,977.018 |
| $\beta_{\sigma,\text{Proposed}}$ | 1.000 | 1.001 | 1,3898.05 |
| $\tau_{b0}$ | 1.000 | 1.000 | 8,343.782 |
| $\tau_{b\text{Proposed}}$ | 1.000 | 1.000 | 6,636.922 |
| $\tau_c$ | 1.000 | 1.001 | 2,771.133 |
| $R_{12}$ | 1.000 | 1.001 | 5,044.315 |

**Supplementary Table 6: Posterior summaries for Model 3 of firing identification PNR in tracked TA MUs across synthetic grid displacement.**

| | 50% | 95% ETI | | Mode | 95% HDI | |
|---|---|---|---|---|---|---|
| | | 2.5% | 97.5% | | low | high |
| $\alpha_\mu$ | 17.771 | 17.446 | 18.112 | 17.765 | 17.447 | 18.113 |
| $\alpha_\sigma$ | 0.069 | 0.063 | 0.076 | 0.069 | 0.063 | 0.076 |
| $\beta_{\mu,\text{Proposed}}$ | 1.903 | 1.816 | 1.989 | 1.904 | 1.815 | 1.989 |
| $\beta_{\sigma,\text{Proposed}}$ | 1.938 | 1.739 | 2.182 | 1.915 | 1.733 | 2.174 |
| $\tau_{b0}$ | 0.043 | 0.030 | 0.062 | 0.042 | 0.029 | 0.060 |
| $\tau_{b\text{Proposed}}$ | 0.112 | 0.082 | 0.156 | 0.109 | 0.077 | 0.151 |
| $\tau_c$ | 0.012 | 0.001 | 0.031 | 0.005 | 0.000 | 0.028 |
| $R_{12}$ | -0.259 | -0.607 | 0.183 | -0.303 | -0.628 | 0.154 |

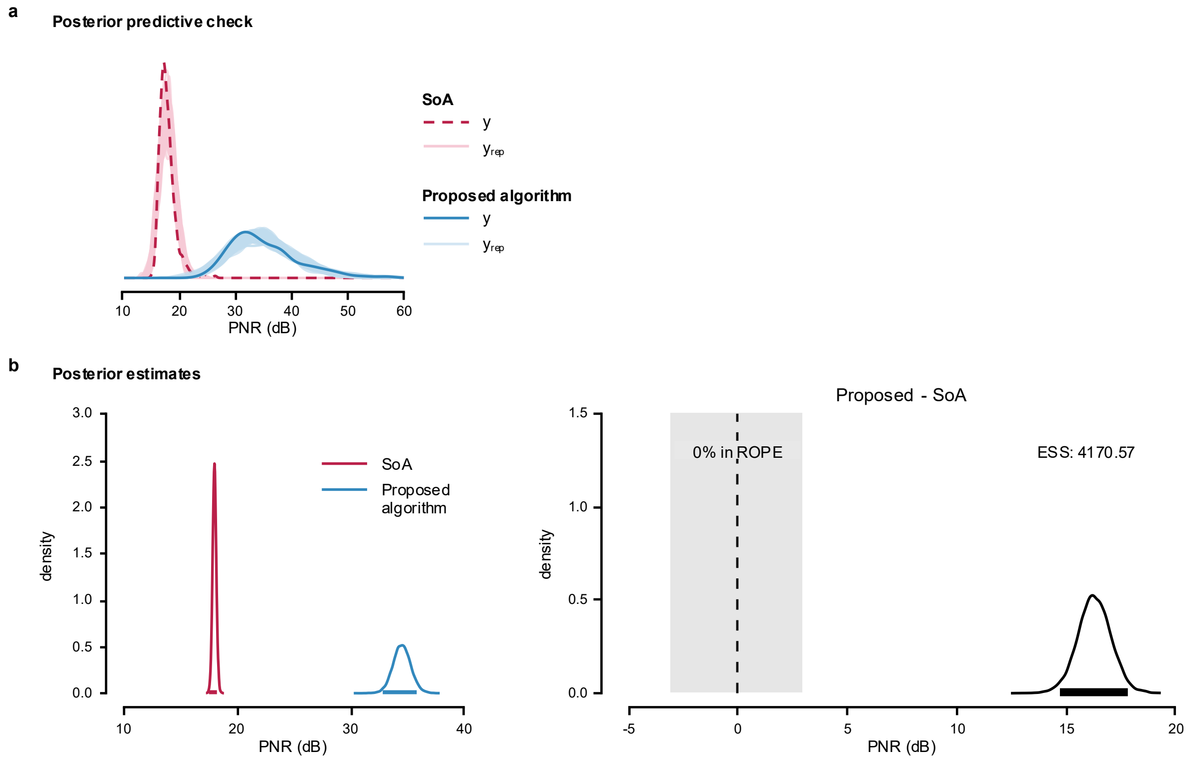


**Supplementary Fig. 6: Posterior predictive check and posterior estimates for Model 3 of firing identification PNR in tracked TA MUs across synthetic grid displacement.** **a**, Posterior predictive check of the model; **b**, Posterior distributions and the difference in means from Bayesian Model 3, used to compare the performance of the proposed algorithm and SoA method.

MODEL COMPARISON

Model 2 yielded the highest expected predictive accuracy. Relative to Model 2, the ELPD diff was -0.3 (SE = 0.7) for Model 3 and -57.1 (SE = 12.5) for Model 1, indicating comparable performance for Models 2 and 3 but poorer performance for Model 1.

**Supplementary Note 7. Comparison of firing identification sensitivity in tracked TA MUs across synthetic grid displacement.**

Model implementation, likelihood function, and priors are detailed in Supplementary Note 5. The full dataset contains firing identification sensitivity values for 368 tracked TA MUs (hereafter referred to as 'sensitivity'). The MUs firings were obtained by applying the MU filters calculated using the SoA algorithm[5] and the proposed algorithm to the HDsEMG signals generated with synthetic grid displacement. Distribution of the data is shown in Supplementary Fig. 7.

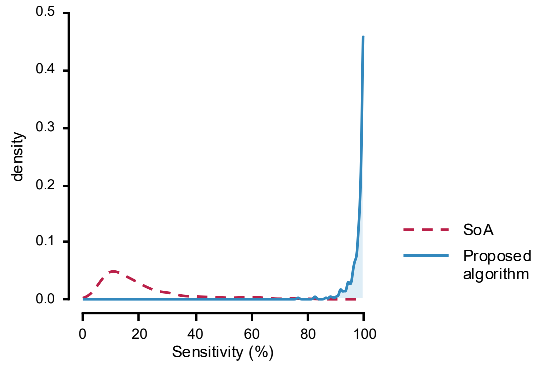


**Supplementary Fig. 7: Distribution of firing identification sensitivity in tracked TA MUs across synthetic grid displacement.**

Sensitivity values were calculated from firing identification obtained by applying MU filters calculated using SoA (dashed line) and the proposed algorithms (solid line) to the HDsEMG signals generated with synthetic grid displacement.

In all models, ZOIB parameters were back transformed into their interpretable component scales: *mu*, *zoi*, and *coi* to probabilities, and *phi* to the positive precision scale. Group-level SDs are reported on their native model/link scale.

Model 1

The maximum PSRF across all parameters was 1.001, indicating good convergence, while the minimum ESS was 5,390.882, indicating stable estimates of the bounds of the credible intervals (Supplementary Table 7). Posterior summaries are reported in Supplementary Table 8 as medians with 95% ETIs and 95% HDIs.

Posterior predictive checks were performed by overlaying model-generated predictions on the observed data (Supplementary Fig. 8a). The posterior distributions (Supplementary Fig. 8b) indicated that the proposed algorithm achieved a higher sensitivity (98.0%, [97.2-98.5 %]) than the SoA algorithm (20.9%, [18.6-23.3%]). The posterior distribution of the difference excluded zero, and less than 2.5% of its distribution fell within ROPE.

**Supplementary Table 7: Posterior MCMC diagnostics for Model 1 of firing identification sensitivity in tracked TA MUs across synthetic grid displacement.**

| | PSRF Pt | PSRF UpCI | ESS |
|---|---|---|---|
| $\alpha_{\mu}$ | 1.000 | 1.000 | 5,390.882 |
| $\alpha_{\phi}$ | 1.000 | 1.000 | 1,9961.068 |
| $\alpha_{\zeta}$ | 1.000 | 1.001 | 9,074.187 |
| $\alpha_{\kappa}$ | 1.000 | 1.000 | 1,4038.931 |
| $\beta_{\mu,\text{Proposed}}$ | 1.000 | 1.000 | 1,5246.186 |
| $\beta_{\phi,\text{Proposed}}$ | 1.000 | 1.001 | 1,4830.966 |
| $\beta_{\zeta,\text{Proposed}}$ | 1.000 | 1.001 | 9,149.810 |
| $\beta_{\kappa,\text{Proposed}}$ | 1.000 | 1.000 | 1,4782.282 |
| $\tau_{b}$ | 1.000 | 1.001 | 5,796.351 |

**Supplementary Table 8: Posterior summaries for Model 1 of firing identification sensitivity in tracked TA MUs across synthetic grid displacement.**

| | 50% | 95% ETI | | Mode | 95% HDI | |
|---|---|---|---|---|---|---|
| | | 2.5% | 97.5% | | low | high |
| $\alpha_{\mu}$ | 0.209 | 0.186 | 0.233 | 0.209 | 0.186 | 0.233 |
| $\alpha_{\phi}$ | 10.480 | 8.892 | 12.226 | 10.297 | 8.812 | 12.137 |
| $\alpha_{\zeta}$ | 0.005 | 0.001 | 0.014 | 0.003 | 0.000 | 0.012 |
| $\alpha_{\kappa}$ | 0.136 | 0.006 | 0.816 | 0.035 | 0.001 | 0.715 |
| $\beta_{\mu,\text{Proposed}}$ | 136.590 | 117.520 | 158.554 | 135.365 | 117.269 | 158.172 |
| $\beta_{\phi,\text{Proposed}}$ | 3.903 | 2.956 | 5.145 | 3.882 | 2.867 | 5.034 |
| $\beta_{\zeta,\text{Proposed}}$ | 154.881 | 49.605 | 808.805 | 91.756 | 26.332 | 592.249 |
| $\beta_{\kappa,\text{Proposed}}$ | 819.936 | 17.966 | 4,7147.870 | 3,725.074 | 0.763 | 2,4486.400 |
| $\tau_{b}$ | 0.303 | 0.189 | 0.453 | 0.306 | 0.180 | 0.441 |

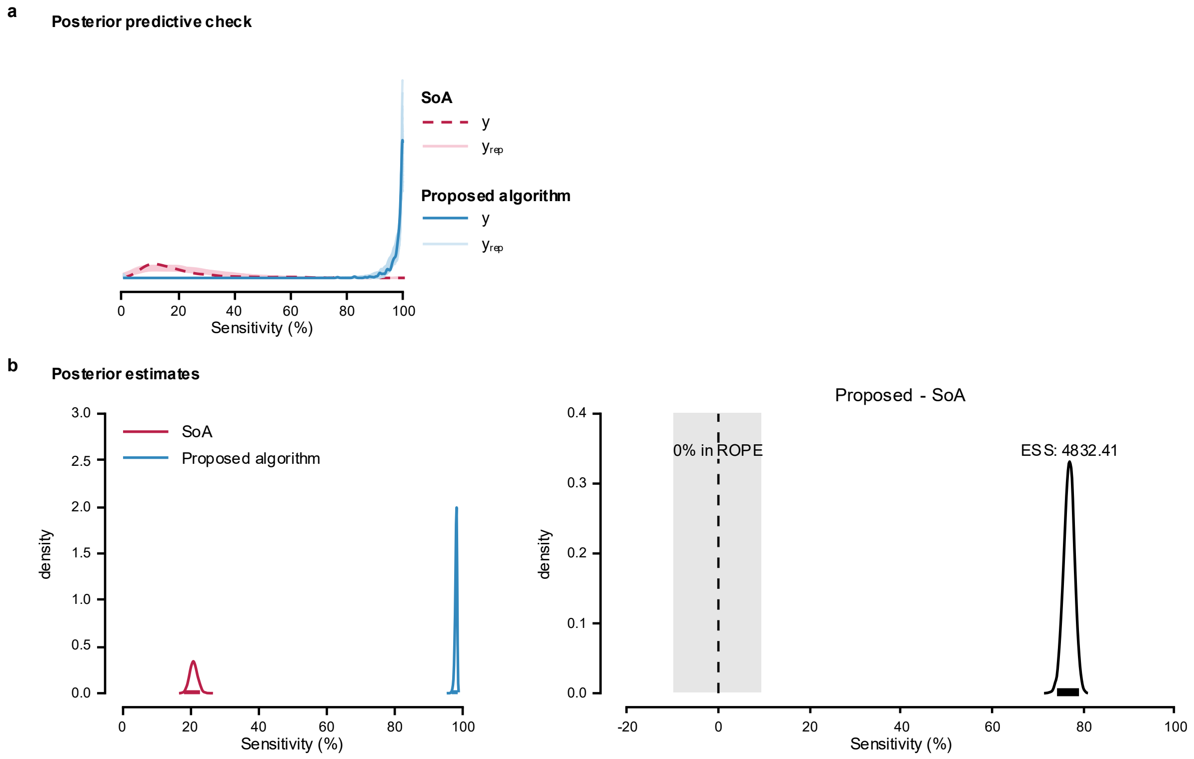


**Supplementary Fig. 8: Posterior predictive check and posterior estimates for Model 1 of firing identification sensitivity in tracked TA MUs across synthetic grid displacement.** **a**, Posterior predictive check of the model; **b**, Posterior distributions and the difference between means estimated using Bayesian Model 1 to compare the performance of the proposed algorithm and SoA method.

<u>Model 2</u>

The maximum PSRF across all parameters was 1.002, indicating good convergence, while the minimum ESS was 5,153.011, indicating stable estimates of the bounds of the credible intervals (Supplementary Table 9). Medians with 95% ETIs and 95% HDIs are presented in Supplementary Table 10. Group-level correlations are reported on the correlation scale.

Posterior predictive checks were performed by overlaying model-generated predictions on the observed data (Supplementary Fig. 9a). The posterior distributions (Supplementary Fig. 9b) indicated that the proposed algorithm achieved a higher sensitivity (97.9%, [97.1-98.4%])

than the SoA algorithm (21.2%, [18.5-24.2%]). The posterior distribution of the difference excluded zero, and less than 2.5% of its distribution fell within ROPE.

**Supplementary Table 9: Posterior MCMC diagnostics for Model 2 of firing identification sensitivity in tracked TA MUs across synthetic grid displacement.**

| | PSRF Pt | PSRF UpCI | ESS |
|---|---|---|---|
| $\alpha_{\mu}$ | 1.000 | 1.001 | 5,153.011 |
| $\alpha_{\phi}$ | 1.000 | 1.000 | 2,4832.207 |
| $\alpha_{\zeta}$ | 1.001 | 1.002 | 1,1001.276 |
| $\alpha_{\kappa}$ | 1.000 | 1.001 | 1,4437.182 |
| $\beta_{\mu,\text{Proposed}}$ | 1.001 | 1.002 | 6,605.306 |
| $\beta_{\phi,\text{Proposed}}$ | 1.000 | 1.001 | 1,7148.260 |
| $\beta_{\zeta,\text{Proposed}}$ | 1.001 | 1.002 | 1,1093.090 |
| $\beta_{\kappa,\text{Proposed}}$ | 1.000 | 1.001 | 1,5091.433 |
| $\tau_{b0}$ | 1.000 | 1.001 | 5,283.492 |
| $\tau_{b\text{Proposed}}$ | 1.000 | 1.000 | 6,925.981 |
| $R_{12}$ | 1.000 | 1.001 | 9,115.252 |

**Supplementary Table 10: Posterior summaries for Model 2 of firing identification sensitivity in tracked TA MUs across synthetic grid displacement.**

| | 50% | 95% ETI | | Mode | 95% HDI | |
|---|---|---|---|---|---|---|
| | | 2.5% | 97.5% | | low | high |
| $\alpha_{\mu}$ | 0.212 | 0.184 | 0.242 | 0.212 | 0.183 | 0.241 |
| $\alpha_{\phi}$ | 11.389 | 9.768 | 13.208 | 11.162 | 9.671 | 13.106 |
| $\alpha_{\zeta}$ | 0.005 | 0.001 | 0.014 | 0.003 | 0.000 | 0.012 |
| $\alpha_{\kappa}$ | 0.132 | 0.005 | 0.808 | 0.035 | 0.000 | 0.713 |
| $\beta_{\mu,\text{Proposed}}$ | 125.267 | 97.696 | 157.976 | 124.716 | 96.548 | 156.372 |
| $\beta_{\phi,\text{Proposed}}$ | 3.870 | 2.965 | 4.989 | 3.860 | 2.909 | 4.918 |
| $\beta_{\zeta,\text{Proposed}}$ | 155.744 | 48.843 | 823.357 | 106.188 | 26.608 | 608.646 |
| $\beta_{\kappa,\text{Proposed}}$ | 849.763 | 18.440 | 5,1723.880 | -859.945 | 0.506 | 2,6485.510 |
| $\tau_{b0}$ | 0.404 | 0.288 | 0.567 | 0.392 | 0.277 | 0.551 |
| $\tau_{b\text{Proposed}}$ | 0.486 | 0.301 | 0.720 | 0.463 | 0.285 | 0.700 |
| $R_{12}$ | -0.910 | -0.986 | -0.665 | -0.944 | -0.997 | -0.720 |

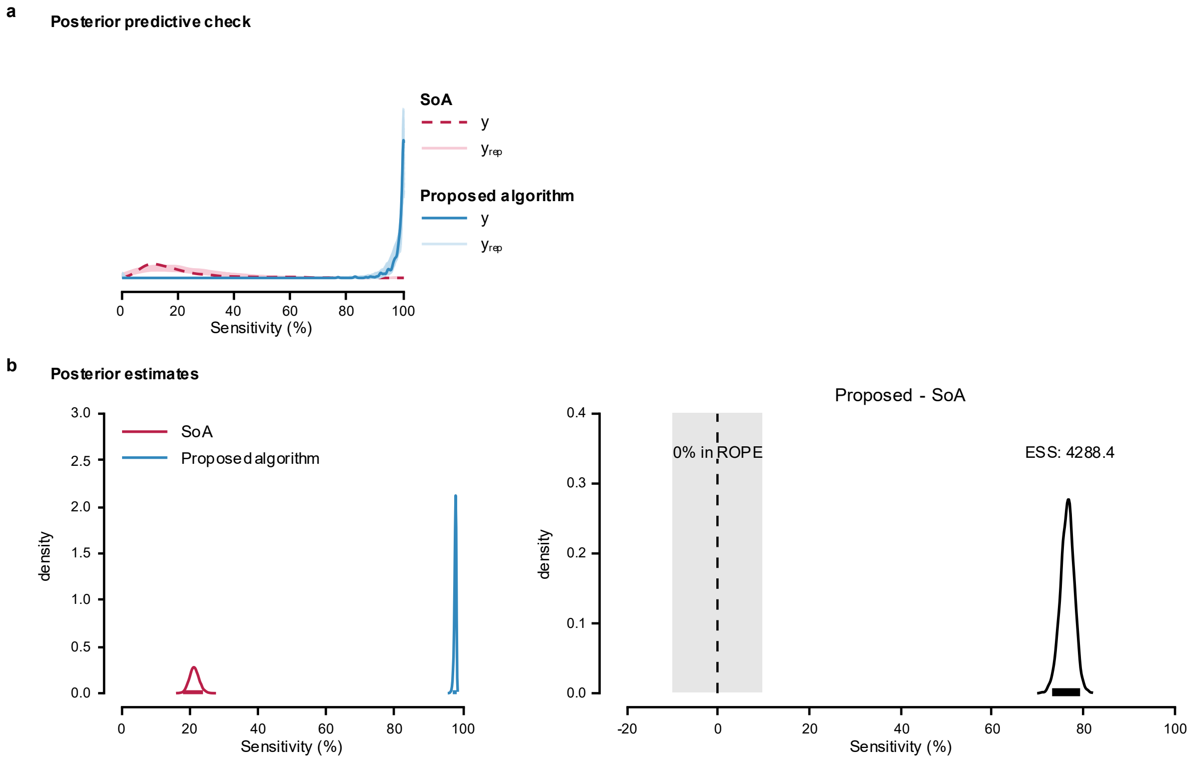


**Supplementary Fig. 9: Posterior predictive check and posterior estimates for Model 2 of firing identification sensitivity in tracked TA MUs across synthetic grid displacement.** **a**, Posterior predictive check of the model; **b**, Posterior distributions and the difference in means from Bayesian Model 2, used to compare the performance of the proposed algorithm and SoA method.

<u>Model 3</u>

The maximum PSRF across all parameters was 1.030, indicating good convergence, while the minimum ESS was 2,051.832, indicating stable estimates of the bounds of the credible intervals (Supplementary Table 11). Posterior estimates are reported in Supplementary Table 12 as medians with 95% ETIs and 95% HDIs. Correlations were summarized on their native scales.

Posterior predictive checks were performed by overlaying model-generated predictions on the observed data (Supplementary Fig. 10a). The posterior distributions (Supplementary Fig.

10b) indicated that the proposed algorithm achieved a higher sensitivity (97.9%, [97.1-98.4%]) than the SoA algorithm (21.2%, [18.2-24.3%]). The posterior distribution of the difference excluded zero, and less than 2.5% of its distribution fell within ROPE.

**Supplementary Table 11: Posterior MCMC diagnostics for Model 3 of firing identification sensitivity in tracked TA MUs across synthetic grid displacement.**

| | PSRF Pt | PSRF UpCI | ESS |
|---|---|---|---|
| $\alpha_{\mu}$ | 1.000 | 1.001 | 4,911.977 |
| $\alpha_{\phi}$ | 1.012 | 1.021 | 4,141.927 |
| $\alpha_{\zeta}$ | 1.000 | 1.000 | 1,2040.026 |
| $\alpha_{\kappa}$ | 1.000 | 1.000 | 2,0798.665 |
| $\beta_{\mu,\text{Proposed}}$ | 1.000 | 1.001 | 6,849.684 |
| $\beta_{\phi,\text{Proposed}}$ | 1.002 | 1.003 | 1,4848.922 |
| $\beta_{\zeta,\text{Proposed}}$ | 1.000 | 1.000 | 1,2402.239 |
| $\beta_{\kappa,\text{Proposed}}$ | 1.000 | 1.000 | 2,1512.980 |
| $\tau_{b0}$ | 1.001 | 1.003 | 6,115.368 |
| $\tau_{b\text{Proposed}}$ | 1.001 | 1.002 | 7,745.286 |
| $\tau_{c}$ | 1.014 | 1.030 | 2,051.832 |
| $R_{12}$ | 1.000 | 1.000 | 9,786.335 |

**Supplementary Table 12: Posterior summaries for Model 3 of firing identification sensitivity in tracked TA MUs across synthetic grid displacement.**

| | 50% | 95% ETI | | Mode | 95% HDI | |
|---|---|---|---|---|---|---|
| | | 2.5% | 97.5% | | low | high |
| $\alpha_{\mu}$ | 0.211 | 0.182 | 0.243 | 0.211 | 0.181 | 0.241 |
| $\alpha_{\phi}$ | 11.694 | 9.924 | 14.578 | 11.584 | 9.667 | 14.102 |
| $\alpha_{\zeta}$ | 0.005 | 0.001 | 0.014 | 0.003 | 0.000 | 0.012 |
| $\alpha_{\kappa}$ | 0.129 | 0.006 | 0.799 | 0.035 | 0.000 | 0.705 |
| $\beta_{\mu,\text{Proposed}}$ | 126.414 | 98.098 | 161.830 | 125.433 | 97.551 | 161.046 |
| $\beta_{\phi,\text{Proposed}}$ | 3.821 | 2.876 | 4.959 | 3.833 | 2.830 | 4.886 |
| $\beta_{\zeta,\text{Proposed}}$ | 154.619 | 49.252 | 781.366 | 96.331 | 32.067 | 583.902 |
| $\beta_{\kappa,\text{Proposed}}$ | 872.646 | 19.123 | 5,0303.580 | 2,539.043 | 0.507 | 2,5354.570 |
| $\tau_{b0}$ | 0.410 | 0.292 | 0.577 | 0.406 | 0.281 | 0.560 |
| $\tau_{b\text{Proposed}}$ | 0.497 | 0.314 | 0.733 | 0.476 | 0.314 | 0.731 |
| $\tau_{c}$ | 0.074 | 0.003 | 0.275 | 0.019 | 0.000 | 0.234 |
| $R_{12}$ | -0.910 | -0.987 | -0.664 | -0.944 | -0.997 | -0.723 |

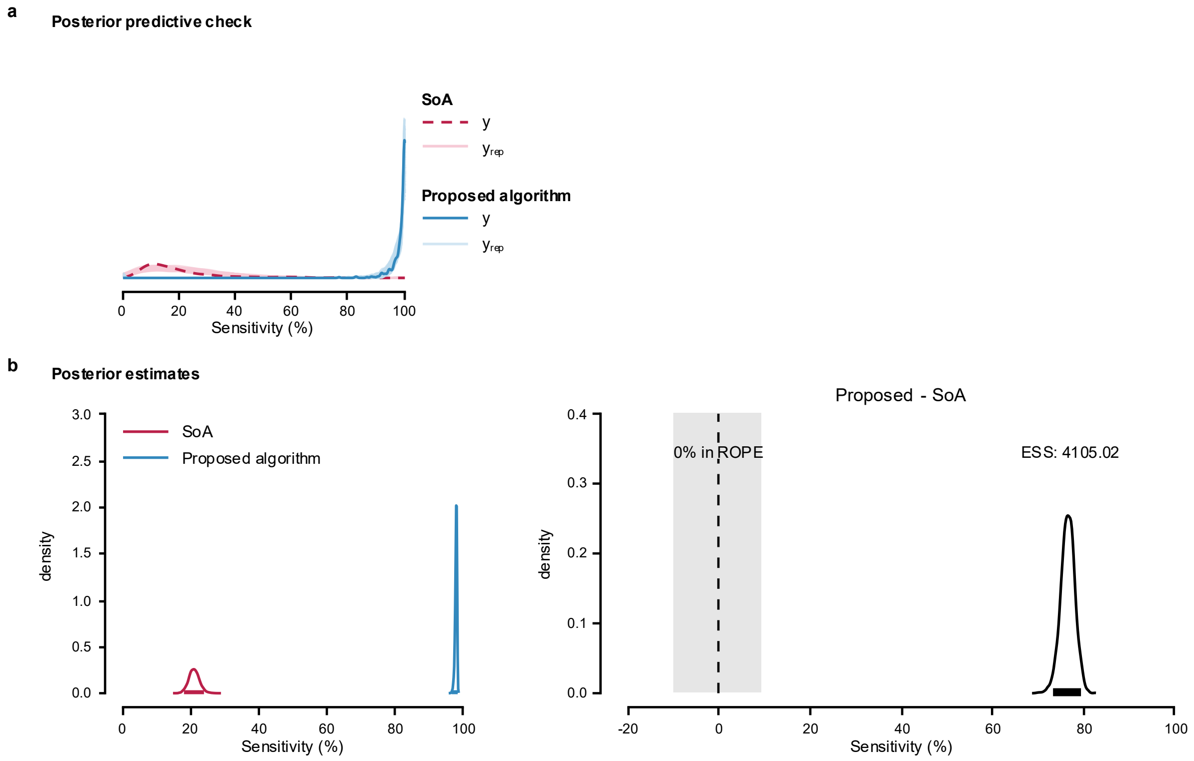


**Supplementary Fig. 10: Posterior predictive check and posterior estimates for Model 3 of firing identification sensitivity in tracked TA MUs across synthetic grid displacement.**

**a**, Posterior predictive check of the model; **b**, Posterior distributions and the difference in means from Bayesian Model 3, used to compare the performance of the proposed algorithm and SoA method.

MODEL COMPARISON

Model 2 showed the greatest expected predictive accuracy. ELPD diff relative to Model 2 were −0.5 (SE = 0.5) for Model 3 and −18.1 (SE = 5.5) for Model 1, suggesting similar predictive performance for Models 2 and 3 and reduced performance for Model 1.

**Supplementary Note 8. Comparison of firing identification precision in tracked TA MUs across synthetic grid displacement.**

Model implementation, likelihood function, and priors are detailed in Supplementary Note 5. The full dataset contains firing identification precision values for 368 tracked TA MUs (hereafter referred to as 'precision'). The MUs firings were obtained by applying the MU filters calculated using the SoA algorithm[5] and the proposed algorithm to the HDsEMG signals generated with synthetic grid displacement. Distribution of the data is shown in Supplementary Fig. 11.

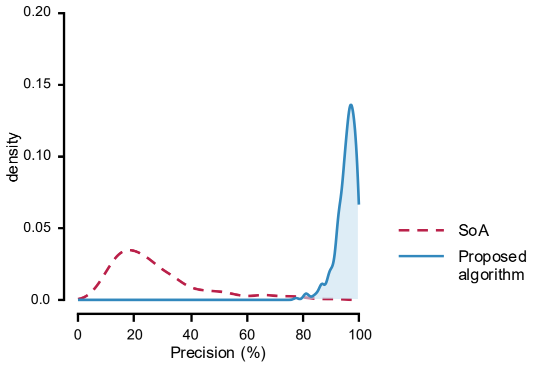


**Supplementary Fig. 11: Distribution of firing identification precision in tracked TA MUs across synthetic grid displacement.**

Precision values were calculated from firing identification obtained by applying MU filters calculated using SoA (dashed line) and the proposed algorithms (solid line) to the HDsEMG signals generated with synthetic grid displacement.

In all models, ZOIB parameters were back transformed into their interpretable component scales: *mu*, *zoi*, and *coi* to probabilities, and *phi* to the positive precision scale. Group-level SDs are reported on their native model/link scale.

Model 1

The maximum PSRF across all parameters was 1.002, indicating good convergence, while the minimum ESS was 6,125.286, indicating stable estimates of the bounds of the credible intervals (Supplementary Table 13). Supplementary Table 14 summarizes posterior estimates as medians with 95% ETIs and 95% HDIs.

Posterior predictive checks were performed by overlaying model-generated predictions on the observed data (Supplementary Fig. 12a). The posterior distributions (Supplementary Fig. 12b) indicated that the proposed algorithm achieved a higher precision (95.3%, [94.5-96.0 %]) than the SoA algorithm (29.0%, [26.5-31.6%]). The posterior distribution of the difference excluded zero, and less than 2.5% of its distribution fell within ROPE.

**Supplementary Table 13: Posterior MCMC diagnostics for Model 1 of firing identification precision in tracked TA MUs across synthetic grid displacement.**

| | PSRF Pt | PSRF UpCI | ESS |
|---|---|---|---|
| $\alpha_\mu$ | 1.000 | 1.000 | 1,0695.240 |
| $\alpha_\phi$ | 1.000 | 1.000 | 2,1607.562 |
| $\alpha_\zeta$ | 1.000 | 1.001 | 2,0201.825 |
| $\alpha_\kappa$ | 1.000 | 1.000 | 2,6771.201 |
| $\beta_{\mu,\text{Proposed}}$ | 1.000 | 1.000 | 2,8566.824 |
| $\beta_{\phi,\text{Proposed}}$ | 1.000 | 1.000 | 2,4113.835 |
| $\beta_{\zeta,\text{Proposed}}$ | 1.000 | 1.000 | 2,0145.884 |
| $\beta_{\kappa,\text{Proposed}}$ | 1.000 | 1.000 | 2,5032.027 |
| $\tau_b$ | 1.001 | 1.002 | 6,125.286 |

**Supplementary Table 14: Posterior summaries for Model 1 of firing identification precision in tracked TA MUs across synthetic grid displacement.**

| | 50% | 95% ETI | | Mode | 95% HDI | |
|---|---|---|---|---|---|---|
| | | 2.5% | 97.5% | | low | high |
| $\alpha_{\mu}$ | 0.290 | 0.266 | 0.316 | 0.291 | 0.265 | 0.315 |
| $\alpha_{\phi}$ | 8.211 | 7.012 | 9.557 | 8.121 | 7.004 | 9.543 |
| $\alpha_{\zeta}$ | 0.005 | 0.001 | 0.016 | 0.004 | 0.001 | 0.014 |
| $\alpha_{\kappa}$ | 0.214 | 0.010 | 0.880 | 0.049 | 0.000 | 0.813 |
| $\beta_{\mu,\text{Proposed}}$ | 49.916 | 44.774 | 55.675 | 49.997 | 44.801 | 55.701 |
| $\beta_{\phi,\text{Proposed}}$ | 4.902 | 3.896 | 6.152 | 4.891 | 3.799 | 6.030 |
| $\beta_{\zeta,\text{Proposed}}$ | 13.889 | 4.277 | 66.407 | 8.683 | 2.290 | 50.842 |
| $\beta_{\kappa,\text{Proposed}}$ | 109.270 | 2.046 | 7,927.556 | -122.207 | 0.030 | 3,782.908 |
| $\tau_{b}$ | 0.240 | 0.143 | 0.371 | 0.230 | 0.136 | 0.360 |

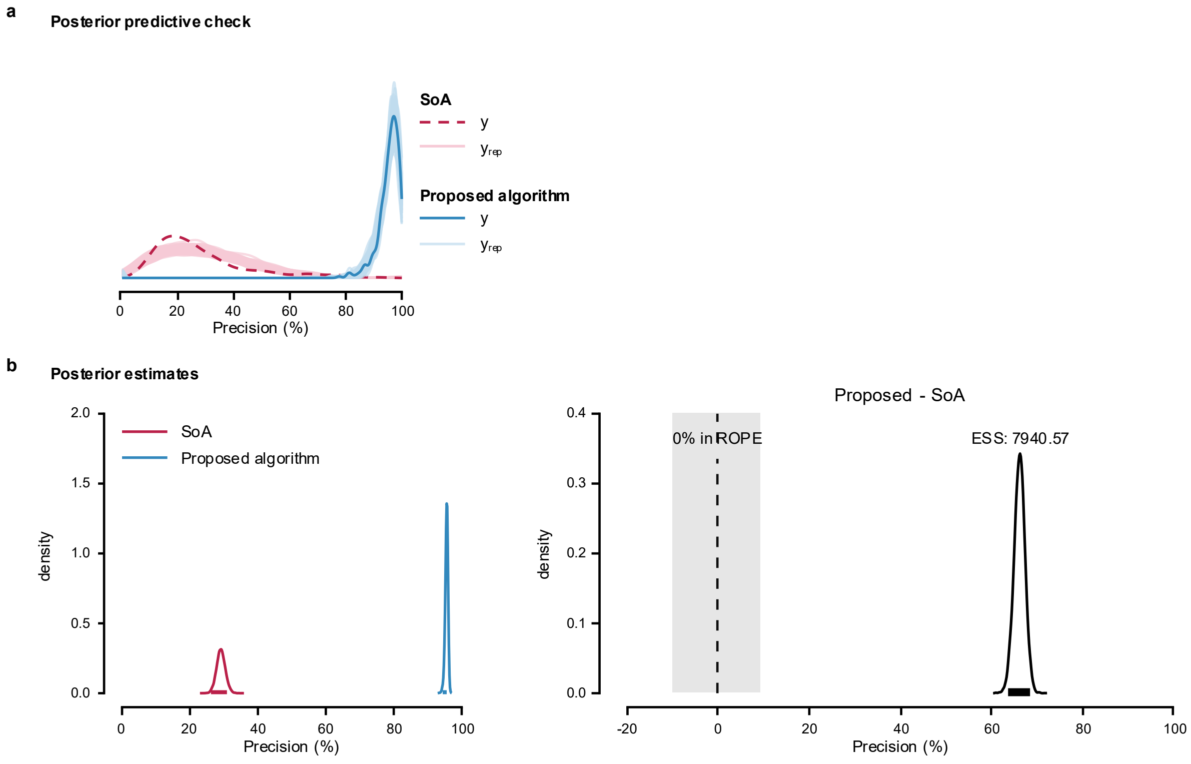


**Supplementary Fig. 12: Posterior predictive check and posterior estimates for Model 1 of firing identification precision in tracked TA MUs across synthetic grid displacement.** **a**, Posterior predictive check of the model; **b**, Posterior distributions and the difference between means estimated using Bayesian Model 1 to compare the performance of the proposed algorithm and SoA method.

<u>Model 2</u>

The maximum PSRF across all parameters was 1.003, indicating good convergence, while the minimum ESS was 4,034.166, indicating stable estimates of the bounds of the credible intervals (Supplementary Table 15). Posterior summaries are reported as medians with 95% ETIs and 95% HDIs on the original or component-specific interpretable scale (Supplementary Table 16). Group-level correlations are reported on the correlation scale.

Posterior predictive checks were performed by overlaying model-generated predictions on the observed data (Supplementary Fig. 13a). The posterior distributions (Supplementary Fig.

13b) indicated that the proposed algorithm achieved a higher precision (95.2%, [94.4-95.8%]) than the SoA algorithm (29.6%, [26.1-33.2%]). The posterior distribution of the difference excluded zero, and less than 2.5% of its distribution fell within ROPE.

**Supplementary Table 15: Posterior MCMC diagnostics for Model 2 of firing identification precision in tracked TA MUs across synthetic grid displacement.**

| | PSRF Pt | PSRF UpCI | ESS |
|---|---|---|---|
| $\alpha_{\mu}$ | 1.001 | 1.003 | 4,034.166 |
| $\alpha_{\phi}$ | 1.000 | 1.000 | 2,0916.869 |
| $\alpha_{\zeta}$ | 1.000 | 1.001 | 1,1142.008 |
| $\alpha_{\kappa}$ | 1.000 | 1.000 | 1,5214.434 |
| $\beta_{\mu,\text{Proposed}}$ | 1.000 | 1.001 | 5,560.573 |
| $\beta_{\phi,\text{Proposed}}$ | 1.000 | 1.000 | 2,1036.767 |
| $\beta_{\zeta,\text{Proposed}}$ | 1.000 | 1.001 | 1,1210.422 |
| $\beta_{\kappa,\text{Proposed}}$ | 1.000 | 1.000 | 1,7114.483 |
| $\tau_{b0}$ | 1.001 | 1.003 | 5,221.211 |
| $\tau_{b\text{Proposed}}$ | 1.000 | 1.001 | 6,010.290 |
| $R_{12}$ | 1.001 | 1.002 | 7,980.684 |

**Supplementary Table 16: Posterior summaries for Model 2 of firing identification precision in tracked TA MUs across synthetic grid displacement.**

| | 50% | 95% ETI | | Mode | 95% HDI | |
|---|---|---|---|---|---|---|
| | | 2.5% | 97.5% | | low | high |
| $\alpha_{\mu}$ | 0.295 | 0.262 | 0.332 | 0.294 | 0.262 | 0.332 |
| $\alpha_{\phi}$ | 9.262 | 7.962 | 10.712 | 9.213 | 7.938 | 10.679 |
| $\alpha_{\zeta}$ | 0.005 | 0.001 | 0.016 | 0.004 | 0.001 | 0.014 |
| $\alpha_{\kappa}$ | 0.208 | 0.010 | 0.876 | 0.049 | 0.001 | 0.810 |
| $\beta_{\mu,\text{Proposed}}$ | 47.337 | 39.257 | 56.773 | 47.143 | 39.068 | 56.558 |
| $\beta_{\phi,\text{Proposed}}$ | 4.468 | 3.588 | 5.557 | 4.439 | 3.505 | 5.446 |
| $\beta_{\zeta,\text{Proposed}}$ | 13.819 | 4.239 | 68.057 | 10.173 | 2.332 | 51.009 |
| $\beta_{\kappa,\text{Proposed}}$ | 110.880 | 2.127 | 8,030.003 | -118.164 | 0.081 | 3,843.218 |
| $\tau_{b0}$ | 0.400 | 0.278 | 0.565 | 0.394 | 0.267 | 0.549 |
| $\tau_{b\text{Proposed}}$ | 0.385 | 0.235 | 0.575 | 0.375 | 0.232 | 0.569 |
| $R_{12}$ | -0.877 | -0.974 | -0.621 | -0.916 | -0.987 | -0.668 |

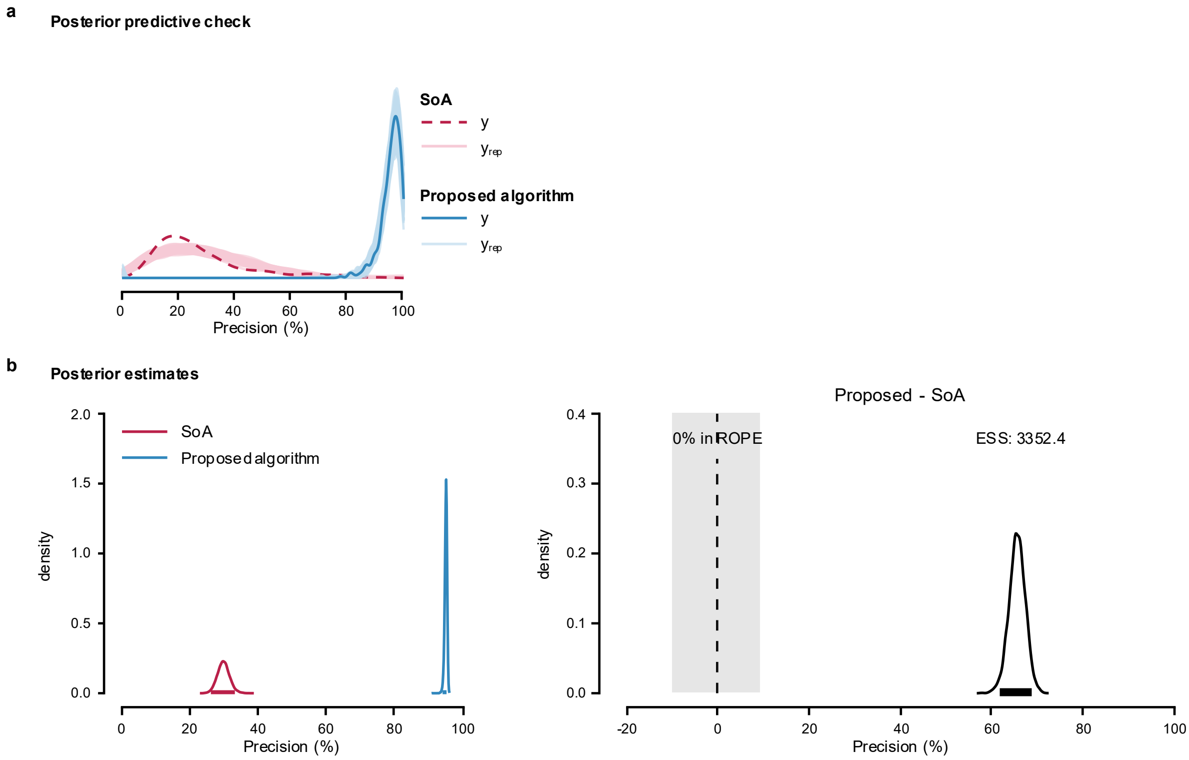


**Supplementary Fig. 13: Posterior predictive check and posterior estimates for Model 2 of firing identification precision in tracked TA MUs across synthetic grid displacement.** **a**, Posterior predictive check of the model; **b**, Posterior distributions and the difference between means estimated using Bayesian Model 2 to compare the performance of the proposed algorithm and SoA method.

<u>Model 3</u>

The maximum PSRF across all parameters was 1.001, indicating good convergence, while the minimum ESS was 3,553.260, indicating stable estimates of the bounds of the credible intervals (Supplementary Table 17). Posterior summaries are presented as medians with 95% ETIs and 95% HDIs (Supplementary Table 18). Group-level correlations were retained on their native scales.

Posterior predictive checks were performed by overlaying model-generated predictions on the observed data (Supplementary Fig. 14a). The posterior distributions (Supplementary Fig.

14b) indicated that the proposed algorithm achieved a higher precision (95.2%, [94.4-95.8%]) than the SoA algorithm (29.5%, [26.0-33.1%]). The posterior distribution of the difference excluded zero, and less than 2.5% of its distribution fell within ROPE.

**Supplementary Table 17: Posterior MCMC diagnostics for Model 3 of firing identification precision in tracked TA MUs across synthetic grid displacement.**

| | PSRF Pt | PSRF UpCI | ESS |
|---|---|---|---|
| $\alpha_{\mu}$ | 1.000 | 1.001 | 6,795.168 |
| $\alpha_{\phi}$ | 1.000 | 1.000 | 1,5773.480 |
| $\alpha_{\zeta}$ | 1.000 | 1.000 | 2,2353.260 |
| $\alpha_{\kappa}$ | 1.000 | 1.001 | 3,2691.363 |
| $\beta_{\mu,\text{Proposed}}$ | 1.000 | 1.001 | 8,952.588 |
| $\beta_{\phi,\text{Proposed}}$ | 1.000 | 1.000 | 3,3791.316 |
| $\beta_{\zeta,\text{Proposed}}$ | 1.000 | 1.000 | 2,1912.922 |
| $\beta_{\kappa,\text{Proposed}}$ | 1.000 | 1.001 | 2,9337.459 |
| $\tau_{b0}$ | 1.000 | 1.001 | 7,083.752 |
| $\tau_{b\text{Proposed}}$ | 1.000 | 1.001 | 9,589.969 |
| $\tau_{c}$ | 1.000 | 1.000 | 3,553.260 |
| $R_{12}$ | 1.000 | 1.001 | 1,1441.921 |

**Supplementary Table 18: Posterior summaries for Model 3 of firing identification precision in tracked TA MUs across synthetic grid displacement.**

| | 50% | 95% ETI | | Mode | 95% HDI | |
|---|---|---|---|---|---|---|
| | | 2.5% | 97.5% | | low | high |
| $\alpha_{\mu}$ | 0.294 | 0.260 | 0.331 | 0.292 | 0.259 | 0.330 |
| $\alpha_{\phi}$ | 9.450 | 8.091 | 11.097 | 9.290 | 8.057 | 11.039 |
| $\alpha_{\zeta}$ | 0.005 | 0.001 | 0.016 | 0.004 | 0.001 | 0.014 |
| $\alpha_{\kappa}$ | 0.212 | 0.009 | 0.881 | 0.049 | 0.000 | 0.817 |
| $\beta_{\mu,\text{Proposed}}$ | 47.747 | 39.405 | 57.501 | 47.403 | 38.789 | 56.728 |
| $\beta_{\phi,\text{Proposed}}$ | 4.475 | 3.587 | 5.563 | 4.502 | 3.545 | 5.509 |
| $\beta_{\zeta,\text{Proposed}}$ | 13.829 | 4.306 | 64.855 | 9.154 | 2.304 | 49.753 |
| $\beta_{\kappa,\text{Proposed}}$ | 110.214 | 2.057 | 8,939.829 | -123.690 | 0.030 | 4,369.792 |
| $\tau_{b0}$ | 0.403 | 0.286 | 0.572 | 0.394 | 0.281 | 0.562 |
| $\tau_{b\text{Proposed}}$ | 0.390 | 0.244 | 0.580 | 0.374 | 0.235 | 0.567 |
| $\tau_{c}$ | 0.069 | 0.003 | 0.208 | 0.019 | 0.000 | 0.185 |
| $R_{12}$ | -0.878 | -0.973 | -0.623 | -0.913 | -0.989 | -0.677 |

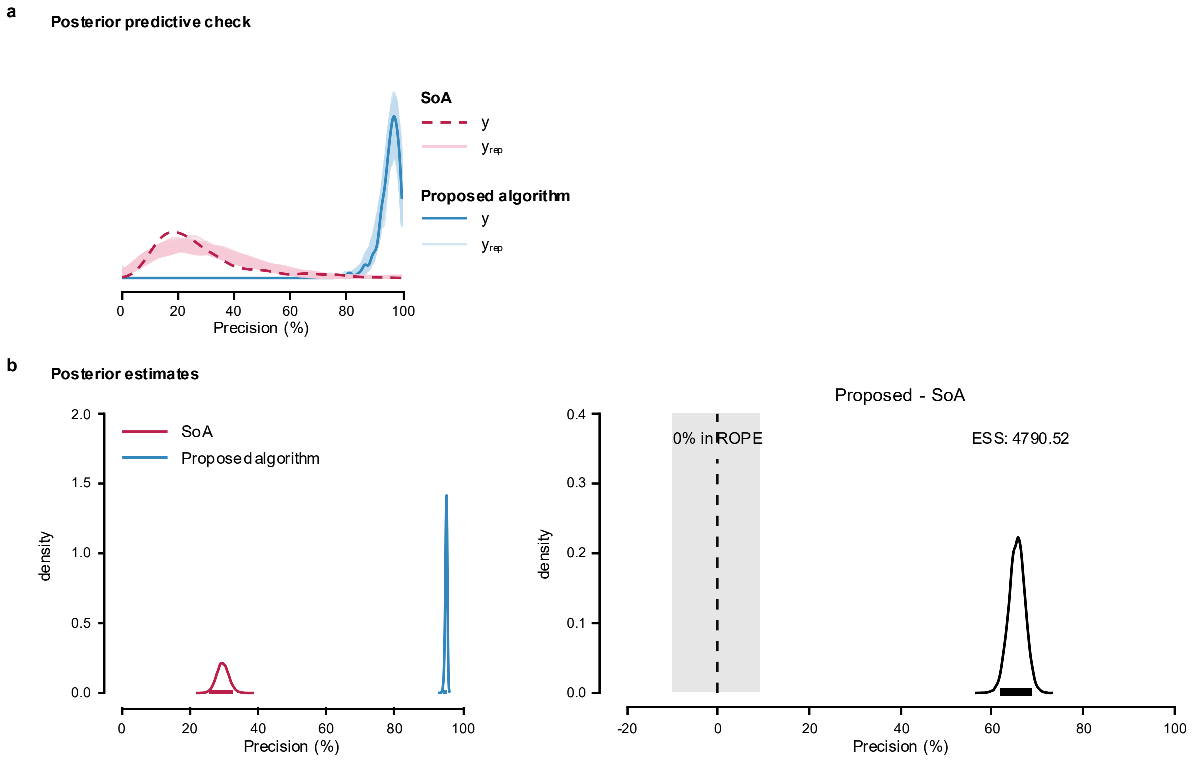


**Supplementary Fig. 14: Posterior predictive check and posterior estimates for Model 3 of firing identification precision in tracked TA MUs across synthetic grid displacement.** **a**, Posterior predictive check of the model; **b**, Posterior distributions and the difference between means estimated using Bayesian Model 1 to compare the performance of the proposed algorithm and SoA method.

MODEL COMPARISON

Model 2 showed the greatest expected predictive accuracy. ELPD diff relative to Model 2 were −0.7 (SE = 0.5) for Model 3 and −18.5 (SE = 5.8) for Model 1, suggesting similar predictive performance for Models 2 and 3 and reduced performance for Model 1.

**Supplementary Note 9. Comparison of firing identification PNR for tracked MG MUs across synthetic grid displacement.**

Model implementation, likelihood function, and priors are detailed in Supplementary Note 5. The full dataset contains firing identification PNR values for 300 tracked MUs in the medial gastrocnemius (MG) muscle (Supplementary Fig. 15). The MUs firings were obtained by applying the MU filters calculated using the SoA algorithm[5] and the proposed algorithm to the HDsEMG signals generated with synthetic grid displacement.

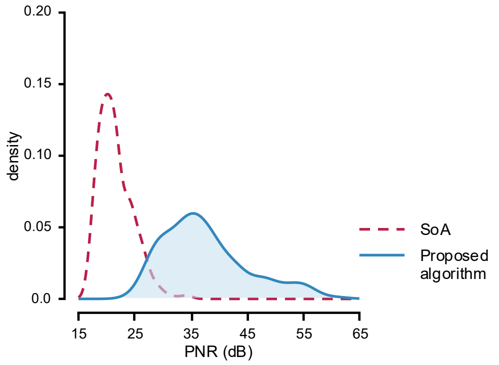


**Supplementary Fig. 15: Distribution of firing identification PNR in tracked MG MUs across synthetic grid displacement.**

PNR values were calculated from firing identification obtained by applying MU filters calculated using SoA (dashed line) and the proposed algorithms (solid line) to the HDsEMG signals generated with synthetic grid displacement.

In all models, the lognormal mean and sigma effects were exponentiated before the posterior summaries were calculated. Group-level standard deviations (SDs) are reported on their native model or link scales. Posterior diagnostics include the point estimate and upper confidence limit of the potential scale reduction factor (PSRF Pt and PSRF UpCI, respectively), as well as the effective sample size (ESS). Model selection was based on the difference in expected log pointwise predictive density (ELPD diff) and its standard error (SE diff).

Model 1

The maximum PSRF across all parameters was 1.001, indicating good convergence, while the minimum ESS was 6,091.227, indicating stable estimates of the bounds of the credible intervals (Supplementary Table 19). Posterior summaries are presented as medians with 95% ETI and the 95% HDI (Supplementary Table 20).

**Supplementary Table 19: Posterior MCMC diagnostics for Model 1 of firing identification PNR in tracked MG MUs across synthetic grid displacement.**

| | PSRF Pt | PSRF UpCI | ESS |
|---|---|---|---|
| $\alpha_{\mu}$ | 1.000 | 1.000 | 7,750.348 |
| $\alpha_{\sigma}$ | 1.000 | 1.000 | 2,0,672.677 |
| $\beta_{\mu,\text{Proposed}}$ | 1.000 | 1.000 | 3,2,281.335 |
| $\beta_{\sigma,\text{Proposed}}$ | 1.000 | 1.000 | 2,0,251.462 |
| $\tau_{\mu}$ | 1.000 | 1.001 | 6,091.227 |

**Supplementary Table 20: Posterior summaries for Model 1 of firing identification PNR in tracked MG MUs across synthetic grid displacement.**

| | 50% | 95% ETI | | Mode | 95% HDI | |
|---|---|---|---|---|---|---|
| | | 2.5% | 97.5% | | low | high |
| $\alpha_{\mu}$ | 21.721 | 21.085 | 22.387 | 21.703 | 21.070 | 22.370 |
| $\alpha_{\sigma}$ | 1.718 | 1.673 | 1.765 | 1.718 | 1.673 | 1.764 |
| $\beta_{\mu,\text{Proposed}}$ | 0.117 | 0.106 | 0.129 | 0.116 | 0.105 | 0.129 |
| $\beta_{\sigma,\text{Proposed}}$ | 1.726 | 1.481 | 1.996 | 1.711 | 1.483 | 1.998 |
| $\tau_{\mu}$ | 0.059 | 0.039 | 0.090 | 0.058 | 0.038 | 0.088 |

Posterior predictive checks were performed by overlaying model-generated predictions on the observed data (Supplementary Fig. 16a). The posterior distributions (Supplementary Fig. 16b) indicated that the proposed algorithm achieved a higher PNR (38.1 dB, [36.8-39.5 dB]) than the SoA algorithm (21.9 dB, [21.2-22.5 dB]). The posterior distribution of the difference excluded zero, and less than 2.5% of its distribution fell within ROPE.

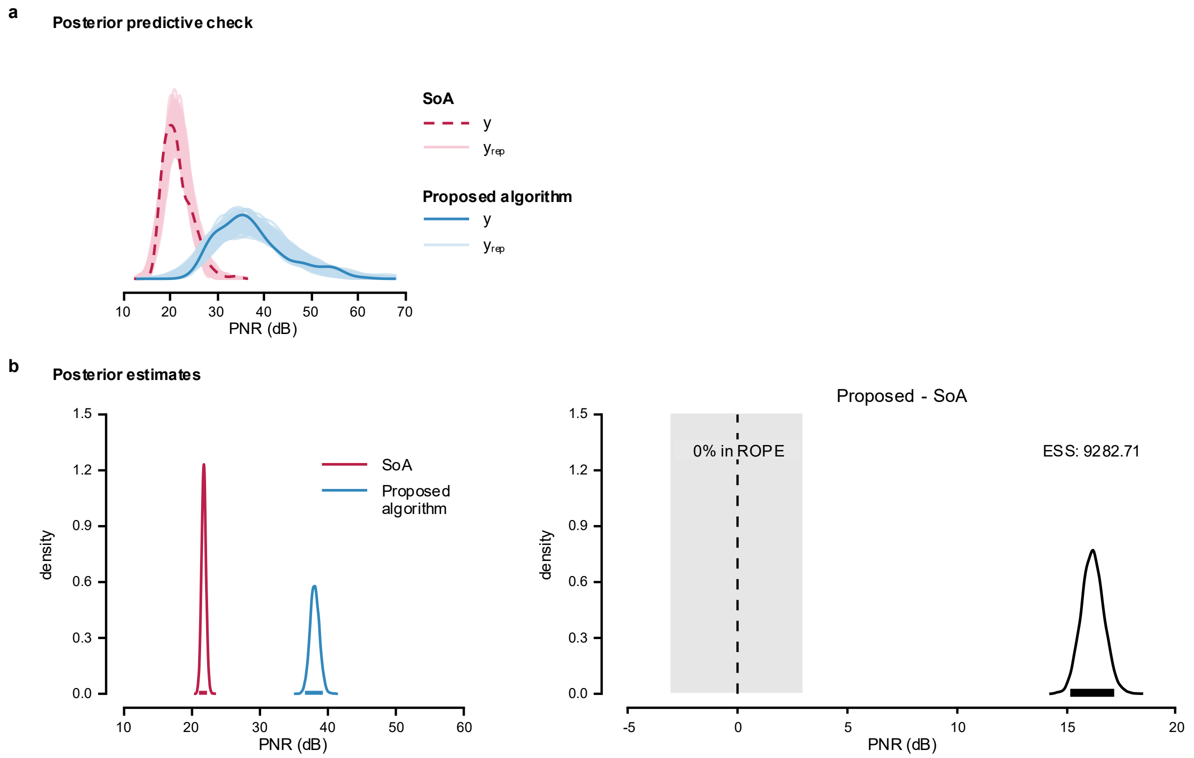


**Supplementary Fig. 16: Posterior predictive check and posterior estimates for Model 1 of firing identification PNR in tracked MG MUs across synthetic grid displacement.** **a**, Posterior predictive check of the model; **b**, Posterior distributions and the difference between means estimated using Bayesian Model 1 to compare the performance of the proposed algorithm and SoA method.

Model 2

The maximum PSRF across all parameters was 1.004, indicating good convergence, while the minimum ESS was 5,179.098, indicating stable estimates of the bounds of the credible intervals (Supplementary Table 21). Posterior summaries are reported as medians with 95% ETIs and HDIs. Group-level correlations were retained on their native scales (Supplementary Table 22).

Posterior predictive checks were performed by overlaying model-generated predictions on the observed data (Supplementary Fig. 17a). The posterior distributions (Supplementary Fig. 17b) indicated that the proposed algorithm achieved a higher PNR (35.9 dB, [33.6-38.2 dB]) than the SoA algorithm (22.3 dB, [21.4-23.2 dB]). The posterior distribution of the difference excluded zero, and less than 2.5% of its distribution fell within ROPE.

Model 3

The maximum PSRF across all parameters was 1.011, indicating good convergence, while the minimum ESS was 1,576.913, indicating stable estimates of the bounds of the credible intervals (Supplementary Table 23). Posterior estimates are summarized in Supplementary Table 24 by the median, 95% ETI and 95% HDI. Correlations were summarized on their native scales.

Posterior predictive checks were performed by overlaying model-generated predictions on the observed data (Supplementary Fig. 18a). The posterior distributions (Supplementary Fig. 18b) indicated that the proposed algorithm achieved a higher PNR (35.9 dB, [33.6-38.2 dB]) than the SoA algorithm (22.3 dB, [21.4-23.2 dB]). The posterior distribution of the difference excluded zero, and less than 2.5% of its distribution fell within ROPE.

**Supplementary Table 21: Posterior MCMC diagnostics for Model 2 of firing identification PNR in tracked MG MUs across synthetic grid displacement.**

| | PSRF Pt | PSRF UpCI | ESS |
|---|---|---|---|
| $\alpha_{\mu}$ | 1.001 | 1.001 | 5,308.435 |
| $\alpha_{\sigma}$ | 1.000 | 1.000 | 2,4,854.807 |
| $\beta_{\mu,\text{Proposed}}$ | 1.001 | 1.001 | 5,386.448 |
| $\beta_{\sigma,\text{Proposed}}$ | 1.000 | 1.001 | 2,5,237.093 |
| $\tau_{b0}$ | 1.001 | 1.002 | 5,263.395 |
| $\tau_{b\text{Proposed}}$ | 1.000 | 1.001 | 5,111.800 |
| $R_{12}$ | 1.001 | 1.004 | 5,179.098 |

**Supplementary Table 22: Posterior summaries for Model 2 of firing identification PNR in tracked MG MUs across synthetic grid displacement.**

| | 50% | 95% ETI | | Mode | 95% HDI | |
|---|---|---|---|---|---|---|
| | | 2.5% | 97.5% | | low | high |
| $\alpha_{\mu}$ | 22.163 | 21.303 | 23.098 | 22.149 | 21.278 | 23.062 |
| $\alpha_{\sigma}$ | 0.104 | 0.096 | 0.114 | 0.104 | 0.096 | 0.113 |
| $\beta_{\mu,\text{Proposed}}$ | 1.603 | 1.470 | 1.734 | 1.600 | 1.468 | 1.731 |
| $\beta_{\sigma,\text{Proposed}}$ | 1.456 | 1.296 | 1.631 | 1.464 | 1.292 | 1.624 |
| $\tau_{b0}$ | 0.087 | 0.064 | 0.125 | 0.084 | 0.061 | 0.121 |
| $\tau_{b\text{Proposed}}$ | 0.184 | 0.136 | 0.260 | 0.178 | 0.131 | 0.253 |
| $R_{12}$ | -0.689 | -0.872 | -0.337 | -0.738 | -0.896 | -0.395 |

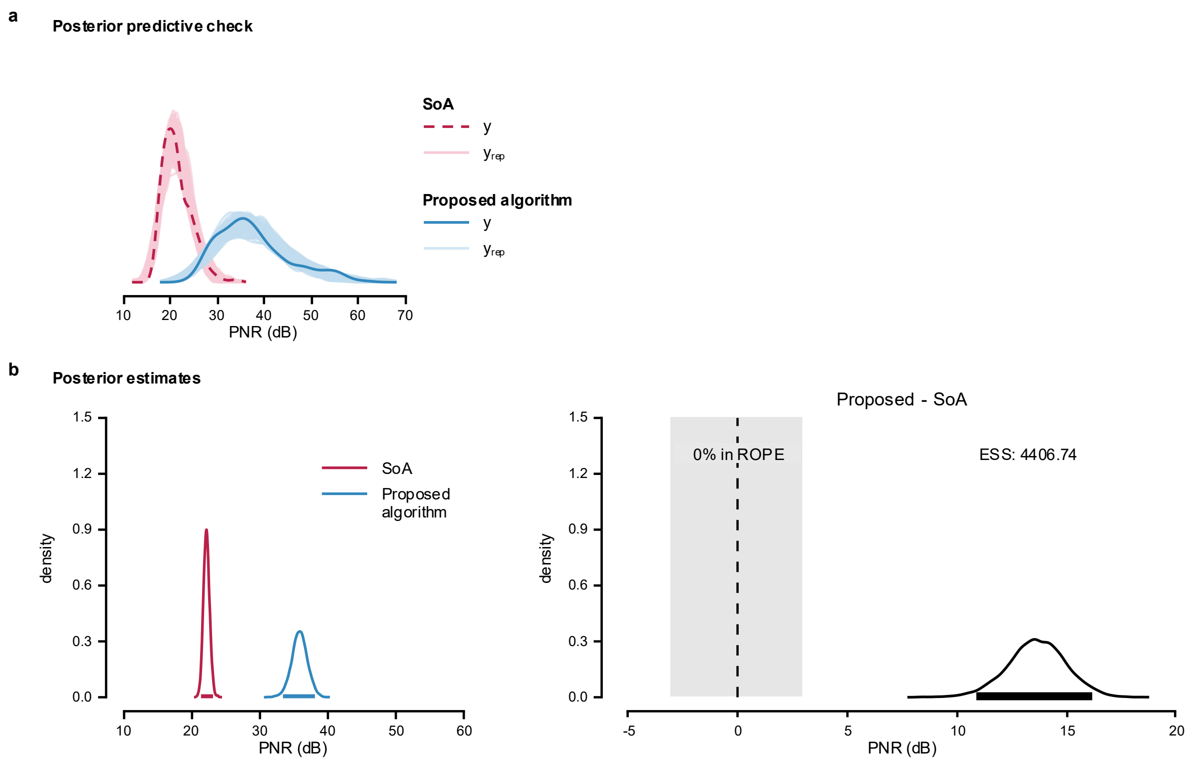


**Supplementary Fig. 17: Posterior predictive check and posterior estimates for Model 2 of firing identification PNR in tracked MG muscle MUs across synthetic grid displacement.**

**a**, Posterior predictive check of the model; **b**, Posterior distributions and the difference in means from Bayesian Model 2, used to compare the performance of the proposed algorithm and SoA method.

**Supplementary Table 23: Posterior MCMC diagnostics for Model 3 of firing identification PNR in tracked MG MUs across synthetic grid displacement.**

| | PSRF Pt | PSRF UpCI | ESS |
|---|---|---|---|
| $\alpha_{\mu}$ | 1.001 | 1.002 | 3,967.724 |
| $\alpha_{\sigma}$ | 1.001 | 1.004 | 3,130.600 |
| $\beta_{\mu,\text{Proposed}}$ | 1.000 | 1.001 | 4,121.262 |
| $\beta_{\sigma,\text{Proposed}}$ | 1.001 | 1.001 | 9,478.626 |
| $\tau_{b0}$ | 1.000 | 1.000 | 4,946.821 |
| $\tau_{b\text{Proposed}}$ | 1.000 | 1.000 | 4,814.260 |
| $\tau_{c}$ | 1.004 | 1.011 | 1,576.913 |
| $R_{12}$ | 1.002 | 1.004 | 4,792.227 |

**Supplementary Table 24: Posterior summaries for Model 3 of firing identification PNR in tracked MG MUs across synthetic grid displacement.**

| | 50% | 95% ETI | | Mode | 95% HDI | |
|---|---|---|---|---|---|---|
| | | 2.5% | 97.5% | | low | high |
| $\alpha_{\mu}$ | 22.160 | 21.307 | 23.111 | 22.169 | 21.318 | 23.121 |
| $\alpha_{\sigma}$ | 0.099 | 0.087 | 0.110 | 0.099 | 0.087 | 0.110 |
| $\beta_{\mu,\text{Proposed}}$ | 1.603 | 1.471 | 1.733 | 1.608 | 1.476 | 1.736 |
| $\beta_{\sigma,\text{Proposed}}$ | 1.495 | 1.318 | 1.718 | 1.478 | 1.313 | 1.710 |
| $\tau_{b0}$ | 0.088 | 0.064 | 0.127 | 0.085 | 0.060 | 0.122 |
| $\tau_{b\text{Proposed}}$ | 0.184 | 0.136 | 0.265 | 0.175 | 0.131 | 0.255 |
| $\tau_{c}$ | 0.033 | 0.003 | 0.056 | 0.038 | 0.000 | 0.053 |
| $R_{12}$ | -0.697 | -0.876 | -0.340 | -0.735 | -0.900 | -0.391 |

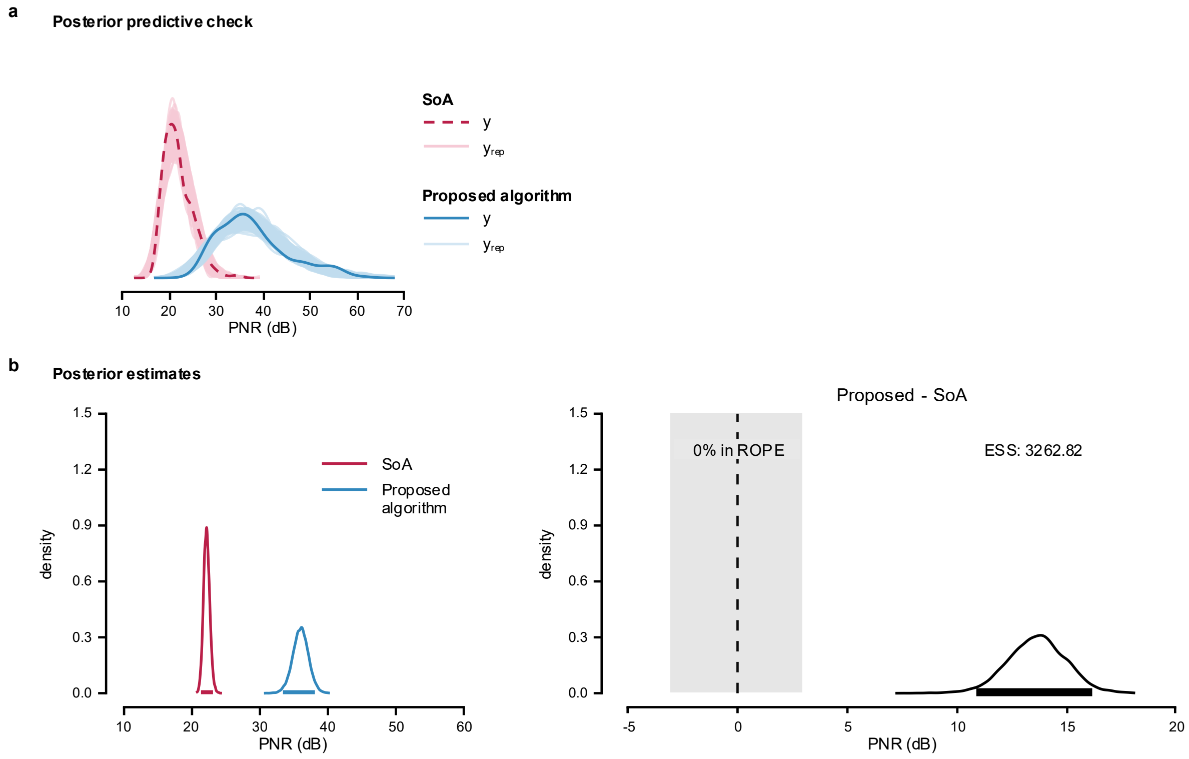


**Supplementary Fig. 18: Posterior predictive check and posterior estimates for Model 3 of firing identification PNR in tracked MG MUs across synthetic grid displacement.** **a**, Posterior predictive check of the model; **b**, Posterior distributions and the difference in means from Bayesian Model 3, used to compare the performance of the proposed algorithm and SoA method.

### MODEL COMPARISON

Model 3 showed the greatest expected predictive accuracy. ELPD diff relative to Model 3 were −1.6 (SE = 1.8) for Model 2 and −110.6 (SE = 13.5) for Model 1, suggesting similar predictive performance for Models 2 and 3 and reduced performance for Model 1.

**Supplementary Note 10. Comparison of firing identification sensitivity in tracked MG MUs across synthetic grid displacement.**

Model implementation, likelihood function, and priors are detailed in Supplementary Note 5. The full dataset contains firing identification sensitivity values for 300 tracked MG MUs (hereafter referred to as 'sensitivity'). The MUs firings were obtained by applying the MU filters calculated using the SoA algorithm[5] and the proposed algorithm to the HDsEMG signals generated with synthetic grid displacement. Distribution of the data is shown in Supplementary Fig. 19.

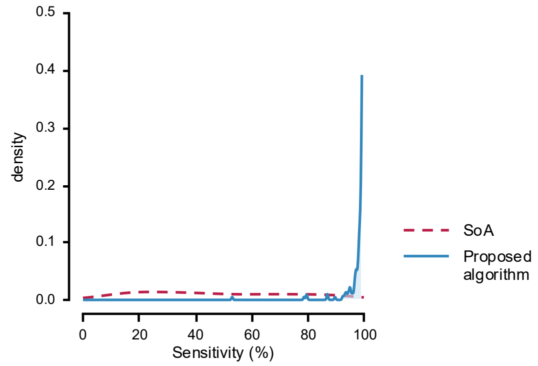


**Supplementary Fig. 19: Distribution of firing identification sensitivity in tracked MG MUs across synthetic grid displacement.**

Sensitivity values were calculated from firing identification obtained by applying MU filters calculated using SoA (dashed line) and the proposed algorithms (solid line) to the HDsEMG signals generated with synthetic grid displacement.

In all models, ZOIB parameters were back transformed into their interpretable component scales: *mu*, *zoi*, and *coi* to probabilities, and *phi* to the positive precision scale. Group-level SDs are reported on their native model/link scale.

Model 1

The maximum PSRF across all parameters was 1.004, indicating good convergence, while the minimum ESS was 5,659.884, indicating stable estimates of the bounds of the credible intervals (Supplementary Table 25). Posterior summaries are reported in Supplementary Table 26 as medians with 95% ETIs and 95% HDIs.

Posterior predictive checks were performed by overlaying model-generated predictions on the observed data (Supplementary Fig. 20a). The posterior distributions (Supplementary Fig. 20b) indicated that the proposed algorithm achieved a higher sensitivity (98.3%, [97.3-98.9 %]) than the SoA algorithm (51.0%, [46.0-56.3%]). The posterior distribution of the difference excluded zero, and less than 2.5% of its distribution fell within ROPE.

**Supplementary Table 25: Posterior MCMC diagnostics for Model 1 of firing identification sensitivity in tracked MG MUs across synthetic grid displacement.**

| | PSRF Pt | PSRF UpCI | ESS |
|---|---|---|---|
| $\alpha_{\mu}$ | 1.001 | 1.003 | 5,659.884 |
| $\alpha_{\phi}$ | 1.000 | 1.000 | 2,7610.853 |
| $\alpha_{\zeta}$ | 1.000 | 1.001 | 1,0711.309 |
| $\alpha_{\kappa}$ | 1.000 | 1.001 | 1,4355.252 |
| $\beta_{\mu,\mathrm{Proposed}}$ | 1.000 | 1.000 | 1,4347.463 |
| $\beta_{\phi,\mathrm{Proposed}}$ | 1.000 | 1.000 | 1,4874.786 |
| $\beta_{\zeta,\mathrm{Proposed}}$ | 1.000 | 1.001 | 1,1259.577 |
| $\beta_{\kappa,\mathrm{Proposed}}$ | 1.000 | 1.001 | 1,5329.729 |
| $\tau_{b}$ | 1.001 | 1.004 | 6,270.831 |

**Supplementary Table 26: Posterior summaries for Model 1 of firing identification sensitivity in tracked MG MUs across synthetic grid displacement.**

| | 50% | 95% ETI | | Mode | 95% HDI | |
|---|---|---|---|---|---|---|
| | | 2.5% | 97.5% | | low | high |
| $\alpha_{\mu}$ | 0.512 | 0.461 | 0.564 | 0.511 | 0.461 | 0.564 |
| $\alpha_{\phi}$ | 3.672 | 3.110 | 4.309 | 3.641 | 3.091 | 4.286 |
| $\alpha_{\zeta}$ | 0.005 | 0.001 | 0.018 | 0.004 | 0.000 | 0.015 |
| $\alpha_{\kappa}$ | 0.131 | 0.005 | 0.818 | 0.035 | 0.000 | 0.721 |
| $\beta_{\mu,\text{Proposed}}$ | 41.295 | 33.249 | 50.844 | 41.114 | 32.674 | 50.109 |
| $\beta_{\phi,\text{Proposed}}$ | 7.995 | 5.802 | 10.869 | 7.864 | 5.691 | 10.672 |
| $\beta_{\zeta,\text{Proposed}}$ | 167.521 | 49.887 | 872.358 | 101.611 | 29.368 | 654.505 |
| $\beta_{\kappa,\text{Proposed}}$ | 804.378 | 17.382 | 4,7843.100 | -828.998 | 0.506 | 2,4758.800 |
| $\tau_b$ | 0.391 | 0.251 | 0.604 | 0.374 | 0.237 | 0.580 |

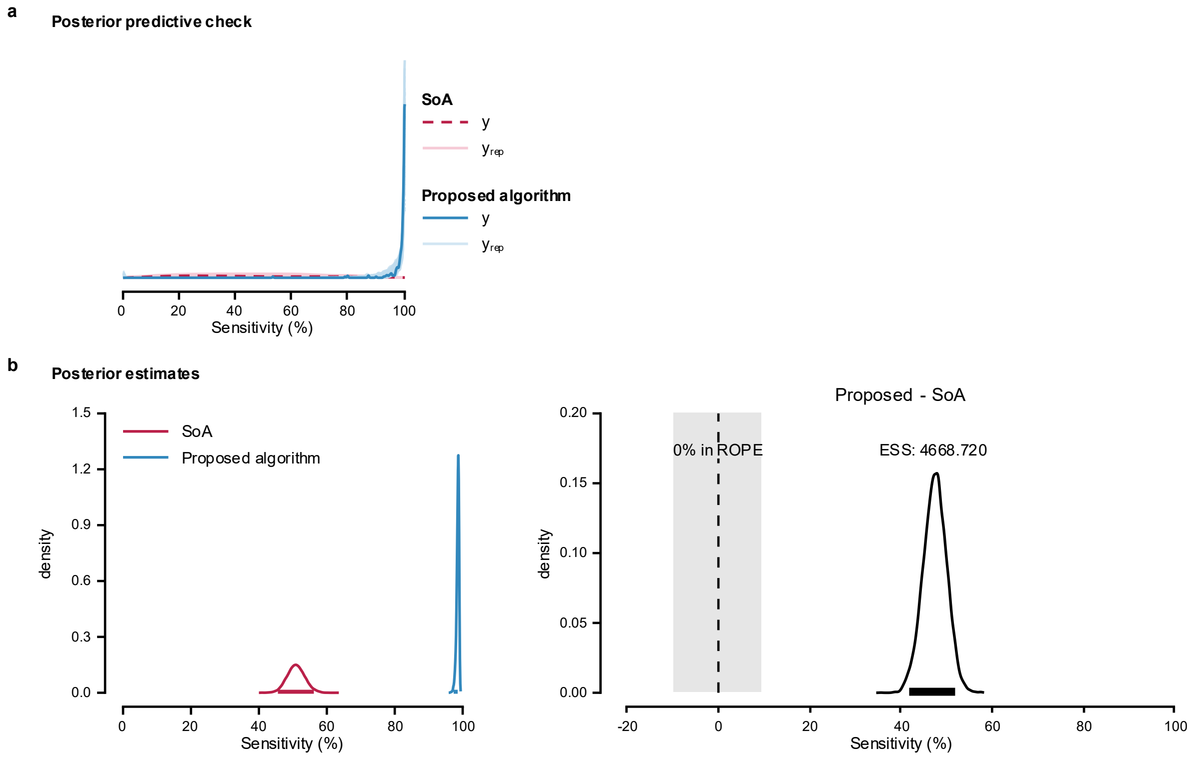


**Supplementary Fig. 20: Posterior predictive check and posterior estimates for Model 1 of firing identification sensitivity in tracked MG MUs across synthetic grid displacement.**

**a**, Posterior predictive check of the model; **b**, Posterior distributions and the difference between means estimated using Bayesian Model 1 to compare the performance of the proposed algorithm and SoA method.

<u>Model 2</u>

The maximum PSRF across all parameters was 1.002, indicating good convergence, while the minimum ESS was 5,153.011, indicating stable estimates of the bounds of the credible intervals (Supplementary Table 27). The medians with 95% ETIs and 95% HDIs of the posterior estimates are presented in Supplementary Table 28. Group-level correlations were summarized without transformation.

Posterior predictive checks were performed by overlaying model-generated predictions on the observed data (Supplementary Fig. 21a). The posterior distributions (Supplementary Fig. 21b) indicated that the proposed algorithm achieved a higher sensitivity (98.2%, [97.2-98.8%]) than the SoA algorithm (53.4%, [46.7-60.1%]). The posterior distribution of the difference excluded zero, and less than 2.5% of its distribution fell within ROPE.

**Supplementary Table 27: Posterior MCMC diagnostics for Model 2 of firing identification sensitivity in tracked MG MUs across synthetic grid displacement.**

| | PSRF Pt | PSRF UpCI | ESS |
|---|---|---|---|
| $\alpha_{\mu}$ | 1.001 | 1.002 | 4,552.855 |
| $\alpha_{\phi}$ | 1.000 | 1.001 | 2,9216.538 |
| $\alpha_{\zeta}$ | 1.000 | 1.001 | 1,0302.998 |
| $\alpha_{\kappa}$ | 1.000 | 1.001 | 1,4547.824 |
| $\beta_{\mu,\text{Proposed}}$ | 1.000 | 1.001 | 6,159.842 |
| $\beta_{\phi,\text{Proposed}}$ | 1.000 | 1.000 | 1,4114.821 |
| $\beta_{\zeta,\text{Proposed}}$ | 1.000 | 1.000 | 1,0420.945 |
| $\beta_{\kappa,\text{Proposed}}$ | 1.000 | 1.001 | 1,5274.517 |
| $\tau_{b0}$ | 1.001 | 1.002 | 5,141.714 |
| $\tau_{b\text{Proposed}}$ | 1.001 | 1.002 | 6,820.759 |
| $R_{12}$ | 1.000 | 1.001 | 1,0623.000 |

**Supplementary Table 28: Posterior summaries for Model 2 of firing identification sensitivity in tracked MG MUs across synthetic grid displacement.**

| | 50% | 95% ETI | | Mode | 95% HDI | |
|---|---|---|---|---|---|---|
| | | 2.5% | 97.5% | | low | high |
| $\alpha_{\mu}$ | 0.536 | 0.468 | 0.603 | 0.534 | 0.470 | 0.604 |
| $\alpha_{\phi}$ | 4.260 | 3.619 | 4.961 | 4.240 | 3.615 | 4.956 |
| $\alpha_{\zeta}$ | 0.005 | 0.001 | 0.017 | 0.004 | 0.000 | 0.015 |
| $\alpha_{\kappa}$ | 0.131 | 0.005 | 0.809 | 0.034 | 0.000 | 0.709 |
| $\beta_{\mu,\text{Proposed}}$ | 33.914 | 24.279 | 47.343 | 33.062 | 23.438 | 46.113 |
| $\beta_{\phi,\text{Proposed}}$ | 7.284 | 5.364 | 9.779 | 7.110 | 5.158 | 9.543 |
| $\beta_{\zeta,\text{Proposed}}$ | 165.861 | 50.989 | 841.294 | 119.567 | 25.673 | 637.582 |
| $\beta_{\kappa,\text{Proposed}}$ | 813.029 | 17.770 | 4,6822.670 | -843.483 | 0.470 | 2,4428.118 |
| $\tau_{b0}$ | 0.574 | 0.413 | 0.822 | 0.569 | 0.394 | 0.793 |
| $\tau_{b\text{Proposed}}$ | 0.585 | 0.380 | 0.885 | 0.572 | 0.359 | 0.855 |
| $R_{12}$ | -0.931 | -0.991 | -0.708 | -0.962 | -0.999 | -0.761 |

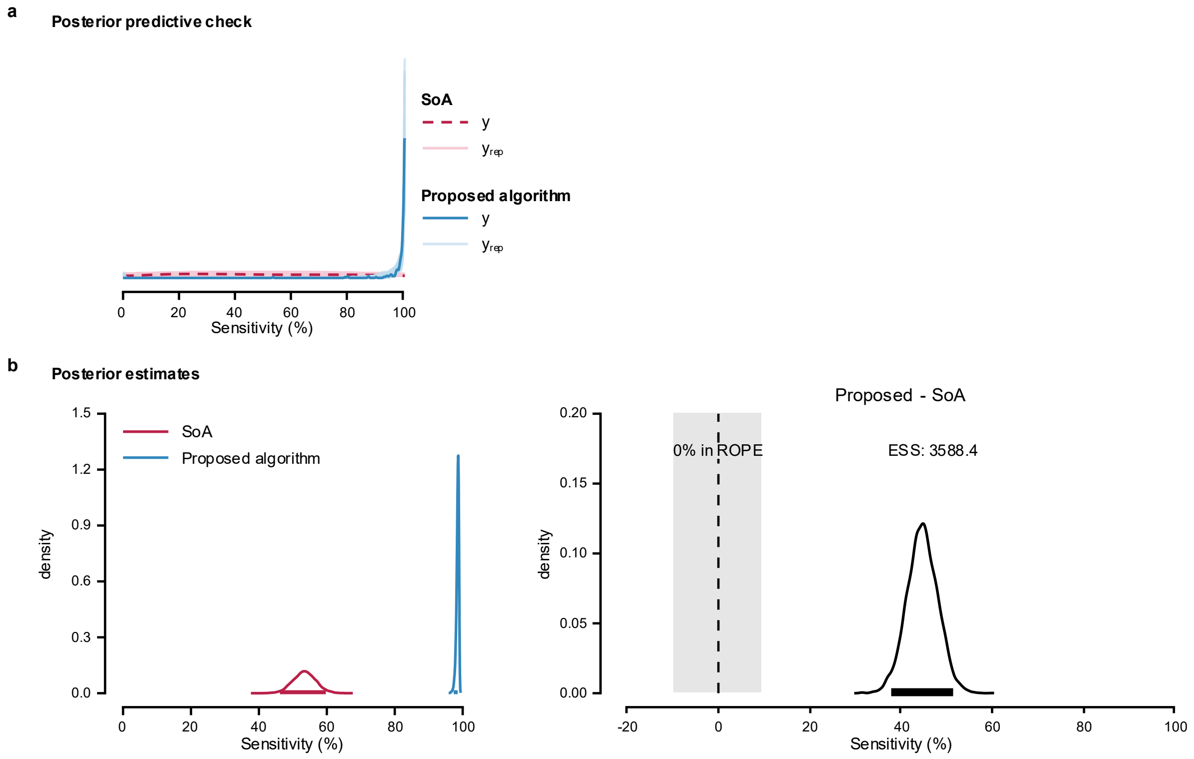


**Supplementary Fig. 21: Posterior predictive check and posterior estimates for Model 2 of firing identification sensitivity in tracked MG MUs across synthetic grid displacement.**

**a**, Posterior predictive check of the model; **b**, Posterior distributions and the difference in means from Bayesian Model 2, used to compare the performance of the proposed algorithm and SoA method.

<u>Model 3</u>

The maximum PSRF across all parameters was 1.002, indicating good convergence, while the minimum ESS was 6,270.581, indicating stable estimates of the bounds of the credible intervals (Supplementary Table 29). Posterior medians, 95% ETIs and 95% HDIs are reported on interpretable scales (Supplementary Table 30). Correlations are reported on their native model/link and correlation scales.

Posterior predictive checks were performed by overlaying model-generated predictions on the observed data (Supplementary Fig. 22a). The posterior distributions (Supplementary Fig. 22b) indicated that the proposed algorithm achieved a higher sensitivity (98.2%, [97.2-98.8%]) than the SoA algorithm (53.3%, [46.7-60.2%]). The posterior distribution of the difference excluded zero, and less than 2.5% of its distribution fell within ROPE.

**Supplementary Table 29: Posterior MCMC diagnostics for Model 3 of firing identification sensitivity in tracked MG MUs across synthetic grid displacement.**

| | PSRF Pt | PSRF UpCI | ESS |
|---|---|---|---|
| $\alpha_{\mu}$ | 1.001 | 1.002 | 6,270.581 |
| $\alpha_{\phi}$ | 1.000 | 1.000 | 2,3045.385 |
| $\alpha_{\zeta}$ | 1.000 | 1.000 | 1,8057.101 |
| $\alpha_{\kappa}$ | 1.000 | 1.001 | 2,5932.481 |
| $\beta_{\mu,\text{Proposed}}$ | 1.000 | 1.001 | 8,603.863 |
| $\beta_{\phi,\text{Proposed}}$ | 1.000 | 1.000 | 2,3671.972 |
| $\beta_{\zeta,\text{Proposed}}$ | 1.000 | 1.000 | 1,8006.556 |
| $\beta_{\kappa,\text{Proposed}}$ | 1.000 | 1.001 | 2,1635.468 |
| $\tau_{b0}$ | 1.000 | 1.001 | 6,798.793 |
| $\tau_{b\text{Proposed}}$ | 1.000 | 1.000 | 8,910.656 |
| $\tau_{c}$ | 1.000 | 1.000 | 6,711.978 |
| $R_{12}$ | 1.000 | 1.000 | 1,2045.112 |

**Supplementary Table 30: Posterior summaries for Model 3 of firing identification sensitivity in tracked MG MUs across synthetic grid displacement.**

| | 50% | 95% ETI | | Mode | 95% HDI | |
|---|---|---|---|---|---|---|
| | | 2.5% | 97.5% | | low | high |
| $\alpha_{\mu}$ | 0.536 | 0.469 | 0.603 | 0.539 | 0.470 | 0.603 |
| $\alpha_{\phi}$ | 4.310 | 3.671 | 5.064 | 4.234 | 3.645 | 5.022 |
| $\alpha_{\zeta}$ | 0.005 | 0.001 | 0.017 | 0.004 | 0.001 | 0.015 |
| $\alpha_{\kappa}$ | 0.138 | 0.006 | 0.811 | 0.035 | 0.000 | 0.715 |
| $\beta_{\mu,\text{Proposed}}$ | 34.080 | 24.216 | 47.391 | 33.097 | 23.828 | 46.774 |
| $\beta_{\phi,\text{Proposed}}$ | 7.259 | 5.311 | 9.717 | 7.078 | 5.252 | 9.638 |
| $\beta_{\zeta,\text{Proposed}}$ | 164.977 | 51.514 | 832.822 | 113.996 | 26.886 | 635.875 |
| $\beta_{\kappa,\text{Proposed}}$ | 764.369 | 16.831 | 4,7034.930 | -796.872 | 0.475 | 2,3437.830 |
| $\tau_{b0}$ | 0.577 | 0.408 | 0.824 | 0.557 | 0.389 | 0.795 |
| $\tau_{b\text{Proposed}}$ | 0.592 | 0.383 | 0.888 | 0.558 | 0.366 | 0.862 |
| $\tau_{c}$ | 0.064 | 0.003 | 0.224 | 0.017 | 0.000 | 0.193 |
| $R_{12}$ | -0.929 | -0.990 | -0.710 | -0.964 | -0.998 | -0.763 |

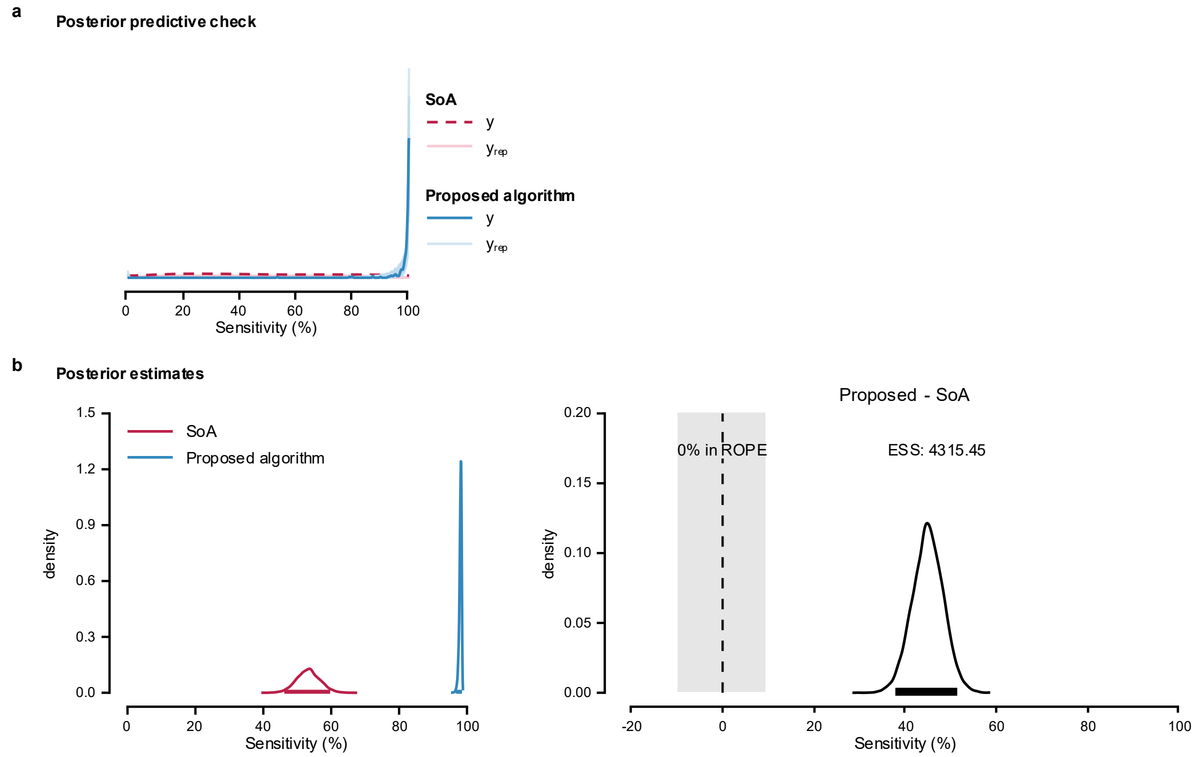


**Supplementary Fig. 22: Posterior predictive check and posterior estimates for Model 3 of firing identification sensitivity in tracked MG MUs across synthetic grid displacement.**

**a**, Posterior predictive check of the model; **b**, Posterior distributions and the difference in means from Bayesian Model 3, used to compare the performance of the proposed algorithm and SoA method.

### MODEL COMPARISON

Model 2 showed the greatest expected predictive accuracy. ELPD diff relative to Model 2 were −0.8 (SE = 0.2) for Model 3 and −24.9 (SE = 5.4) for Model 1, suggesting similar predictive performance for Models 2 and 3 and reduced performance for Model 1.

**Supplementary Note 11. Comparison of firing identification precision in tracked MG MUs across synthetic grid displacement.**

Model implementation, likelihood function, and priors are detailed in Supplementary Note 5. The full dataset contains firing identification precision values for 300 tracked MG MUs (hereafter referred to as 'precision'). The MUs firings were obtained by applying the MU filters calculated using the SoA algorithm[5] and the proposed algorithm to the HDsEMG signals generated with synthetic grid displacement. Distribution of the data is shown in Supplementary Fig. 23.

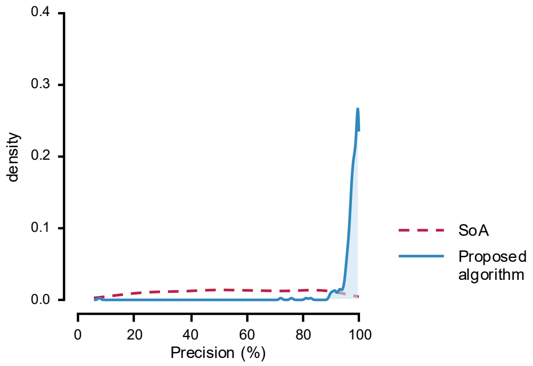


**Supplementary Fig. 23: Distribution of firing identification precision in tracked MG MUs across synthetic grid displacement.**

Precision values were calculated from firing identification obtained by applying MU filters calculated using SoA (dashed line) and the proposed algorithms (solid line) to the HDsEMG signals generated with synthetic grid displacement.

In all models, ZOIB parameters were back transformed into their interpretable component scales: *mu*, *zoi*, and *coi* to probabilities, and *phi* to the positive precision scale. Group-level SDs are reported on their native model/link scale.

Model 1

The maximum PSRF across all parameters was 1.002, indicating good convergence, while the minimum ESS was 7,975.097, indicating stable estimates of the bounds of the credible intervals (Supplementary Table 31). Posterior summaries are reported in Supplementary Table 32 as medians with 95% ETIs and 95% HDIs.

Posterior predictive checks were performed by overlaying model-generated predictions on the observed data (Supplementary Fig. 24a). The posterior distributions (Supplementary Fig. 24b) indicated that the proposed algorithm achieved a higher precision (96.7%, [95.7-97.4 %]) than the SoA algorithm (57.0%, [53.3-60.7%]). The posterior distribution of the difference excluded zero, and less than 2.5% of its distribution fell within ROPE.

**Supplementary Table 31: Posterior MCMC diagnostics for Model 1 of firing identification precision in tracked MG MUs across synthetic grid displacement.**

| | PSRF Pt | PSRF UpCI | ESS |
|---|---|---|---|
| $\alpha_{\mu}$ | 1.000 | 1.001 | 1,4238.646 |
| $\alpha_{\phi}$ | 1.000 | 1.000 | 2,5652.337 |
| $\alpha_{\zeta}$ | 1.001 | 1.002 | 1,7246.424 |
| $\alpha_{\kappa}$ | 1.000 | 1.001 | 2,6810.992 |
| $\beta_{\mu,\text{Proposed}}$ | 1.000 | 1.000 | 2,6089.046 |
| $\beta_{\phi,\text{Proposed}}$ | 1.000 | 1.001 | 2,3659.052 |
| $\beta_{\zeta,\text{Proposed}}$ | 1.001 | 1.002 | 1,7319.186 |
| $\beta_{\kappa,\text{Proposed}}$ | 1.000 | 1.001 | 2,4694.811 |
| $\tau_b$ | 1.000 | 1.000 | 7,975.097 |

**Supplementary Table 32: Posterior summaries for Model 1 of firing identification precision in tracked MG MUs across synthetic grid displacement.**

| | 50% | 95% ETI | | Mode | 95% HDI | |
|---|---|---|---|---|---|---|
| | | **2.5%** | **97.5%** | | **low** | **high** |
| $\alpha_{\mu}$ | 0.573 | 0.536 | 0.610 | 0.572 | 0.536 | 0.610 |
| $\alpha_{\phi}$ | 3.766 | 3.197 | 4.405 | 3.753 | 3.174 | 4.379 |
| $\alpha_{\zeta}$ | 0.006 | 0.001 | 0.018 | 0.004 | 0.001 | 0.016 |
| $\alpha_{\kappa}$ | 0.168 | 0.007 | 0.854 | 0.043 | 0.000 | 0.774 |
| $\beta_{\mu,\text{Proposed}}$ | 19.984 | 16.761 | 23.780 | 19.711 | 16.601 | 23.562 |
| $\beta_{\phi,\text{Proposed}}$ | 5.637 | 4.295 | 7.325 | 5.431 | 4.232 | 7.243 |
| $\beta_{\zeta,\text{Proposed}}$ | 40.759 | 12.805 | 204.304 | 26.014 | 7.296 | 152.293 |
| $\beta_{\kappa,\text{Proposed}}$ | 288.282 | 5.695 | 2,0479.150 | -313.086 | 0.081 | 9,662.875 |
| $\tau_b$ | 0.243 | 0.127 | 0.397 | 0.243 | 0.115 | 0.383 |

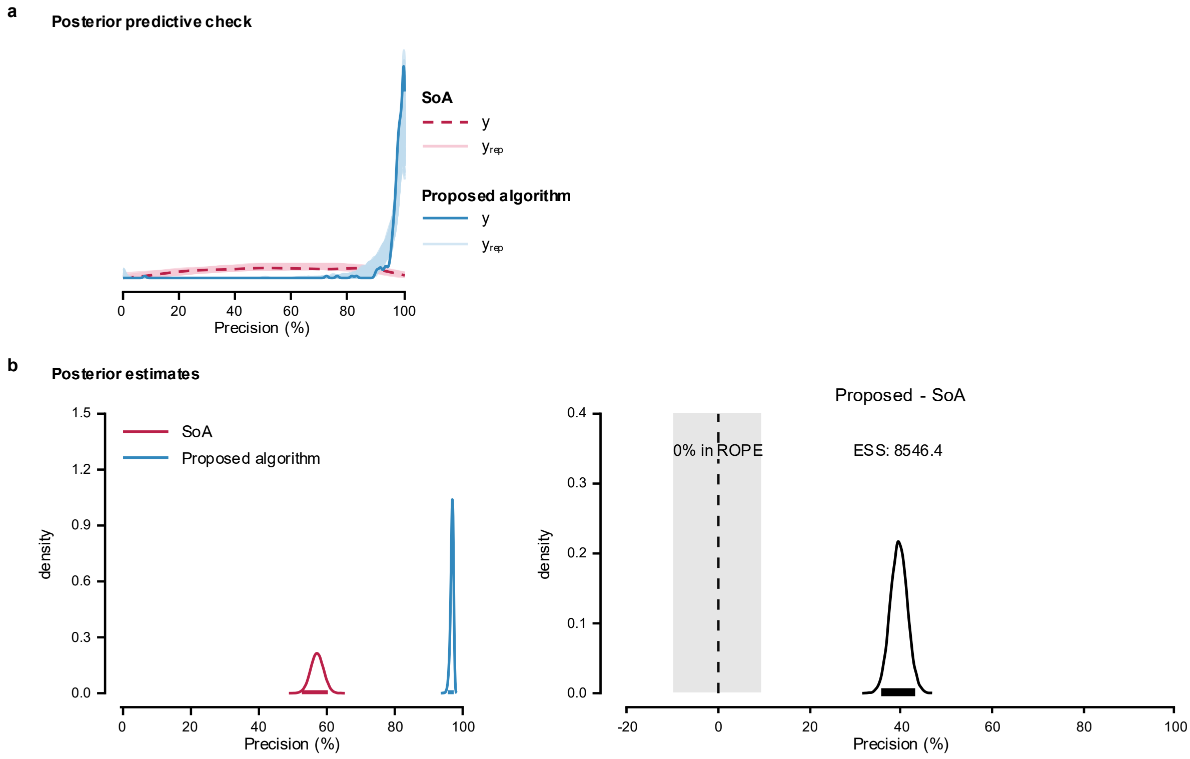


**Supplementary Fig. 24: Posterior predictive check and posterior estimates for Model 1 of firing identification precision in tracked MG MUs across synthetic grid displacement.** **a**, Posterior predictive check of the model; **b**, Posterior distributions and the difference between means estimated using Bayesian Model 1 to compare the performance of the proposed algorithm and SoA method.

<u>Model 2</u>

The maximum PSRF across all parameters was 1.005, indicating good convergence, while the minimum ESS was 4,697.237, indicating stable estimates of the bounds of the credible intervals (Supplementary Table 33). Posterior summaries as medians with 95% ETIs and 95% HDIs are reported in Supplementary Table 34. Group-level correlations are reported on the correlation scale.

Posterior predictive checks were performed by overlaying model-generated predictions on the observed data (Supplementary Fig. 25a). The posterior distributions (Supplementary Fig.

25b) indicated that the proposed algorithm achieved a higher precision (96.5%, [95.5-97.2%]) than the SoA algorithm (60.0%, [54.6-65.6%]). The posterior distribution of the difference excluded zero, and less than 2.5% of its distribution fell within ROPE.

**Supplementary Table 33: Posterior MCMC diagnostics for Model 2 of firing identification precision in tracked MG MUs across synthetic grid displacement.**

| | PSRF Pt | PSRF UpCI | ESS |
|---|---|---|---|
| $\alpha_{\mu}$ | 1.000 | 1.001 | 4,697.237 |
| $\alpha_{\phi}$ | 1.000 | 1.001 | 2,6948.408 |
| $\alpha_{\zeta}$ | 1.000 | 1.000 | 1,0601.011 |
| $\alpha_{\kappa}$ | 1.000 | 1.001 | 1,4062.031 |
| $\beta_{\mu,\text{Proposed}}$ | 1.000 | 1.001 | 5,975.843 |
| $\beta_{\phi,\text{Proposed}}$ | 1.000 | 1.001 | 1,6215.100 |
| $\beta_{\zeta,\text{Proposed}}$ | 1.000 | 1.000 | 1,0766.322 |
| $\beta_{\kappa,\text{Proposed}}$ | 1.000 | 1.001 | 1,5707.576 |
| $\tau_{b0}$ | 1.002 | 1.005 | 5,390.199 |
| $\tau_{b\text{Proposed}}$ | 1.001 | 1.003 | 6,270.154 |
| $R_{12}$ | 1.000 | 1.000 | 7,627.012 |

**Supplementary Table 34: Posterior summaries for Model 2 of firing identification precision in tracked MG MUs across synthetic grid displacement.**

| | 50% | 95% ETI | | Mode | 95% HDI | |
|---|---|---|---|---|---|---|
| | | **2.5%** | **97.5%** | | **low** | **high** |
| $\alpha_{\mu}$ | 0.602 | 0.549 | 0.658 | 0.603 | 0.550 | 0.658 |
| $\alpha_{\phi}$ | 4.527 | 3.873 | 5.269 | 4.519 | 3.859 | 5.249 |
| $\alpha_{\zeta}$ | 0.006 | 0.001 | 0.018 | 0.004 | 0.001 | 0.016 |
| $\alpha_{\kappa}$ | 0.168 | 0.007 | 0.845 | 0.044 | 0.000 | 0.770 |
| $\beta_{\mu,\text{Proposed}}$ | 16.691 | 12.324 | 22.223 | 16.650 | 12.085 | 21.884 |
| $\beta_{\phi,\text{Proposed}}$ | 4.992 | 3.844 | 6.421 | 4.991 | 3.765 | 6.313 |
| $\beta_{\zeta,\text{Proposed}}$ | 41.291 | 12.603 | 200.589 | 27.793 | 7.144 | 149.862 |
| $\beta_{\kappa,\text{Proposed}}$ | 291.746 | 5.888 | 2,1124.880 | -312.421 | 0.143 | 1,0045.306 |
| $\tau_{b0}$ | 0.472 | 0.329 | 0.685 | 0.447 | 0.321 | 0.670 |
| $\tau_{b\text{Proposed}}$ | 0.547 | 0.364 | 0.809 | 0.540 | 0.359 | 0.799 |
| $R_{12}$ | -0.896 | -0.983 | -0.644 | -0.936 | -0.994 | -0.698 |

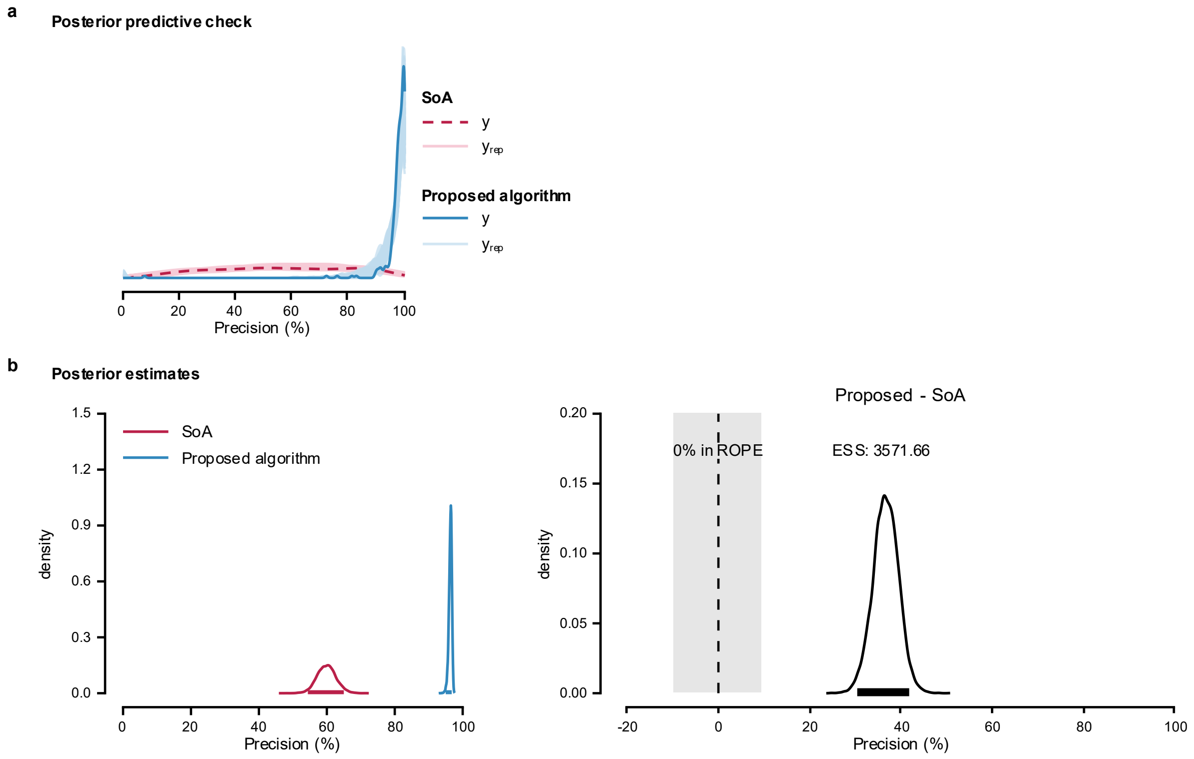


**Supplementary Fig. 25: Posterior predictive check and posterior estimates for Model 2 of firing identification precision in tracked MG MUs across synthetic grid displacement.** **a**, Posterior predictive check of the model; **b**, Posterior distributions and the difference between means estimated using Bayesian Model 2 to compare the performance of the proposed algorithm and SoA method.

<u>Model 3</u>

The maximum PSRF across all parameters was 1.003, indicating good convergence, while the minimum ESS was 3,675.855, indicating stable estimates of the bounds of the credible intervals (Supplementary Table 35). Posterior summaries are reported as medians with 95% ETIs and 95% HDIs (Supplementary Table 36). Group-level correlations are reported on the correlation scale.

Posterior predictive checks were performed by overlaying model-generated predictions on the observed data (Supplementary Fig. 26a). The posterior distributions (Supplementary Fig.

26b) indicated that the proposed algorithm achieved a higher precision (95.2%, [94.4-95.8%]) than the SoA algorithm (29.5%, [26.0-33.1%]). The posterior distribution of the difference excluded zero, and less than 2.5% of its distribution fell within ROPE.

**Supplementary Table 35: Posterior MCMC diagnostics for Model 3 of firing identification precision in tracked MG MUs across synthetic grid displacement.**

| | PSRF Pt | PSRF UpCI | ESS |
|---|---|---|---|
| $\alpha_{\mu}$ | 1.001 | 1.003 | 6,707.515 |
| $\alpha_{\phi}$ | 1.000 | 1.000 | 1,3539.405 |
| $\alpha_{\zeta}$ | 1.000 | 1.001 | 1,4119.397 |
| $\alpha_{\kappa}$ | 1.000 | 1.000 | 2,1987.004 |
| $\beta_{\mu,\text{Proposed}}$ | 1.001 | 1.002 | 7,261.005 |
| $\beta_{\phi,\text{Proposed}}$ | 1.000 | 1.001 | 3,675.855 |
| $\beta_{\zeta,\text{Proposed}}$ | 1.000 | 1.001 | 1,4193.201 |
| $\beta_{\kappa,\text{Proposed}}$ | 1.000 | 1.000 | 1,9256.450 |
| $\tau_{b0}$ | 1.000 | 1.001 | 6,586.077 |
| $\tau_{b\text{Proposed}}$ | 1.000 | 1.000 | 6,752.279 |
| $\tau_{c}$ | 1.000 | 1.001 | 4,099.645 |
| $R_{12}$ | 1.001 | 1.003 | 5,109.176 |

**Supplementary Table 36: Posterior summaries for Model 3 of firing identification precision in tracked MG MUs across synthetic grid displacement.**

| | 50% | 95% ETI | | Mode | 95% HDI | |
|---|---|---|---|---|---|---|
| | | 2.5% | 97.5% | | low | high |
| $\alpha_{\mu}$ | 0.605 | 0.542 | 0.666 | 0.604 | 0.545 | 0.669 |
| $\alpha_{\phi}$ | 5.556 | 4.550 | 6.774 | 5.475 | 4.475 | 6.687 |
| $\alpha_{\zeta}$ | 0.006 | 0.001 | 0.018 | 0.004 | 0.000 | 0.016 |
| $\alpha_{\kappa}$ | 0.169 | 0.007 | 0.856 | 0.043 | 0.001 | 0.775 |
| $\beta_{\mu,\text{Proposed}}$ | 24.856 | 17.430 | 35.033 | 23.991 | 16.691 | 33.977 |
| $\beta_{\phi,\text{Proposed}}$ | 17.705 | 10.757 | 30.779 | 16.286 | 9.877 | 28.974 |
| $\beta_{\zeta,\text{Proposed}}$ | 40.809 | 12.599 | 201.477 | 25.376 | 6.692 | 152.015 |
| $\beta_{\kappa,\text{Proposed}}$ | 287.547 | 5.805 | 1,9533.890 | -303.170 | 0.149 | 9,755.657 |
| $\tau_{b0}$ | 0.505 | 0.338 | 0.749 | 0.487 | 0.322 | 0.722 |
| $\tau_{b\text{Proposed}}$ | 0.712 | 0.514 | 1.017 | 0.692 | 0.499 | 0.989 |
| $\tau_{c}$ | 0.632 | 0.533 | 0.734 | 0.625 | 0.529 | 0.730 |
| $R_{12}$ | -0.773 | -0.933 | -0.421 | -0.825 | -0.961 | -0.482 |

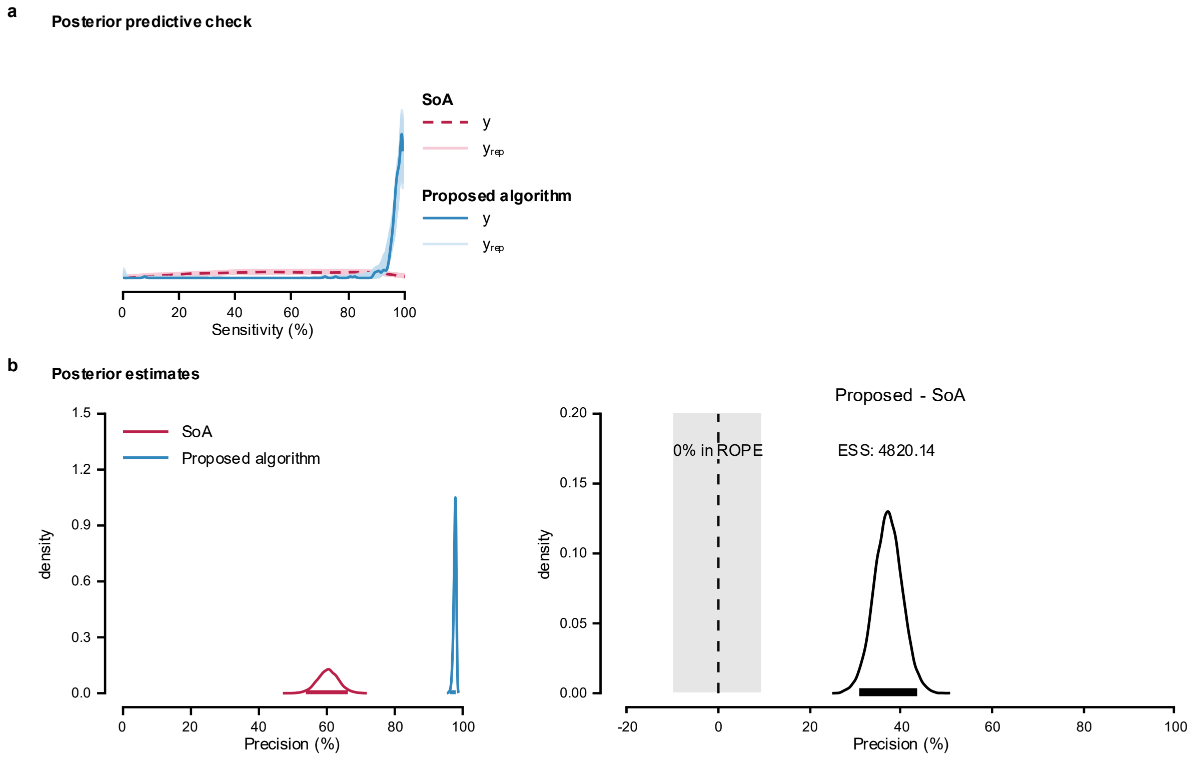


**Supplementary Fig. 26: Posterior predictive check and posterior estimates for Model 3 of firing identification precision in tracked MG MUs across synthetic grid displacement.** **a**, Posterior predictive check of the model; **b**, Posterior distributions and the difference between means estimated using Bayesian Model 1 to compare the performance of the proposed algorithm and SoA method.

### MODEL COMPARISON

Model 2 had the highest expected predictive accuracy. Relative to Model 2, the ELPD diff were −7.5 (SE = 15.1) for Model 3 and −17.6 (SE = 14.6) for Model 1. As both differences were small relative to their uncertainty, the models showed no clear differences in out-of-sample predictive performance, although the point estimates favored Model 2.

**Supplementary Note 12. Comparison of firing identification PNR for tracked TA MUs across testing visits.**

Model implementation, likelihood function, and priors are detailed in Supplementary Note 5. The full dataset contains firing identification PNR values for 119 tracked MUs in TA muscle (Supplementary Fig. 27). The MUs firings were obtained by applying the MU filters calculated using the SoA algorithm[5] and the proposed algorithm to the HDsEMG signals obtained in the second testing visit.

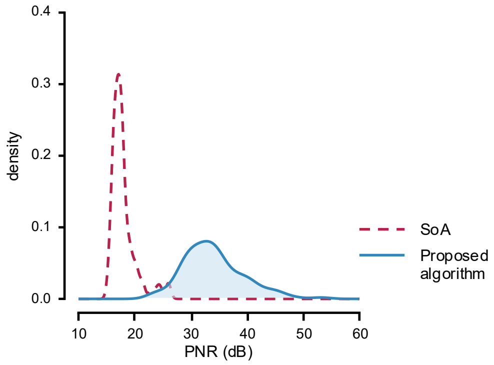


**Supplementary Fig. 27: Distribution of firing identification PNR in tracked TA MUs across testing visits.**

PNR values were calculated from firing identification obtained by applying MU filters calculated using SoA (dashed line) and the proposed algorithms (solid line) to the HDsEMG signals recorded in the second testing visit.

In all models, the lognormal mean and sigma effects were exponentiated before the posterior summaries were calculated. Group-level SDs are reported on their native model or link scales.

<u>Model 1</u>

The maximum PSRF across all parameters was 1.002, indicating good convergence, while the minimum ESS was 4,130.150, indicating stable estimates of the bounds of the credible intervals (Supplementary Table 37). Median of the estimate value in the original scale of PNR, the limits of the 95% ETI, and the limits of the 95% HDI are reported in Supplementary Table 38.

**Supplementary Table 37: Posterior MCMC diagnostics for Model 1 of firing identification PNR in tracked TA MUs across testing visits.**

| | PSRF Pt | PSRF UpCI | ESS |
|---|---|---|---|
| $\alpha_{\mu}$ | 1.001 | 1.002 | 4,130.150 |
| $\alpha_{\sigma}$ | 1.000 | 1.000 | 1,2408.010 |
| $\beta_{\mu,\text{Proposed}}$ | 1.000 | 1.000 | 2,0473.389 |
| $\beta_{\sigma,\text{Proposed}}$ | 1.000 | 1.000 | 1,4264.345 |
| $\tau_{\mu}$ | 1.001 | 1.001 | 5,003.463 |

**Supplementary Table 38: Posterior summaries for Model 1 of firing identification PNR in tracked TA MUs across testing visits.**

| | 50% | 95% ETI | | Mode | 95% HDI | |
|---|---|---|---|---|---|---|
| | | 2.5% | 97.5% | | low | high |
| $\alpha_{\mu}$ | 17.948 | 17.013 | 18.959 | 17.973 | 16.987 | 18.926 |
| $\alpha_{\sigma}$ | 1.871 | 1.811 | 1.935 | 1.867 | 1.811 | 1.935 |
| $\beta_{\mu,\text{Proposed}}$ | 0.080 | 0.069 | 0.094 | 0.080 | 0.068 | 0.093 |
| $\beta_{\sigma,\text{Proposed}}$ | 2.063 | 1.665 | 2.541 | 2.030 | 1.638 | 2.509 |
| $\tau_{\mu}$ | 0.083 | 0.051 | 0.142 | 0.074 | 0.047 | 0.133 |

Posterior predictive checks were performed by overlaying model-generated predictions on the observed data (Supplementary Fig. 28a). The posterior distributions (Supplementary Fig. 28b) indicated that the proposed algorithm achieved a higher PNR (34.1 dB, [32.1-36.2 dB]) than the SoA algorithm (18.0 dB, [17.1-19.0 dB]). The posterior distribution of the difference excluded zero, and less than 2.5% of its distribution fell within ROPE.

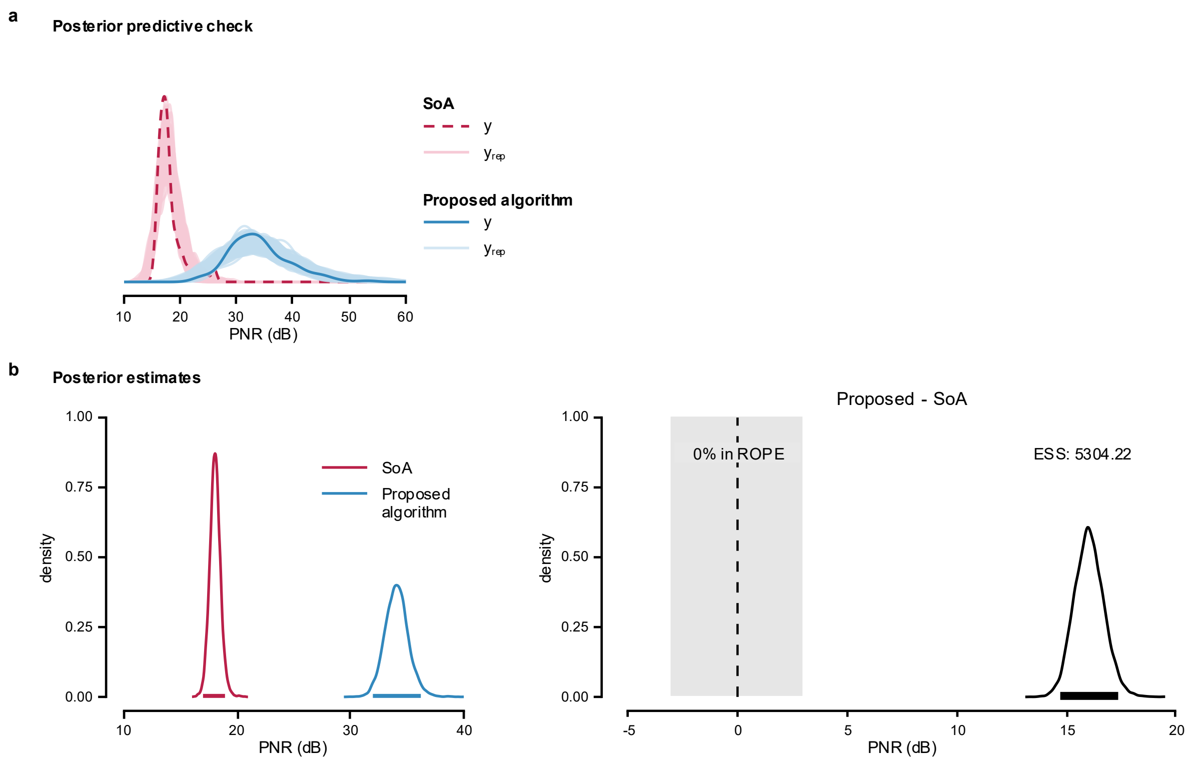


**Supplementary Fig. 28: Posterior predictive check and posterior estimates for Model 1 of firing identification PNR in tracked TA MUs across testing visits.**

**a**, Posterior predictive check of the model; **b**, Posterior distributions and the difference between means estimated using Bayesian Model 1 to compare the performance of the proposed algorithm and SoA method.

Model 2

The maximum PSRF across all parameters was 1.003, indicating good convergence, while the minimum ESS was 4,523.017, indicating stable estimates of the bounds of the credible intervals (Supplementary Table 39). Posterior estimates are summarized by the median, 95% ETIs and 95% HDIs (Supplementary Table 40). Group-level correlations were retained on their native scales.

Posterior predictive checks were performed by overlaying model-generated predictions on the observed data (Supplementary Fig. 29a). The posterior distributions (Supplementary Fig. 29b) indicated that the proposed algorithm achieved a higher PNR (33.2 dB, [31.0-35.0 dB]) than the SoA algorithm (18.3 dB, [17.2-19.6 dB]). The posterior distribution of the difference excluded zero, and less than 2.5% of its distribution fell within ROPE.

Model 3

The maximum PSRF across all parameters was 1.006, indicating good convergence, while the minimum ESS was 4,930.471, indicating stable estimates of the bounds of the credible intervals (Supplementary Table 41). Correlations were summarized on their native scales and values are reported as medians with 95% ETIs and 95% HDIs (Supplementary Table 42).

Posterior predictive checks were performed by overlaying model-generated predictions on the observed data (Supplementary Fig. 30a). The posterior distributions (Supplementary Fig. 30b) indicated that the proposed algorithm achieved a higher PNR (33.1 dB, [31.0-35.0 dB]) than the SoA algorithm (18.3 dB, [17.3-19.5 dB]). The posterior distribution of the difference excluded zero, and less than 2.5% of its distribution fell within ROPE.

**Supplementary Table 39: Posterior MCMC diagnostics for Model 2 of firing identification PNR in tracked TA MUs across testing visits.**

| | PSRF Pt | PSRF UpCI | ESS |
|---|---|---|---|
| $\alpha_{\mu}$ | 1.000 | 1.001 | 4,523.017 |
| $\alpha_{\sigma}$ | 1.000 | 1.000 | 1,7991.923 |
| $\beta_{\mu,\text{Proposed}}$ | 1.000 | 1.000 | 5,467.359 |
| $\beta_{\sigma,\text{Proposed}}$ | 1.000 | 1.000 | 1,9290.546 |
| $\tau_{b0}$ | 1.001 | 1.003 | 5,921.262 |
| $\tau_{b\text{Proposed}}$ | 1.000 | 1.001 | 7,009.741 |
| $R_{12}$ | 1.000 | 1.001 | 8,687.863 |

**Supplementary Table 40: Posterior summaries for Model 2 of firing identification PNR in tracked TA MUs across testing visits.**

| | 50% | 95% ETI | | Mode | 95% HDI | |
|---|---|---|---|---|---|---|
| | | 2.5% | 97.5% | | low | high |
| $\alpha_{\mu}$ | 18.216 | 17.183 | 19.487 | 18.146 | 17.115 | 19.418 |
| $\alpha_{\sigma}$ | 0.074 | 0.065 | 0.085 | 0.073 | 0.065 | 0.085 |
| $\beta_{\mu,\text{Proposed}}$ | 1.802 | 1.636 | 1.947 | 1.798 | 1.646 | 1.955 |
| $\beta_{\sigma,\text{Proposed}}$ | 1.917 | 1.600 | 2.288 | 1.914 | 1.585 | 2.266 |
| $\tau_{b0}$ | 0.096 | 0.064 | 0.156 | 0.088 | 0.059 | 0.147 |
| $\tau_{b\text{Proposed}}$ | 0.122 | 0.075 | 0.208 | 0.117 | 0.069 | 0.195 |
| $R_{12}$ | -0.794 | -0.965 | -0.297 | -0.866 | -0.987 | -0.395 |

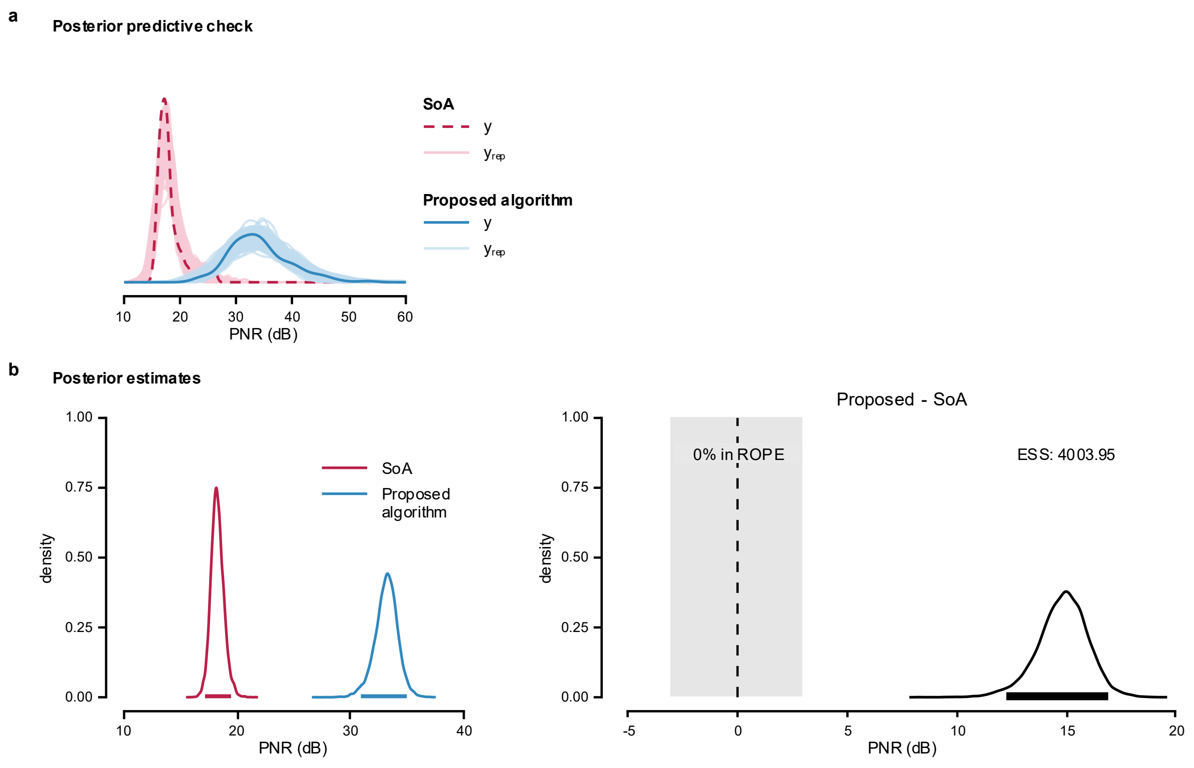


**Supplementary Fig. 29: Posterior predictive check and posterior estimates for Model 2 of firing identification PNR in tracked TA MUs across testing visits.**

**a**, Posterior predictive check of the model; **b**, Posterior distributions and the difference in means from Bayesian Model 2, used to compare the performance of the proposed algorithm and SoA method.

**Supplementary Table 41: Posterior MCMC diagnostics for Model 3 of firing identification PNR in tracked TA MUs across testing visits.**

| | PSRF Pt | PSRF UpCI | ESS |
|---|---|---|---|
| $\alpha_{\mu}$ | 1.000 | 1.001 | 4,930.471 |
| $\alpha_{\sigma}$ | 1.000 | 1.000 | 2,0872.561 |
| $\beta_{\mu,\text{Proposed}}$ | 1.000 | 1.001 | 6,970.288 |
| $\beta_{\sigma,\text{Proposed}}$ | 1.000 | 1.000 | 2,5687.106 |
| $\tau_{b0}$ | 1.000 | 1.002 | 7,382.935 |
| $\tau_{b\text{Proposed}}$ | 1.001 | 1.003 | 7,809.601 |
| $\tau_{c}$ | 1.002 | 1.006 | 8,923.323 |
| $R_{12}$ | 1.000 | 1.000 | 9,823.605 |

**Supplementary Table 42: Posterior summaries for Model 3 of firing identification PNR in tracked TA MUs across testing visits.**

| | 50% | 95% ETI | | Mode | 95% HDI | |
|---|---|---|---|---|---|---|
| | | 2.5% | 97.5% | | low | high |
| $\alpha_{\mu}$ | 18.229 | 17.211 | 19.462 | 18.170 | 17.176 | 19.426 |
| $\alpha_{\sigma}$ | 0.074 | 0.064 | 0.085 | 0.073 | 0.064 | 0.085 |
| $\beta_{\mu,\text{Proposed}}$ | 1.801 | 1.637 | 1.941 | 1.802 | 1.647 | 1.950 |
| $\beta_{\sigma,\text{Proposed}}$ | 1.929 | 1.601 | 2.316 | 1.912 | 1.577 | 2.289 |
| $\tau_{b0}$ | 0.096 | 0.064 | 0.157 | 0.088 | 0.061 | 0.149 |
| $\tau_{b\text{Proposed}}$ | 0.122 | 0.075 | 0.210 | 0.114 | 0.069 | 0.199 |
| $\tau_{c}$ | 0.010 | 0.000 | 0.030 | 0.003 | 0.000 | 0.026 |
| $R_{12}$ | -0.792 | -0.965 | -0.278 | -0.880 | -0.986 | -0.376 |

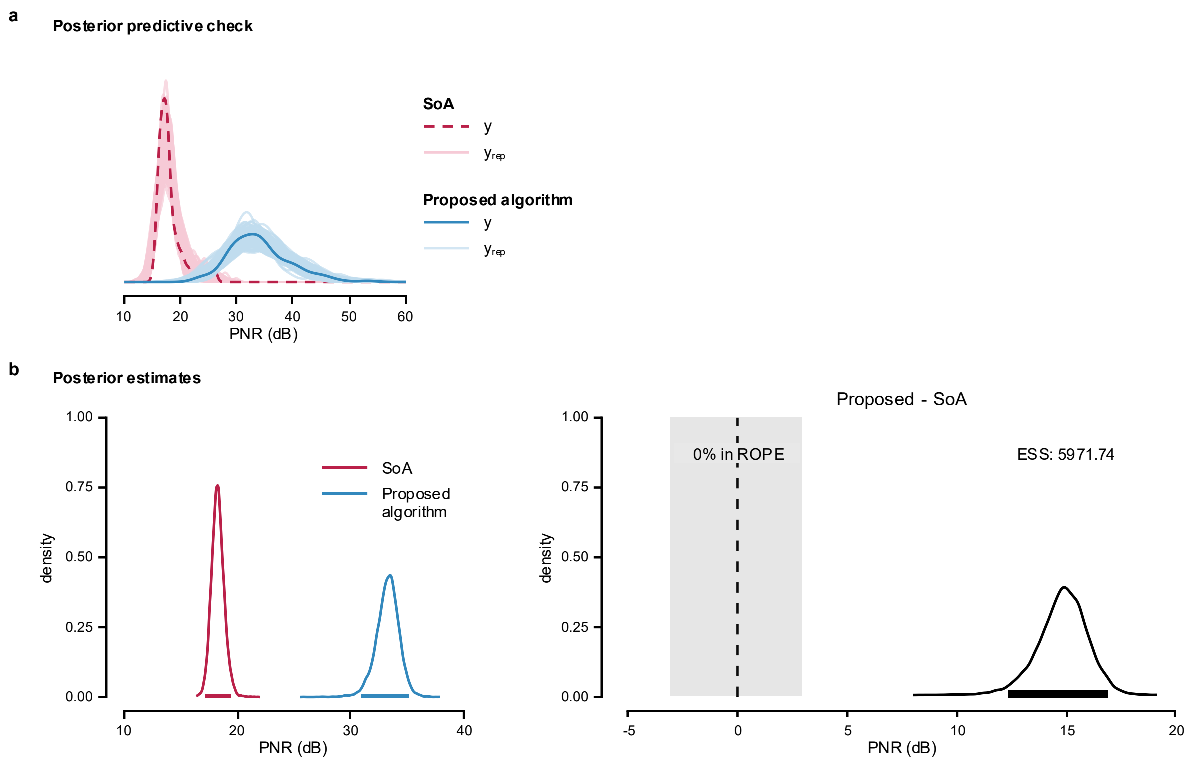


**Supplementary Fig. 30: Posterior predictive check and posterior estimates for Model 3 of firing identification PNR in tracked TA MUs across testing visits.**

**a**, Posterior predictive check of the model; **b**, Posterior distributions and the difference in means from Bayesian Model 3, used to compare the performance of the proposed algorithm and SoA method.

### MODEL COMPARISON

Model 2 showed the greatest expected predictive accuracy. ELPD diff relative to Model 2 were −0.9 (SE = 0.3) for Model 3 and −22.5 (SE = 7.6) for Model 1, suggesting similar predictive performance for Models 2 and 3 and reduced performance for Model 1.

**Supplementary Note 13. Comparison of firing identification sensitivity in tracked TA MUs across testing visits.**

Model implementation, likelihood function, and priors are detailed in Supplementary Note 5. The full dataset contains firing identification sensitivity values for 119 tracked TA MUs (hereafter referred to as 'sensitivity'). The MUs firings were obtained by applying the MU filters calculated using the SoA algorithm[5] and the proposed algorithm to the HDsEMG signals obtained in the second testing visit. Distribution of the data is shown in Supplementary Fig. 31.

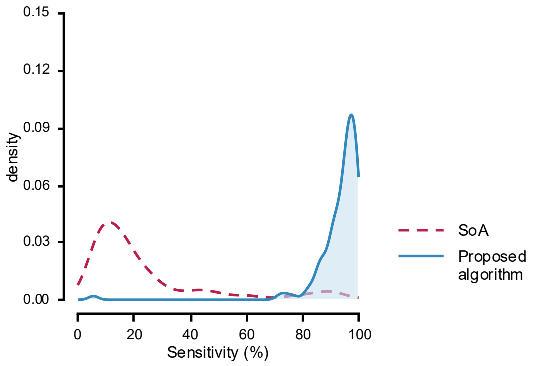


**Supplementary Fig. 31: Distribution of firing identification sensitivity in tracked TA MUs across testing visits.**

Sensitivity values were calculated from firing identification obtained by applying MU filters calculated using SoA (dashed line) and the proposed algorithms (solid line) to the HDsEMG signals generated with synthetic grid displacement.

In all models, ZOIB parameters were back transformed into their interpretable component scales: *mu*, *zoi*, and *coi* to probabilities, and *phi* to the positive precision scale. Group-level SDs are reported on their native model/link scale.

Model 1

The maximum PSRF across all parameters was 1.003, indicating good convergence, while the minimum ESS was 4,215.681, indicating stable estimates of the bounds of the credible intervals (Supplementary Table 43). Posterior summaries are reported in Supplementary Table 44 as medians with 95% ETIs and the 95% HDIs.

Posterior predictive checks were performed by overlaying model-generated predictions on the observed data (Supplementary Fig. 32a). The posterior distributions (Supplementary Fig. 32b) indicated that the proposed algorithm achieved a higher sensitivity (92.0%, [87.5-95.1%]) than the SoA algorithm (22.5%, [14.6-31.6%]). The posterior distribution of the difference excluded zero, and less than 2.5% of its distribution fell within ROPE.

**Supplementary Table 43: Posterior MCMC diagnostics for Model 1 of firing identification sensitivity in tracked TA MUs across testing visits.**

| | PSRF Pt | PSRF UpCI | ESS |
|---|---|---|---|
| $\alpha_{\mu}$ | 1.001 | 1.003 | 4,215.681 |
| $\alpha_{\phi}$ | 1.000 | 1.001 | 1,5084.716 |
| $\alpha_{\zeta}$ | 1.000 | 1.001 | 1,0663.543 |
| $\alpha_{\kappa}$ | 1.000 | 1.000 | 1,4091.757 |
| $\beta_{\mu,\text{Proposed}}$ | 1.000 | 1.001 | 1,5437.385 |
| $\beta_{\phi,\text{Proposed}}$ | 1.000 | 1.001 | 1,6987.396 |
| $\beta_{\zeta,\text{Proposed}}$ | 1.000 | 1.001 | 1,1797.238 |
| $\beta_{\kappa,\text{Proposed}}$ | 1.000 | 1.000 | 1,5458.274 |
| $\tau_{b}$ | 1.000 | 1.001 | 5,863.542 |

**Supplementary Table 44: Posterior summaries for Model 1 of firing identification sensitivity in tracked TA MUs across testing visits.**

| | 50% | 95% ETI | | Mode | 95% HDI | |
|---|---|---|---|---|---|---|
| | | **2.5%** | **97.5%** | | **low** | **high** |
| $\alpha_{\mu}$ | 0.223 | 0.144 | 0.318 | 0.223 | 0.141 | 0.312 |
| $\alpha_{\phi}$ | 9.623 | 7.063 | 12.692 | 9.481 | 6.984 | 12.578 |
| $\alpha_{\zeta}$ | 0.014 | 0.003 | 0.042 | 0.009 | 0.001 | 0.036 |
| $\alpha_{\kappa}$ | 0.248 | 0.011 | 0.900 | 0.054 | 0.001 | 0.845 |
| $\beta_{\mu,\text{Proposed}}$ | 41.576 | 31.925 | 53.373 | 41.717 | 31.163 | 52.277 |
| $\beta_{\phi,\text{Proposed}}$ | 1.023 | 0.670 | 1.556 | 0.968 | 0.644 | 1.509 |
| $\beta_{\zeta,\text{Proposed}}$ | 8.266 | 2.200 | 46.386 | 5.258 | 1.005 | 34.951 |
| $\beta_{\kappa,\text{Proposed}}$ | 48.988 | 0.818 | 4,662.301 | -59.101 | 0.009 | 2,232.798 |
| $\tau_b$ | 0.804 | 0.522 | 1.351 | 0.755 | 0.475 | 1.263 |

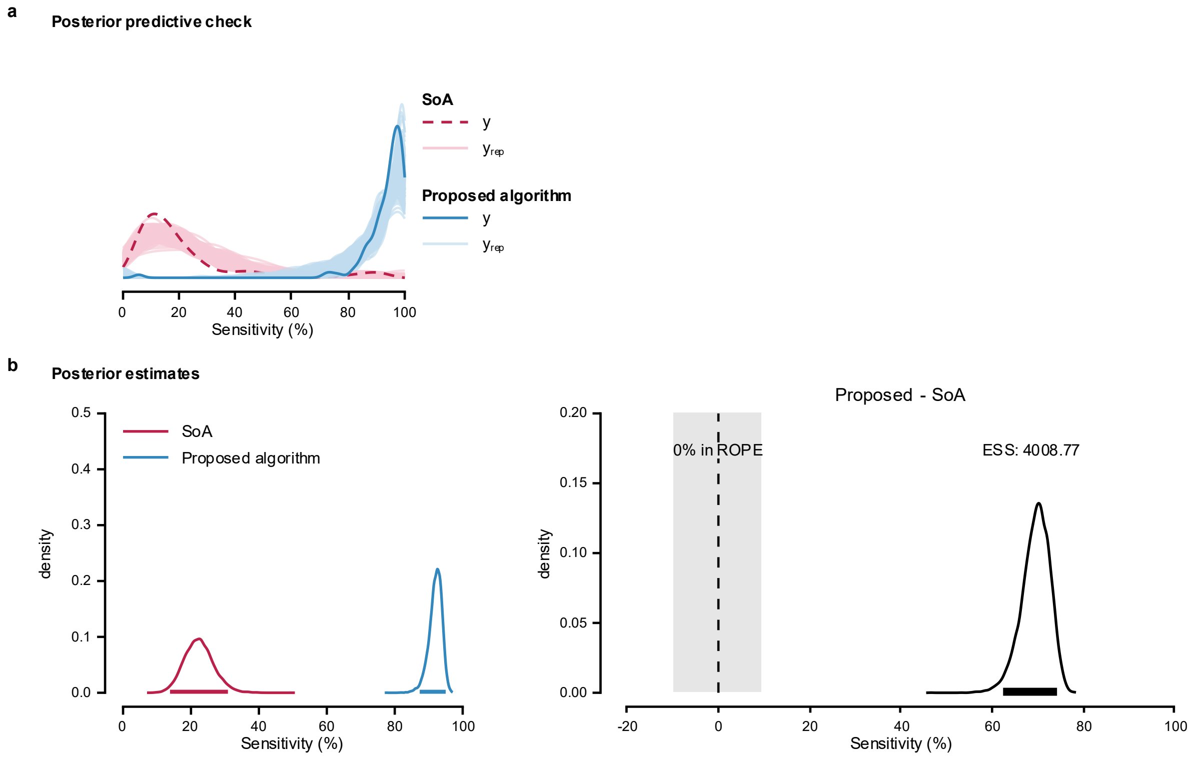


**Supplementary Fig. 32: Posterior predictive check and posterior estimates for Model 1 of firing identification sensitivity in tracked TA MUs across testing visits.**

**a**, Posterior predictive check of the model; **b**, Posterior distributions and the difference between means estimated using Bayesian Model 1 to compare the performance of the proposed algorithm and SoA method.

Model 2

The maximum PSRF across all parameters was 1.001, indicating good convergence, while the minimum ESS was 5,341.062, indicating stable estimates of the bounds of the credible intervals (Supplementary Table 45). Medians with 95% ETIs and 95% HDIs are reported in Supplementary Table 46. Group-level correlations are reported on the correlation scale.

Posterior predictive checks were performed by overlaying model-generated predictions on the observed data (Supplementary Fig. 33a). The posterior distributions (Supplementary Fig. 33b) indicated that the proposed algorithm achieved a higher sensitivity (91.3%, [88.1-

93.7%]) than the SoA algorithm (23.5%, [15.1-33.5%]). The posterior distribution of the difference excluded zero, and less than 2.5% of its distribution fell within ROPE.

**Supplementary Table 45: Posterior MCMC diagnostics for Model 2 of firing identification sensitivity in tracked TA MUs across testing visits.**

| | PSRF Pt | PSRF UpCI | ESS |
|---|---|---|---|
| $\alpha_{\mu}$ | 1.000 | 1.001 | 5,341.062 |
| $\alpha_{\phi}$ | 1.000 | 1.000 | 2,3605.135 |
| $\alpha_{\zeta}$ | 1.000 | 1.001 | 1,5785.997 |
| $\alpha_{\kappa}$ | 1.000 | 1.000 | 2,0488.653 |
| $\beta_{\mu,\text{Proposed}}$ | 1.000 | 1.001 | 6,985.508 |
| $\beta_{\phi,\text{Proposed}}$ | 1.000 | 1.000 | 2,4487.073 |
| $\beta_{\zeta,\text{Proposed}}$ | 1.000 | 1.001 | 1,6925.057 |
| $\beta_{\kappa,\text{Proposed}}$ | 1.000 | 1.000 | 2,0792.586 |
| $\tau_{b0}$ | 1.000 | 1.000 | 8,042.371 |
| $\tau_{b\text{Proposed}}$ | 1.000 | 1.000 | 1,0041.826 |
| $R_{12}$ | 1.000 | 1.001 | 1,1376.206 |

**Supplementary Table 46: Posterior summaries for Model 2 of firing identification sensitivity in tracked TA MUs across testing visits.**

| | 50% | 95% ETI | | Mode | 95% HDI | |
|---|---|---|---|---|---|---|
| | | 2.5% | 97.5% | | low | high |
| $\alpha_{\mu}$ | 0.230 | 0.149 | 0.335 | 0.229 | 0.147 | 0.332 |
| $\alpha_{\phi}$ | 11.018 | 8.326 | 14.181 | 10.979 | 8.147 | 13.941 |
| $\alpha_{\zeta}$ | 0.013 | 0.003 | 0.042 | 0.009 | 0.001 | 0.036 |
| $\alpha_{\kappa}$ | 0.248 | 0.012 | 0.905 | 0.056 | 0.001 | 0.850 |
| $\beta_{\mu,\text{Proposed}}$ | 35.187 | 20.475 | 59.348 | 33.674 | 18.915 | 56.528 |
| $\beta_{\phi,\text{Proposed}}$ | 0.929 | 0.628 | 1.377 | 0.883 | 0.595 | 1.324 |
| $\beta_{\zeta,\text{Proposed}}$ | 8.332 | 2.159 | 47.273 | 4.762 | 1.042 | 35.144 |
| $\beta_{\kappa,\text{Proposed}}$ | 49.394 | 0.784 | 4,082.842 | -57.578 | 0.004 | 1,963.511 |
| $\tau_{b0}$ | 0.842 | 0.583 | 1.302 | 0.800 | 0.546 | 1.240 |
| $\tau_{b\text{Proposed}}$ | 0.752 | 0.461 | 1.225 | 0.703 | 0.433 | 1.175 |
| $R_{12}$ | -0.920 | -0.992 | -0.579 | -0.967 | -0.999 | -0.657 |

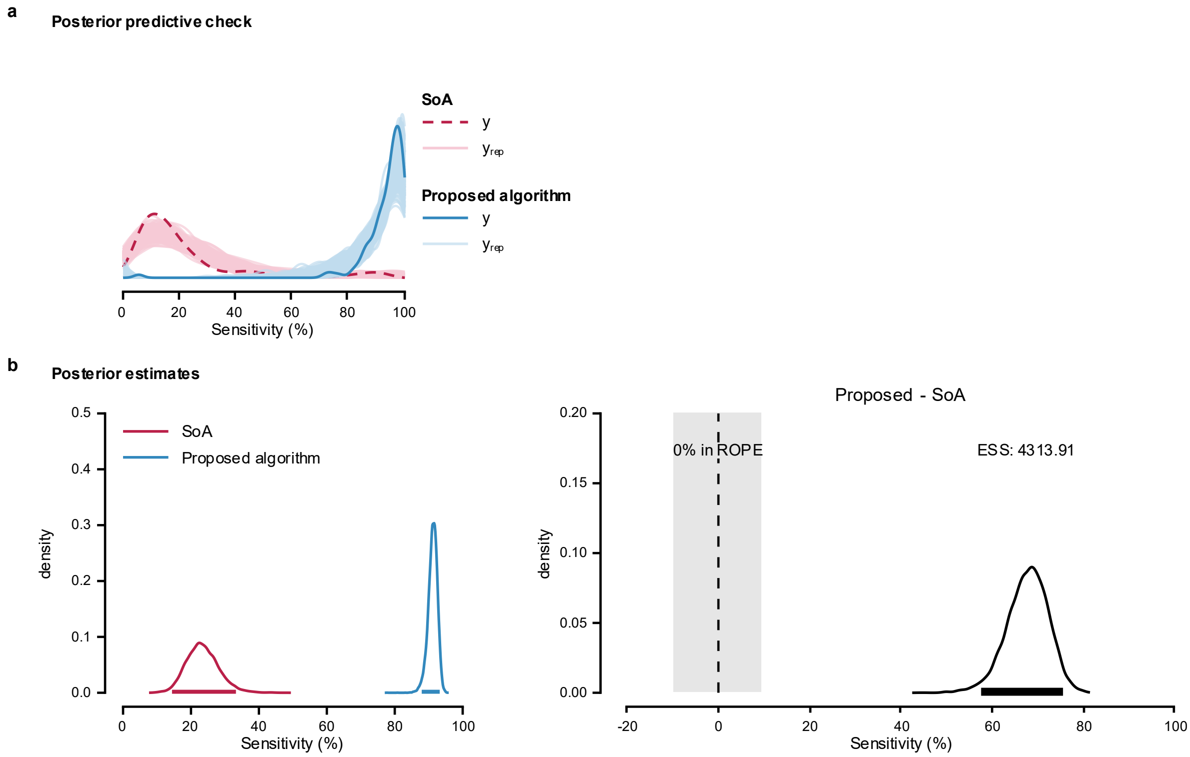


**Supplementary Fig. 33: Posterior predictive check and posterior estimates for Model 2 of firing identification sensitivity in tracked TA MUs across testing visits.**

**a**, Posterior predictive check of the model; **b**, Posterior distributions and the difference in means from Bayesian Model 2, used to compare the performance of the proposed algorithm and SoA method.

<u>Model 3</u>

The maximum PSRF across all parameters was 1.005, indicating good convergence, while the minimum ESS was 3,962.626, indicating stable estimates of the bounds of the credible intervals (Supplementary Table 47). Posterior parameter summaries are given as medians with 95% ETIs and 95% HDIs (Supplementary Table 48). Correlations were summarized without transformation.

Posterior predictive checks were performed by overlaying model-generated predictions on the observed data (Supplementary Fig. 34a). The posterior distributions (Supplementary Fig.

34b) indicated that the proposed algorithm achieved a higher sensitivity (91.3%, [88.1-93.7%]) than the SoA algorithm (23.4%, [14.7-33.6%]). The posterior distribution of the difference excluded zero, and less than 2.5% of its distribution fell within ROPE.

**Supplementary Table 47: Posterior MCMC diagnostics for Model 3 of firing identification sensitivity in tracked TA MUs across testing visits.**

| | PSRF Pt | PSRF UpCI | ESS |
|---|---|---|---|
| $\alpha_{\mu}$ | 1.002 | 1.005 | 3,962.626 |
| $\alpha_{\phi}$ | 1.000 | 1.000 | 2,9278.024 |
| $\alpha_{\zeta}$ | 1.000 | 1.000 | 1,6426.892 |
| $\alpha_{\kappa}$ | 1.000 | 1.000 | 2,5438.194 |
| $\beta_{\mu,\text{Proposed}}$ | 1.001 | 1.003 | 5,509.277 |
| $\beta_{\phi,\text{Proposed}}$ | 1.000 | 1.000 | 3,1206.007 |
| $\beta_{\zeta,\text{Proposed}}$ | 1.000 | 1.000 | 1,7555.071 |
| $\beta_{\kappa,\text{Proposed}}$ | 1.000 | 1.000 | 2,6403.029 |
| $\tau_{b0}$ | 1.000 | 1.001 | 7,358.389 |
| $\tau_{b\text{Proposed}}$ | 1.000 | 1.000 | 9,447.571 |
| $\tau_{c}$ | 1.000 | 1.000 | 1,2668.569 |
| $R_{12}$ | 1.000 | 1.000 | 1,1599.114 |

**Supplementary Table 48: Posterior summaries for Model 3 of firing identification sensitivity in tracked TA MUs across testing visits.**

| | 50% | 95% ETI | | Mode | 95% HDI | |
|---|---|---|---|---|---|---|
| | | 2.5% | 97.5% | | low | high |
| $\alpha_{\mu}$ | 0.230 | 0.146 | 0.337 | 0.233 | 0.143 | 0.332 |
| $\alpha_{\phi}$ | 11.141 | 8.424 | 14.418 | 11.043 | 8.278 | 14.257 |
| $\alpha_{\zeta}$ | 0.013 | 0.003 | 0.043 | 0.009 | 0.001 | 0.037 |
| $\alpha_{\kappa}$ | 0.253 | 0.012 | 0.907 | 0.055 | 0.001 | 0.852 |
| $\beta_{\mu,\text{Proposed}}$ | 35.258 | 20.413 | 60.053 | 33.004 | 18.812 | 56.833 |
| $\beta_{\phi,\text{Proposed}}$ | 0.922 | 0.613 | 1.367 | 0.911 | 0.580 | 1.320 |
| $\beta_{\zeta,\text{Proposed}}$ | 8.261 | 2.132 | 48.618 | 4.999 | 0.848 | 35.860 |
| $\beta_{\kappa,\text{Proposed}}$ | 47.391 | 0.763 | 4,531.740 | -56.369 | 0.015 | 2,054.125 |
| $\tau_{b0}$ | 0.837 | 0.578 | 1.298 | 0.793 | 0.553 | 1.246 |
| $\tau_{b\text{Proposed}}$ | 0.750 | 0.460 | 1.213 | 0.698 | 0.421 | 1.154 |
| $\tau_{c}$ | 0.057 | 0.002 | 0.201 | 0.015 | 0.000 | 0.173 |
| $R_{12}$ | -0.918 | -0.992 | -0.566 | -0.967 | -1.000 | -0.655 |

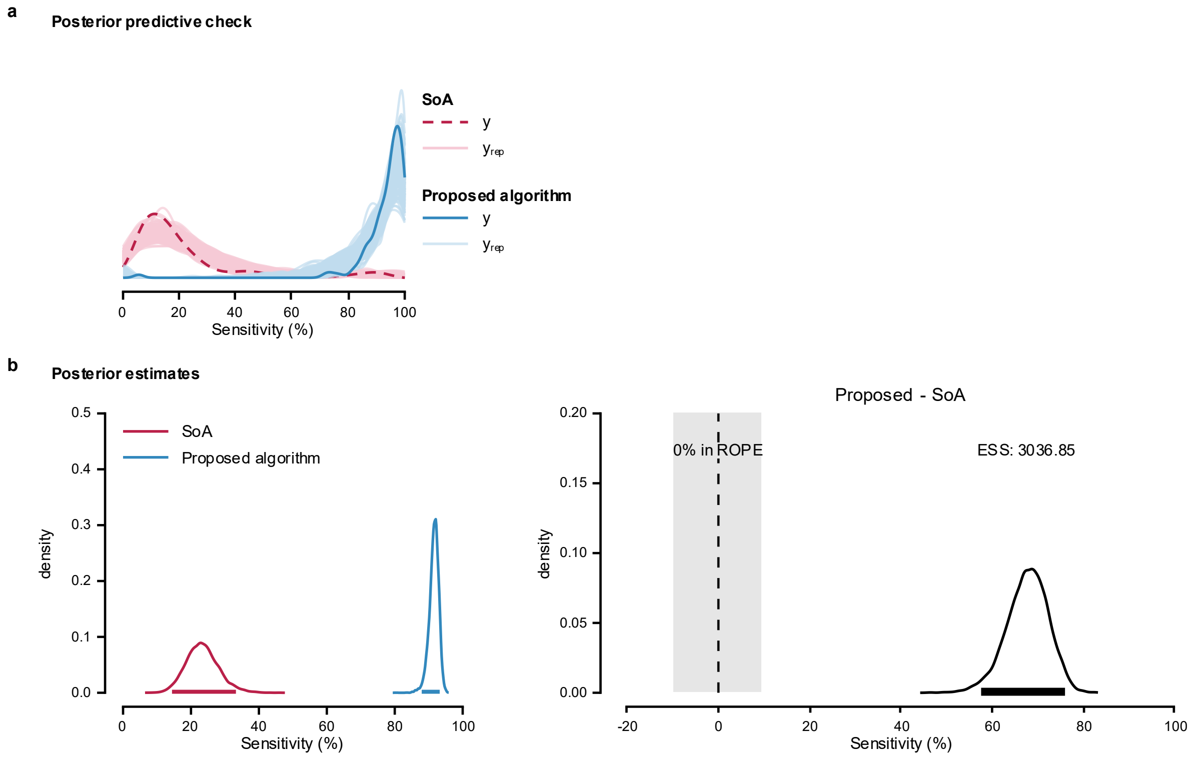


**Supplementary Fig. 34: Posterior predictive check and posterior estimates for Model 3 of firing identification sensitivity in tracked TA MUs across testing visits.**

**a**, Posterior predictive check of the model; **b**, Posterior distributions and the difference in means from Bayesian Model 3, used to compare the performance of the proposed algorithm and SoA method.

### MODEL COMPARISON

Model 2 showed the greatest expected predictive accuracy. ELPD diff relative to Model 2 were −4.0 (SE = 3.1) for Model 3 and −16.0 (SE = 5.3) for Model 1, suggesting similar predictive performance for Models 2 and 3 and reduced performance for Model 1.

**Supplementary Note 14. Comparison of firing identification precision in tracked TA MUs across testing visits.**

Model implementation, likelihood function, and priors are detailed in Supplementary Note 5. The full dataset contains firing identification precision values for 119 tracked MUs in TA muscle (hereafter referred to as 'precision'). The MUs firings were obtained by applying the MU filters calculated using the SoA algorithm[5] and the proposed algorithm to the HDsEMG signals obtained in the second testing visit.  Distribution of the data is shown in Supplementary Fig. 35.

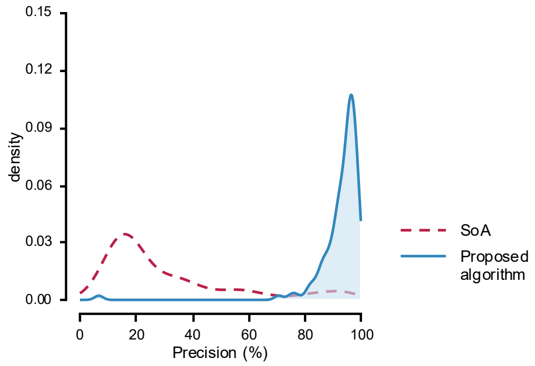


**Supplementary Fig. 35: Distribution of firing identification precision in tracked TA MUs across testing visits.**

Precision values were calculated from firing identification obtained by applying MU filters calculated using SoA (dashed line) and the proposed algorithms (solid line) to the HDsEMG signals generated with synthetic grid displacement.

In all models, ZOIB parameters were back transformed into their interpretable component scales: *mu*, *zoi*, and *coi* to probabilities, and *phi* to the positive precision scale. Group-level SDs are reported on their native model/link scale.

Model 1

The maximum PSRF across all parameters was 1.003, indicating good convergence, while the minimum ESS was 4,540.438, indicating stable estimates of the bounds of the credible intervals (Supplementary Table 49). Posterior summaries are reported as medians with 95% ETIs and 95% HDIs (Supplementary Table 50).

Posterior predictive checks were performed by overlaying model-generated predictions on the observed data (Supplementary Fig. 36a). The posterior distributions (Supplementary Fig. 36b) indicated that the proposed algorithm achieved a higher precision (91.2%, [87.1-94.2%]) compared to the SoA algorithm (30.3%, [22.0-39.3%]). The posterior distribution of the difference excluded zero, and less than 2.5% of its distribution fell within ROPE.

**Supplementary Table 49: Posterior MCMC diagnostics for Model 1 of firing identification precision in tracked TA MUs across testing visits.**

| | **PSRF Pt** | **PSRF UpCI** | **ESS** |
|---|---|---|---|
| $\alpha_{\mu}$ | 1.001 | 1.002 | 4,540.438 |
| $\alpha_{\phi}$ | 1.000 | 1.001 | 1,5449.885 |
| $\alpha_{\zeta}$ | 1.000 | 1.001 | 1,5118.605 |
| $\alpha_{\kappa}$ | 1.000 | 1.001 | 1,6093.714 |
| $\beta_{\mu,\text{Proposed}}$ | 1.000 | 1.001 | 1,5564.318 |
| $\beta_{\phi,\text{Proposed}}$ | 1.000 | 1.000 | 1,8582.117 |
| $\beta_{\zeta,\text{Proposed}}$ | 1.000 | 1.001 | 1,8112.688 |
| $\beta_{\kappa,\text{Proposed}}$ | 1.000 | 1.001 | 1,8588.570 |
| $\tau_{b}$ | 1.001 | 1.003 | 6,051.129 |

**Supplementary Table 50: Posterior summaries for Model 1 of firing identification precision in tracked TA MUs across testing visits.**

| | 50% | 95% ETI | | Mode | 95% HDI | |
|---|---|---|---|---|---|---|
| | | 2.5% | 97.5% | | low | high |
| $\alpha_{\mu}$ | 0.300 | 0.218 | 0.389 | 0.297 | 0.218 | 0.389 |
| $\alpha_{\phi}$ | 6.870 | 5.046 | 9.047 | 6.860 | 4.960 | 8.930 |
| $\alpha_{\zeta}$ | 0.015 | 0.003 | 0.045 | 0.010 | 0.001 | 0.039 |
| $\alpha_{\kappa}$ | 0.352 | 0.017 | 0.936 | 0.067 | 0.004 | 0.901 |
| $\beta_{\mu,\text{Proposed}}$ | 26.418 | 20.655 | 33.566 | 26.016 | 20.231 | 33.048 |
| $\beta_{\phi,\text{Proposed}}$ | 1.626 | 1.086 | 2.452 | 1.565 | 1.008 | 2.350 |
| $\beta_{\zeta,\text{Proposed}}$ | 2.076 | 0.385 | 13.144 | 1.292 | 0.086 | 9.588 |
| $\beta_{\kappa,\text{Proposed}}$ | 8.321 | 0.086 | 1,250.553 | -11.953 | 0.001 | 553.799 |
| $\tau_b$ | 0.635 | 0.387 | 1.073 | 0.587 | 0.344 | 1.002 |

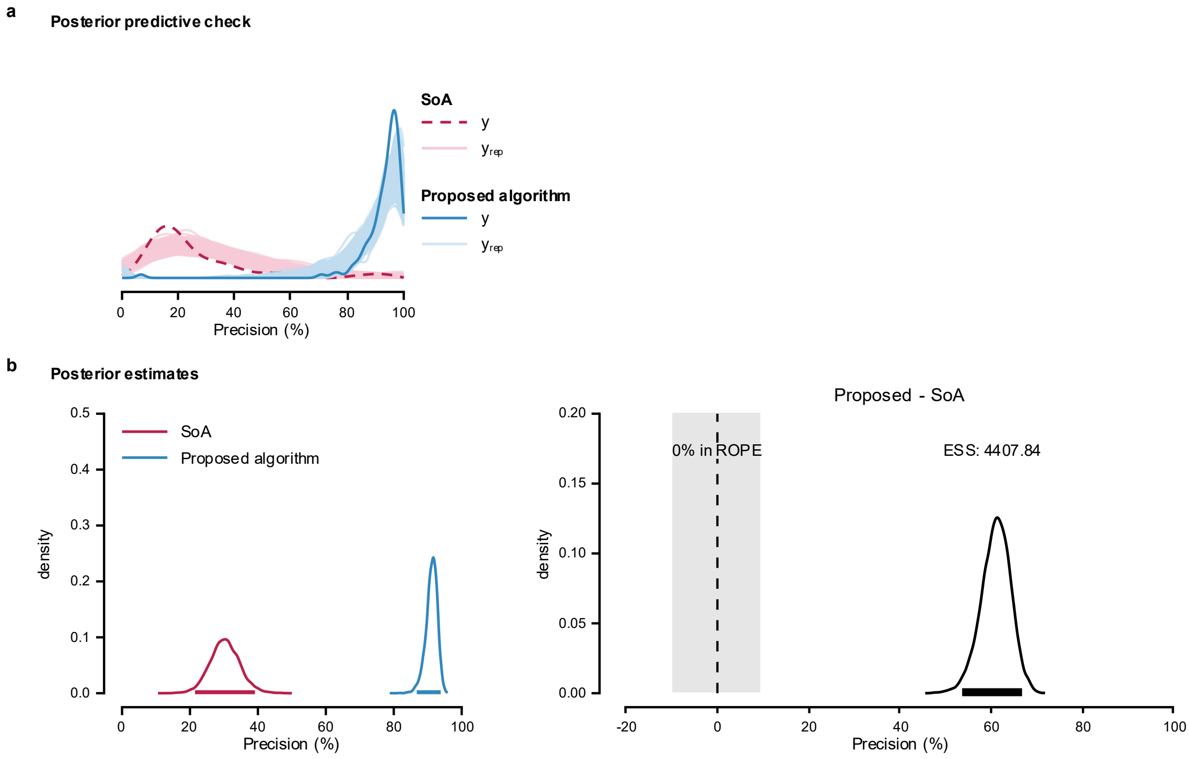


**Supplementary Fig. 36: Posterior predictive check and posterior estimates for Model 1 of firing identification precision in tracked TA MUs across testing visits.**
**a**, Posterior predictive check of the model; **b**, Posterior distributions and the difference between means estimated using Bayesian Model 1 to compare the performance of the proposed algorithm and SoA method.

<u>Model 2</u>

The maximum PSRF across all parameters was 1.002, indicating good convergence, while the minimum ESS was 4,533.320, indicating stable estimates of the bounds of the credible intervals (Supplementary Table 51). Posterior summaries are reported in Supplementary Table 52 as medians with 95% ETIs and 95% HDIs. Group-level correlations are reported on the correlation scale.

Posterior predictive checks were performed by overlaying model-generated predictions on the observed data (Supplementary Fig. 37a). The posterior distributions (Supplementary Fig.

37b) indicated that the proposed algorithm achieved a higher precision (90.8%, [87.7-93.1%]) compared to the SoA algorithm (31.1%, [21.6-41.5%]). The posterior distribution of the difference excluded zero, and less than 2.5% of its distribution fell within ROPE.

**Supplementary Table 51: Posterior MCMC diagnostics for Model 2 of firing identification precision in tracked TA MUs across testing visits.**

| | PSRF Pt | PSRF UpCI | ESS |
|---|---|---|---|
| $\alpha_{\mu}$ | 1.001 | 1.002 | 4,533.320 |
| $\alpha_{\phi}$ | 1.000 | 1.000 | 2,0022.001 |
| $\alpha_{\zeta}$ | 1.000 | 1.001 | 1,6363.712 |
| $\alpha_{\kappa}$ | 1.000 | 1.001 | 1,9524.266 |
| $\beta_{\mu,\text{Proposed}}$ | 1.001 | 1.002 | 5,984.829 |
| $\beta_{\phi,\text{Proposed}}$ | 1.000 | 1.000 | 2,0163.696 |
| $\beta_{\zeta,\text{Proposed}}$ | 1.000 | 1.001 | 1,7875.132 |
| $\beta_{\kappa,\text{Proposed}}$ | 1.000 | 1.001 | 2,1373.496 |
| $\tau_{b0}$ | 1.000 | 1.001 | 6,405.917 |
| $\tau_{b\text{Proposed}}$ | 1.000 | 1.001 | 8,052.448 |
| $R_{12}$ | 1.001 | 1.001 | 1,0623.779 |

**Supplementary Table 52: Posterior summaries for Model 2 of firing identification precision in tracked TA MUs across testing visits.**

| | 50% | 95% ETI | | Mode | 95% HDI | |
|---|---|---|---|---|---|---|
| | | 2.5% | 97.5% | | low | high |
| $\alpha_{\mu}$ | 0.307 | 0.214 | 0.413 | 0.306 | 0.211 | 0.410 |
| $\alpha_{\phi}$ | 8.126 | 6.185 | 10.432 | 7.985 | 6.087 | 10.318 |
| $\alpha_{\zeta}$ | 0.015 | 0.003 | 0.044 | 0.010 | 0.001 | 0.038 |
| $\alpha_{\kappa}$ | 0.348 | 0.018 | 0.940 | 0.072 | 0.003 | 0.906 |
| $\beta_{\mu,\text{Proposed}}$ | 24.027 | 14.850 | 38.650 | 22.374 | 14.127 | 37.301 |
| $\beta_{\phi,\text{Proposed}}$ | 1.430 | 0.980 | 2.111 | 1.393 | 0.919 | 2.023 |
| $\beta_{\zeta,\text{Proposed}}$ | 2.069 | 0.402 | 13.479 | 1.132 | 0.107 | 9.879 |
| $\beta_{\kappa,\text{Proposed}}$ | 8.276 | 0.084 | 1,187.696 | -11.894 | 0.000 | 542.914 |
| $\tau_{b0}$ | 0.744 | 0.502 | 1.177 | 0.695 | 0.463 | 1.102 |
| $\tau_{b\text{Proposed}}$ | 0.658 | 0.387 | 1.079 | 0.615 | 0.348 | 1.020 |
| $R_{12}$ | -0.914 | -0.991 | -0.563 | -0.965 | -0.999 | -0.644 |

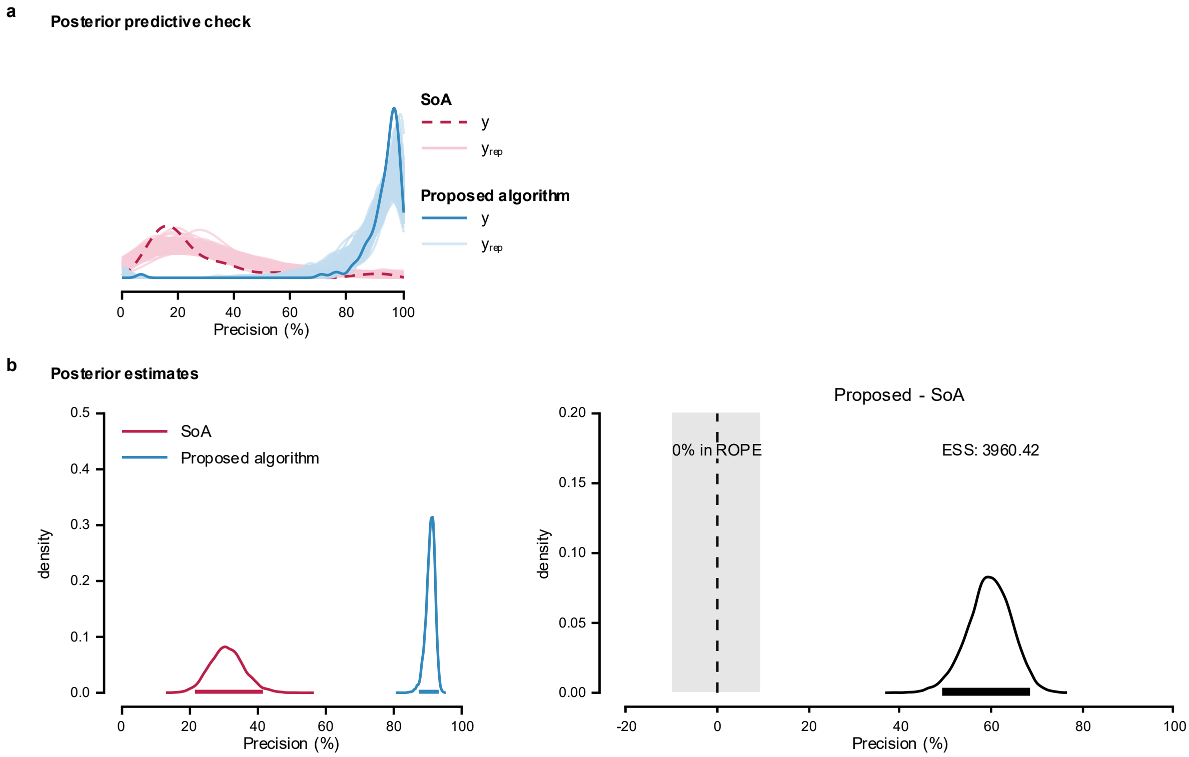


**Supplementary Fig. 37: Posterior predictive check and posterior estimates for Model 2 of firing identification precision in tracked TA MUs across testing visits.**

**a**, Posterior predictive check of the model; **b**, Posterior distributions and the difference between means estimated using Bayesian Model 2 to compare the performance of the proposed algorithm and SoA method.

Model 3

The maximum PSRF across all parameters was 1.006, indicating good convergence, while the minimum ESS was 3,747.211, indicating stable estimates of the bounds of the credible intervals (Supplementary Table 53). Posterior summaries are reported in Supplementary Table 54 as medians with 95% ETIs and 95% HDIs. Group-level correlations are reported on the correlation scale.

Posterior predictive checks were performed by overlaying model-generated predictions on the observed data (Supplementary Fig. 38a). The posterior distributions (Supplementary Fig.

38b) indicated that the proposed algorithm achieved a higher precision (90.8%, [87.6-93.2%]) compared to the SoA algorithm (30.8%, [21.2-41.2%]). The posterior distribution of the difference excluded zero, and less than 2.5% of its distribution fell within ROPE.

**Supplementary Table 53: Posterior MCMC diagnostics for Model 3 of firing identification precision in tracked TA MUs across testing visits.**

| | PSRF Pt | PSRF UpCI | ESS |
|---|---|---|---|
| $\alpha_{\mu}$ | 1.002 | 1.006 | 3,747.211 |
| $\alpha_{\phi}$ | 1.000 | 1.000 | 2,4548.305 |
| $\alpha_{\zeta}$ | 1.000 | 1.000 | 1,9715.834 |
| $\alpha_{\kappa}$ | 1.000 | 1.000 | 2,3351.139 |
| $\beta_{\mu,\text{Proposed}}$ | 1.001 | 1.004 | 4,987.113 |
| $\beta_{\phi,\text{Proposed}}$ | 1.000 | 1.000 | 2,8334.340 |
| $\beta_{\zeta,\text{Proposed}}$ | 1.000 | 1.000 | 2,4179.562 |
| $\beta_{\kappa,\text{Proposed}}$ | 1.000 | 1.000 | 2,8037.912 |
| $\tau_{b0}$ | 1.000 | 1.001 | 6,566.549 |
| $\tau_{b\text{Proposed}}$ | 1.000 | 1.001 | 8,418.065 |
| $\tau_{c}$ | 1.000 | 1.001 | 1,0474.429 |
| $R_{12}$ | 1.001 | 1.001 | 1,2333.569 |

**Supplementary Table 54: Posterior summaries for Model 3 of firing identification precision in tracked TA MUs across synthetic grid displacement.**

| | 50% | 95% ETI | | Mode | 95% HDI | |
|---|---|---|---|---|---|---|
| | | 2.5% | 97.5% | | low | high |
| $\alpha_{\mu}$ | 0.305 | 0.212 | 0.411 | 0.305 | 0.209 | 0.407 |
| $\alpha_{\phi}$ | 8.202 | 6.210 | 10.639 | 7.988 | 6.118 | 10.519 |
| $\alpha_{\zeta}$ | 0.015 | 0.003 | 0.043 | 0.010 | 0.001 | 0.038 |
| $\alpha_{\kappa}$ | 0.354 | 0.017 | 0.943 | 0.074 | 0.004 | 0.910 |
| $\beta_{\mu,\text{Proposed}}$ | 24.273 | 14.967 | 38.998 | 22.776 | 14.044 | 37.212 |
| $\beta_{\phi,\text{Proposed}}$ | 1.418 | 0.965 | 2.085 | 1.360 | 0.929 | 2.026 |
| $\beta_{\zeta,\text{Proposed}}$ | 2.058 | 0.392 | 12.788 | 1.450 | 0.108 | 9.303 |
| $\beta_{\kappa,\text{Proposed}}$ | 7.937 | 0.077 | 1,207.445 | -11.383 | 0.001 | 531.034 |
| $\tau_{b0}$ | 0.745 | 0.506 | 1.169 | 0.716 | 0.478 | 1.119 |
| $\tau_{b\text{Proposed}}$ | 0.658 | 0.389 | 1.091 | 0.620 | 0.357 | 1.041 |
| $\tau_{c}$ | 0.061 | 0.003 | 0.212 | 0.018 | 0.000 | 0.184 |
| $R_{12}$ | -0.915 | -0.992 | -0.550 | -0.967 | -0.999 | -0.638 |

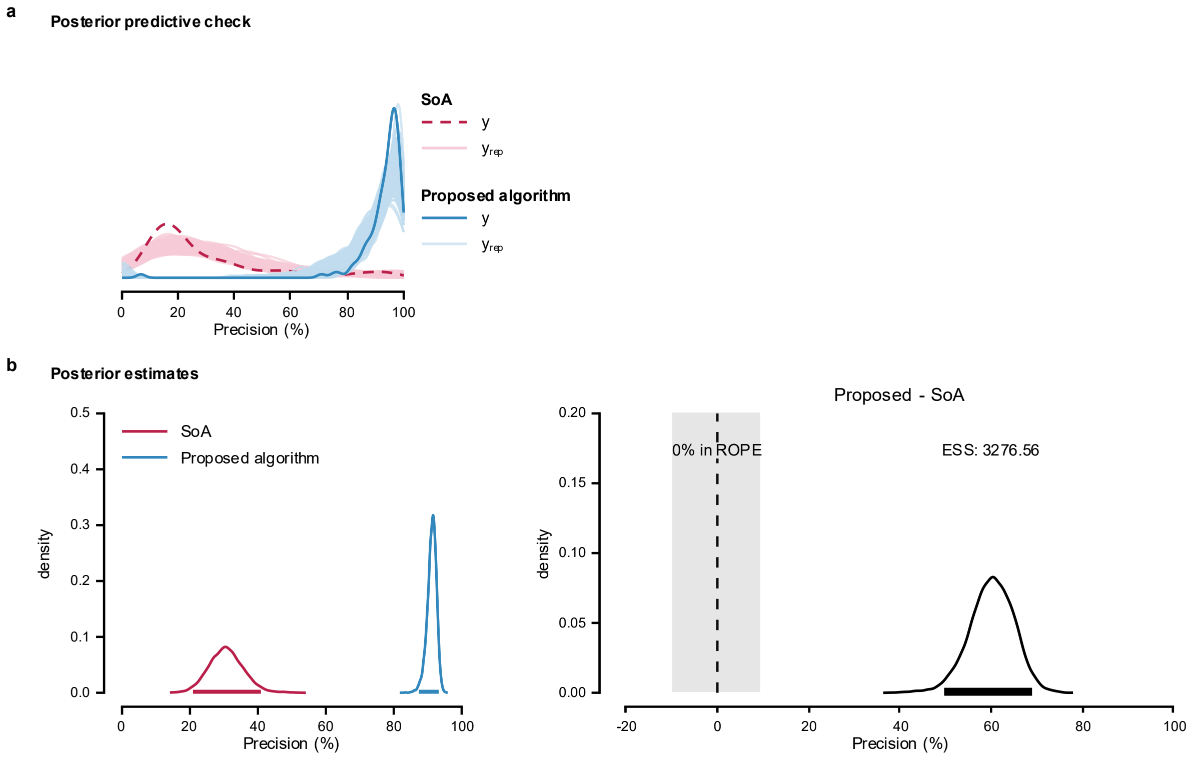


**Supplementary Fig. 38: Posterior predictive check and posterior estimates for Model 3 of firing identification precision in tracked TA MUs across testing visits.**

**a**, Posterior predictive check of the model; **b**, Posterior distributions and the difference between means estimated using Bayesian Model 1 to compare the performance of the proposed algorithm and SoA method.

### MODEL COMPARISON

Model 2 showed the greatest expected predictive accuracy. ELPD diff relative to Model 2 were −1.3 (SE = 0.7) for Model 3 and −14.8 (SE = 5.2) for Model 1, suggesting similar predictive performance for Models 2 and 3 and reduced performance for Model 1.

**Supplementary Note 15. Comparison of firing identification PNR for tracked MG MUs across testing visits.**

Model implementation, likelihood function, and priors are detailed in Supplementary Note 5. The full dataset contains firing identification PNR values for 113 tracked MUs in MG muscle (Supplementary Fig. 39). The MUs firings were obtained by applying the MU filters calculated using the SoA algorithm[5] and the proposed algorithm to the HDsEMG signals obtained in the second testing visit.

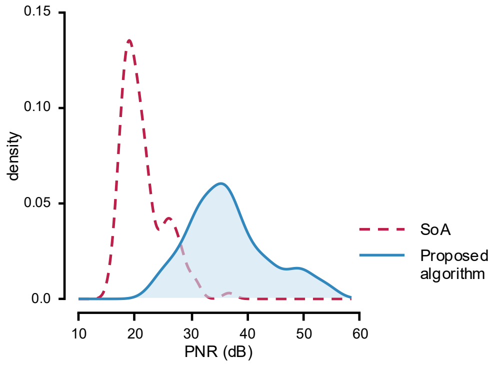


**Supplementary Fig. 39: Distribution of firing identification PNR in tracked MG MUs across testing visits.**

PNR values were calculated from firing identification obtained by applying MU filters calculated using SoA (dashed line) and the proposed algorithms (solid line) to the HDsEMG signals generated with synthetic grid displacement.

In all models, the lognormal mean and sigma effects were exponentiated before the posterior summaries were calculated. Group-level SDs are reported on their native model or link scales.

Model 1

The maximum PSRF across all parameters was 1.001, indicating good convergence, while the minimum ESS was 5,873.055, indicating stable estimates of the bounds of the credible intervals (Supplementary Table 55). Supplementary Table 56 provides median of the estimate value in the original scale of PNR, the limits of the 95% ETI, and the limits of the 95% HDI.

**Supplementary Table 55: Posterior MCMC diagnostics for Model 1 of firing identification PNR in tracked MG MUs across testing visits.**

| | PSRF Pt | PSRF UpCI | ESS |
|---|---|---|---|
| $\alpha_{\mu}$ | 1.000 | 1.000 | 6,332.681 |
| $\alpha_{\sigma}$ | 1.000 | 1.000 | 1,6503.502 |
| $\beta_{\mu,\text{Proposed}}$ | 1.000 | 1.000 | 2,4740.583 |
| $\beta_{\sigma,\text{Proposed}}$ | 1.000 | 1.000 | 1,6925.304 |
| $\tau_{\mu}$ | 1.001 | 1.001 | 5,873.055 |

**Supplementary Table 56: Posterior summaries for Model 1 of firing identification PNR in tracked MG MUs across testing visits.**

| | 50% | 95% ETI | | Mode | 95% HDI | |
|---|---|---|---|---|---|---|
| | | 2.5% | 97.5% | | low | high |
| $\alpha_{\mu}$ | 21.612 | 20.207 | 23.165 | 21.619 | 20.200 | 23.155 |
| $\alpha_{\sigma}$ | 1.701 | 1.629 | 1.779 | 1.699 | 1.626 | 1.775 |
| $\beta_{\mu,\text{Proposed}}$ | 0.155 | 0.134 | 0.181 | 0.154 | 0.133 | 0.180 |
| $\beta_{\sigma,\text{Proposed}}$ | 1.173 | 0.930 | 1.470 | 1.164 | 0.919 | 1.455 |
| $\tau_{\mu}$ | 0.086 | 0.047 | 0.167 | 0.077 | 0.043 | 0.156 |

Posterior predictive checks were performed by overlaying model-generated predictions on the observed data (Supplementary Fig. 40a). The posterior distributions (Supplementary Fig. 40b) indicated that the proposed algorithm achieved a higher PNR (37.4 dB, [34.9-40.1 dB]) compared to the SoA algorithm (21.9 dB, [20.4-23.4 dB]). The posterior distribution of the difference excluded zero, and less than 2.5% of its distribution fell within ROPE.

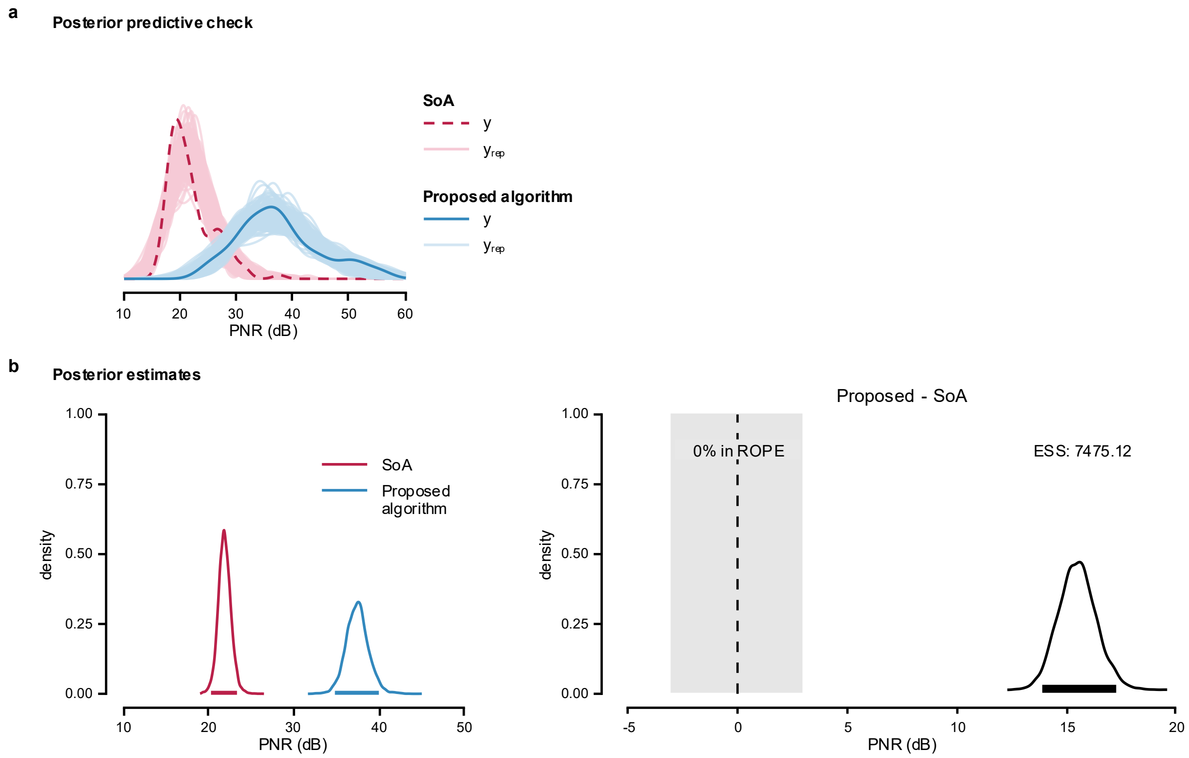


**Supplementary Fig. 40: Posterior predictive check and posterior estimates for Model 1 of firing identification PNR in tracked MG MUs across testing visits.**

**a**, Posterior predictive check of the model; **b**, Posterior distributions and the difference between means estimated using Bayesian Model 1 to compare the performance of the proposed algorithm and SoA method.

Model 2

The maximum PSRF across all parameters was 1.002, indicating good convergence, while the minimum ESS was 5,659.873, indicating stable estimates of the bounds of the credible intervals (Supplementary Table 57). Posterior parameter summaries are given as medians with 95% ETIs and 95% HDIs (Supplementary Table 58). Correlations were summarized on their native scales.

Posterior predictive checks were performed by overlaying model-generated predictions on the observed data (Supplementary Fig. 41a). The posterior distributions (Supplementary Fig. 41b) indicated that the proposed algorithm achieved a higher PNR (35.3 dB, [30.6-39.6 dB]) than the SoA algorithm (22.6 dB, [20.7-25.0 dB]). The posterior distribution of the difference excluded zero, and less than 2.5% of its distribution fell within ROPE.

Model 3

The maximum PSRF across all parameters was 1.004, indicating good convergence, while the minimum ESS was 2,644.540, indicating stable estimates of the bounds of the credible intervals (Supplementary Table 59). Posterior parameter summaries are given as medians with 95% ETIs and 95% HDIs (Supplementary Table 60). Correlations were summarized on their native scales.

Posterior predictive checks were performed by overlaying model-generated predictions on the observed data (Supplementary Fig. 42a). The posterior distributions (Supplementary Fig. 42b) indicated that the proposed algorithm achieved a higher PNR (35.3 dB, [30.4-39.6 dB]) than the SoA algorithm (22.6 dB, [20.7-24.9 dB]). The posterior distribution of the difference excluded zero, and less than 2.5% of its distribution fell within ROPE.

**Supplementary Table 57: Posterior MCMC diagnostics for Model 2 of firing identification PNR in tracked MG MUs across testing visits.**

| | PSRF Pt | PSRF UpCI | ESS |
|---|---|---|---|
| $\alpha_{\mu}$ | 1.001 | 1.002 | 5,805.282 |
| $\alpha_{\sigma}$ | 1.000 | 1.000 | 1,9194.010 |
| $\beta_{\mu,\text{Proposed}}$ | 1.001 | 1.002 | 5,865.638 |
| $\beta_{\sigma,\text{Proposed}}$ | 1.000 | 1.000 | 2,1977.249 |
| $\tau_{b0}$ | 1.000 | 1.000 | 5,828.215 |
| $\tau_{b\text{Proposed}}$ | 1.001 | 1.002 | 5,659.873 |
| $R_{12}$ | 1.000 | 1.000 | 7,316.349 |

**Supplementary Table 58: Posterior summaries for Model 2 of firing identification PNR in tracked MG MUs across testing visits.**

| | 50% | 95% ETI | | Mode | 95% HDI | |
|---|---|---|---|---|---|---|
| | | 2.5% | 97.5% | | low | high |
| $\alpha_{\mu}$ | 22.302 | 20.478 | 24.764 | 22.262 | 20.339 | 24.574 |
| $\alpha_{\sigma}$ | 0.134 | 0.118 | 0.154 | 0.132 | 0.117 | 0.153 |
| $\beta_{\mu,\text{Proposed}}$ | 1.568 | 1.318 | 1.780 | 1.573 | 1.332 | 1.791 |
| $\beta_{\sigma,\text{Proposed}}$ | 1.130 | 0.943 | 1.356 | 1.127 | 0.932 | 1.340 |
| $\tau_{b0}$ | 0.127 | 0.077 | 0.232 | 0.118 | 0.068 | 0.213 |
| $\tau_{b\text{Proposed}}$ | 0.206 | 0.127 | 0.378 | 0.182 | 0.116 | 0.347 |
| $R_{12}$ | -0.558 | -0.873 | 0.071 | -0.653 | -0.918 | -0.030 |

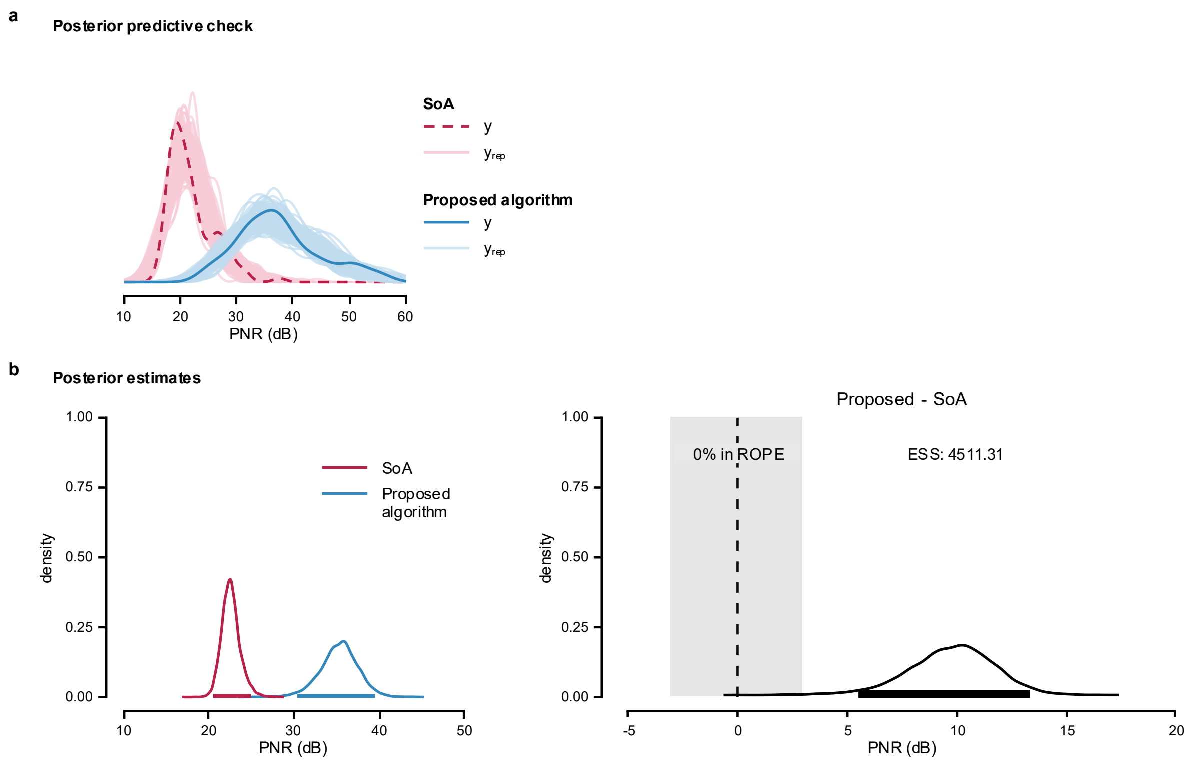


**Supplementary Fig. 41: Posterior predictive check and posterior estimates for Model 2 of firing identification PNR in tracked MG muscle MUs across testing visits.**

**a**, Posterior predictive check of the model; **b**, Posterior distributions and the difference in means from Bayesian Model 2, used to compare the performance of the proposed algorithm and SoA method.

**Supplementary Table 59: Posterior MCMC diagnostics for Model 3 of firing identification PNR in tracked MG MUs across testing visits.**

| | PSRF Pt | PSRF UpCI | ESS |
|---|---|---|---|
| $\alpha_{\mu}$ | 1.000 | 1.001 | 4,358.505 |
| $\alpha_{\sigma}$ | 1.000 | 1.001 | 6,124.032 |
| $\beta_{\mu,\text{Proposed}}$ | 1.000 | 1.001 | 4,564.199 |
| $\beta_{\sigma,\text{Proposed}}$ | 1.000 | 1.000 | 1,8511.083 |
| $\tau_{b0}$ | 1.001 | 1.003 | 6,140.874 |
| $\tau_{b\text{Proposed}}$ | 1.000 | 1.000 | 4,941.607 |
| $\tau_{c}$ | 1.001 | 1.004 | 2,644.540 |
| $R_{12}$ | 1.000 | 1.001 | 5,771.900 |

**Supplementary Table 60: Posterior summaries for Model 3 of firing identification PNR in tracked MG MUs across testing visits.**

| | 50% | 95% ETI | | Mode | 95% HDI | |
|---|---|---|---|---|---|---|
| | | 2.5% | 97.5% | | low | high |
| $\alpha_{\mu}$ | 22.312 | 20.480 | 24.718 | 22.169 | 20.257 | 24.455 |
| $\alpha_{\sigma}$ | 0.127 | 0.106 | 0.148 | 0.128 | 0.106 | 0.148 |
| $\beta_{\mu,\text{Proposed}}$ | 1.567 | 1.319 | 1.775 | 1.587 | 1.323 | 1.778 |
| $\beta_{\sigma,\text{Proposed}}$ | 1.142 | 0.936 | 1.403 | 1.110 | 0.926 | 1.390 |
| $\tau_{b0}$ | 0.127 | 0.076 | 0.226 | 0.114 | 0.069 | 0.211 |
| $\tau_{b\text{Proposed}}$ | 0.208 | 0.129 | 0.377 | 0.186 | 0.114 | 0.344 |
| $\tau_{c}$ | 0.043 | 0.003 | 0.078 | 0.050 | 0.000 | 0.073 |
| $R_{12}$ | -0.562 | -0.875 | 0.053 | -0.662 | -0.910 | -0.034 |

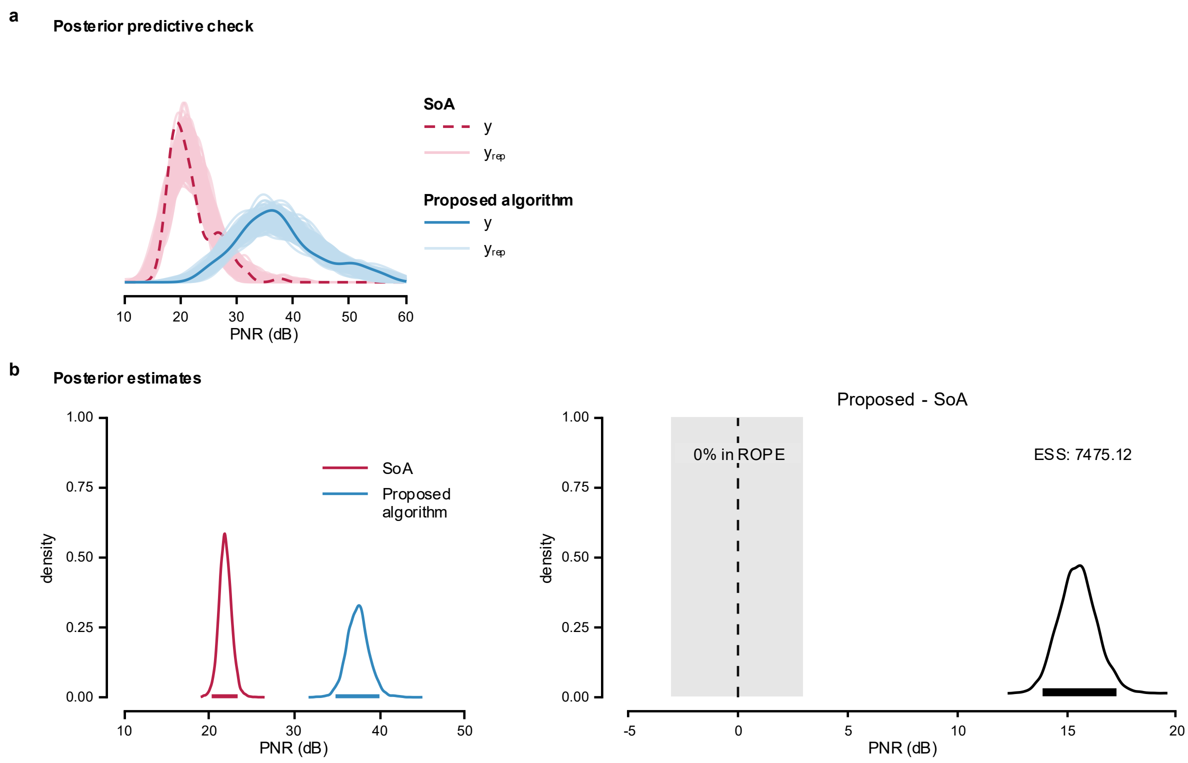


**Supplementary Fig. 42: Posterior predictive check and posterior estimates for Model 3 of firing identification PNR in tracked MG MUs across testing visits.**

**a**, Posterior predictive check of the model; **b**, Posterior distributions and the difference in means from Bayesian Model 3, used to compare the performance of the proposed algorithm and SoA method.

MODEL COMPARISON

Model 2 showed the greatest expected predictive accuracy. ELPD diff relative to Model 2 were −0.01 (SE = 1.7) for Model 3 and −34.0 (SE = 7.6) for Model 1, suggesting similar predictive performance for Models 2 and 3 and reduced performance for Model 1.

**Supplementary Note 16. Comparison of firing identification sensitivity in tracked MG MUs across testing visits.**

Model implementation, likelihood function, and priors are detailed in Supplementary Note 5. The full dataset contains firing identification sensitivity values for 113 tracked MUs (hereafter referred to as 'sensitivity') in MG muscle. The MUs firings were obtained by applying the MU filters calculated using the SoA algorithm[5] and the proposed algorithm to the HDsEMG signals obtained in the second testing visit. Distribution of the data is shown in Supplementary Fig. 43.

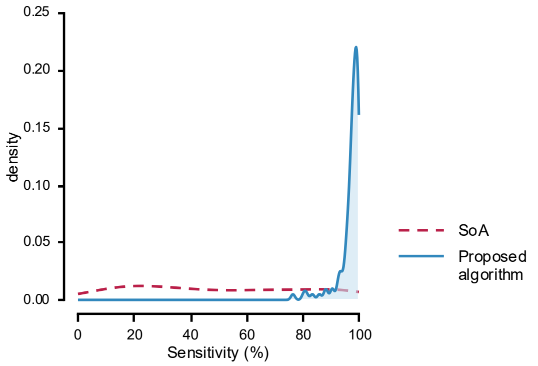


**Supplementary Fig. 43: Distribution of firing identification sensitivity in tracked MG MUs across testing visits.**
Sensitivity values were calculated from firing identification obtained by applying MU filters calculated using SoA (dashed line) and the proposed algorithms (solid line) to the HDsEMG signals generated with synthetic grid displacement.

In all models, ZOIB parameters were back transformed into their interpretable component scales: *mu*, *zoi*, and *coi* to probabilities, and *phi* to the positive precision scale. Group-level SDs are reported on their native model/link scale.

Model 1

The maximum PSRF across all parameters was 1.004, indicating good convergence, while the minimum ESS was 4,839.106, indicating stable estimates of the bounds of the credible intervals (Supplementary Table 61). Posterior summaries are reported in Supplementary Table 62 as medians with 95% ETIs and 95% HDIs. Group-level correlations are reported on the correlation scale.

Posterior predictive checks were performed by overlaying model-generated predictions on the observed data (Supplementary Fig. 44a). The posterior distributions (Supplementary Fig. 44a) indicated that the proposed algorithm achieved a higher sensitivity (96.1%, [93.8-97.5 %]) than the SoA algorithm (51.4%, [43.3-59.0%]). The posterior distribution of the difference excluded zero, and less than 2.5% of its distribution fell within ROPE.

**Supplementary Table 61: Posterior MCMC diagnostics for Model 1 of firing identification sensitivity in tracked MG MUs across testing visits.**

| | PSRF Pt | PSRF UpCI | ESS |
|---|---|---|---|
| $\alpha_\mu$ | 1.000 | 1.000 | 7,797.901 |
| $\alpha_\phi$ | 1.000 | 1.000 | 1,2026.624 |
| $\alpha_\zeta$ | 1.001 | 1.002 | 1,0378.099 |
| $\alpha_\kappa$ | 1.000 | 1.000 | 1,3885.065 |
| $\beta_{\mu,\text{Proposed}}$ | 1.000 | 1.000 | 1,6491.057 |
| $\beta_{\phi,\text{Proposed}}$ | 1.000 | 1.000 | 1,3513.627 |
| $\beta_{\zeta,\text{Proposed}}$ | 1.001 | 1.002 | 1,1425.987 |
| $\beta_{\kappa,\text{Proposed}}$ | 1.000 | 1.000 | 1,5895.304 |
| $\tau_b$ | 1.001 | 1.004 | 4,839.106 |

**Supplementary Table 62: Posterior summaries for Model 1 of firing identification sensitivity in tracked MG MUs across testing visits.**

| | 50% | 95% ETI | | Mode | 95% HDI | |
|---|---|---|---|---|---|---|
| | | 2.5% | 97.5% | | low | high |
| $\alpha_{\mu}$ | 0.518 | 0.435 | 0.594 | 0.520 | 0.435 | 0.594 |
| $\alpha_{\phi}$ | 2.257 | 1.725 | 2.935 | 2.236 | 1.711 | 2.909 |
| $\alpha_{\zeta}$ | 0.014 | 0.002 | 0.043 | 0.009 | 0.001 | 0.037 |
| $\alpha_{\kappa}$ | 0.233 | 0.011 | 0.898 | 0.055 | 0.001 | 0.841 |
| $\beta_{\mu,\text{Proposed}}$ | 26.266 | 19.897 | 34.704 | 25.728 | 19.554 | 34.233 |
| $\beta_{\phi,\text{Proposed}}$ | 13.146 | 8.484 | 19.958 | 12.835 | 8.106 | 19.283 |
| $\beta_{\zeta,\text{Proposed}}$ | 12.022 | 3.101 | 72.367 | 6.865 | 1.274 | 50.916 |
| $\beta_{\kappa,\text{Proposed}}$ | 67.964 | 1.125 | 5,593.709 | -75.612 | 0.015 | 2,599.469 |
| $\tau_{b}$ | 0.335 | 0.070 | 0.753 | 0.302 | 0.009 | 0.665 |

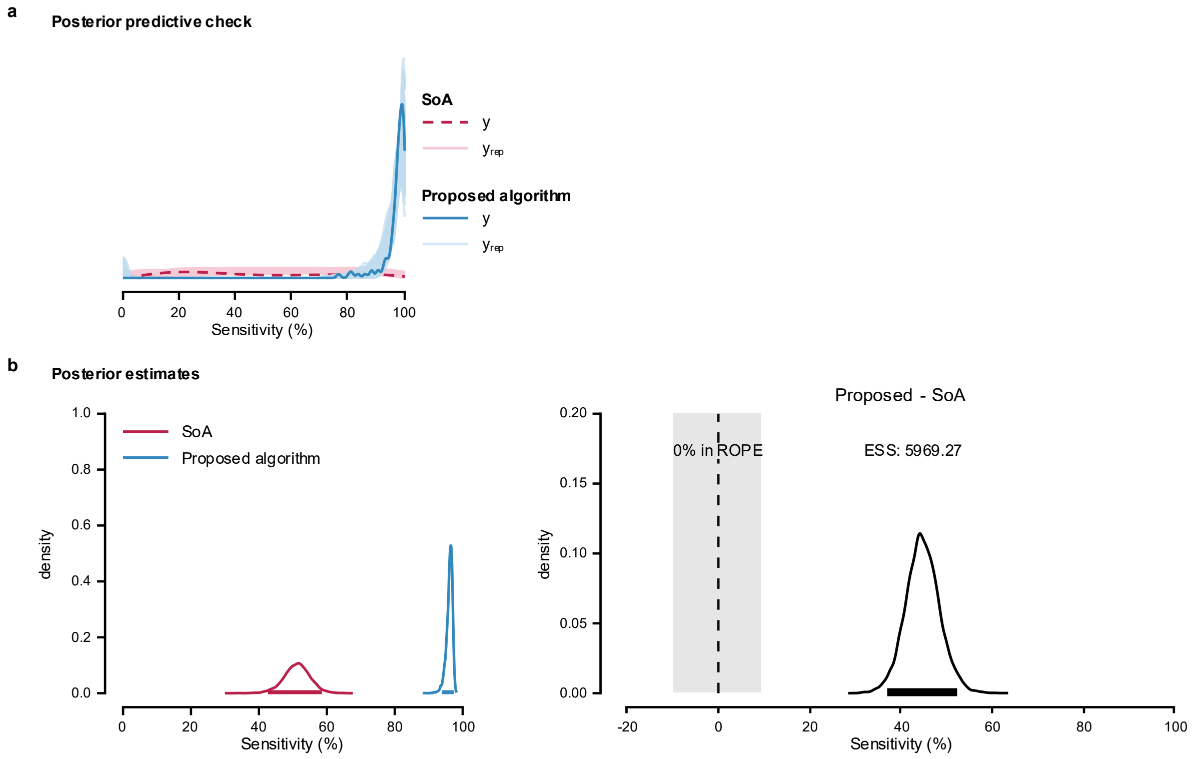


**Supplementary Fig. 44: Posterior predictive check and posterior estimates for Model 1 of firing identification sensitivity in tracked MG MUs across testing visits.**

**a**, Posterior predictive check of the model; **b**, Posterior distributions and the difference between means estimated using Bayesian Model 1 to compare the performance of the proposed algorithm and SoA method.

<u>Model 2</u>

The maximum PSRF across all parameters was 1.003, indicating good convergence, while the minimum ESS was 6,395.834, indicating stable estimates of the bounds of the credible intervals (Supplementary Table 63). Posterior summaries are reported in Supplementary Table 64 as medians with 95% ETIs and 95% HDIs. Group-level correlations are reported on the correlation scale.

Posterior predictive checks were performed by overlaying model-generated predictions on the observed data (Supplementary Fig. 45a). The posterior distributions (Supplementary Fig.

45b) indicated that the proposed algorithm achieved a higher sensitivity (96.0%, [93.6-97.3%]) than the SoA algorithm (52.9%, [39.8-65.1%]). The posterior distribution of the difference excluded zero, and less than 2.5% of its distribution fell within ROPE.

**Supplementary Table 63: Posterior MCMC diagnostics for Model 2 of firing identification sensitivity in tracked MG MUs across testing visits.**

| | PSRF Pt | PSRF UpCI | ESS |
|---|---|---|---|
| $\alpha_{\mu}$ | 1.001 | 1.001 | 6,395.834 |
| $\alpha_{\phi}$ | 1.000 | 1.000 | 2,0443.992 |
| $\alpha_{\zeta}$ | 1.000 | 1.000 | 1,4026.245 |
| $\alpha_{\kappa}$ | 1.000 | 1.000 | 1,8505.763 |
| $\beta_{\mu,\text{Proposed}}$ | 1.000 | 1.000 | 7,832.789 |
| $\beta_{\phi,\text{Proposed}}$ | 1.000 | 1.000 | 2,1415.251 |
| $\beta_{\zeta,\text{Proposed}}$ | 1.000 | 1.000 | 1,4427.101 |
| $\beta_{\kappa,\text{Proposed}}$ | 1.000 | 1.001 | 1,9589.266 |
| $\tau_{b0}$ | 1.001 | 1.002 | 6,746.835 |
| $\tau_{b\text{Proposed}}$ | 1.001 | 1.003 | 7,614.119 |
| $R_{12}$ | 1.001 | 1.002 | 9,532.917 |

**Supplementary Table 64: Posterior summaries for Model 2 of firing identification sensitivity in tracked MG MUs across testing visits.**

| | 50% | 95% ETI | | Mode | 95% HDI | |
|---|---|---|---|---|---|---|
| | | 2.5% | 97.5% | | low | high |
| $\alpha_{\mu}$ | 0.535 | 0.402 | 0.659 | 0.543 | 0.404 | 0.661 |
| $\alpha_{\phi}$ | 3.012 | 2.293 | 3.878 | 2.977 | 2.249 | 3.818 |
| $\alpha_{\zeta}$ | 0.014 | 0.003 | 0.044 | 0.009 | 0.001 | 0.037 |
| $\alpha_{\kappa}$ | 0.235 | 0.012 | 0.896 | 0.054 | 0.001 | 0.838 |
| $\beta_{\mu,\text{Proposed}}$ | 23.360 | 12.961 | 40.580 | 22.169 | 11.791 | 38.420 |
| $\beta_{\phi,\text{Proposed}}$ | 10.358 | 6.865 | 15.461 | 9.984 | 6.577 | 14.964 |
| $\beta_{\zeta,\text{Proposed}}$ | 11.943 | 3.150 | 69.443 | 7.100 | 1.521 | 51.155 |
| $\beta_{\kappa,\text{Proposed}}$ | 67.527 | 1.132 | 5,103.709 | -76.205 | 0.012 | 2,531.192 |
| $\tau_{b0}$ | 0.729 | 0.433 | 1.233 | 0.670 | 0.403 | 1.178 |
| $\tau_{b\text{Proposed}}$ | 0.747 | 0.408 | 1.299 | 0.694 | 0.380 | 1.242 |
| $R_{12}$ | -0.879 | -0.986 | -0.414 | -0.948 | -0.998 | -0.524 |

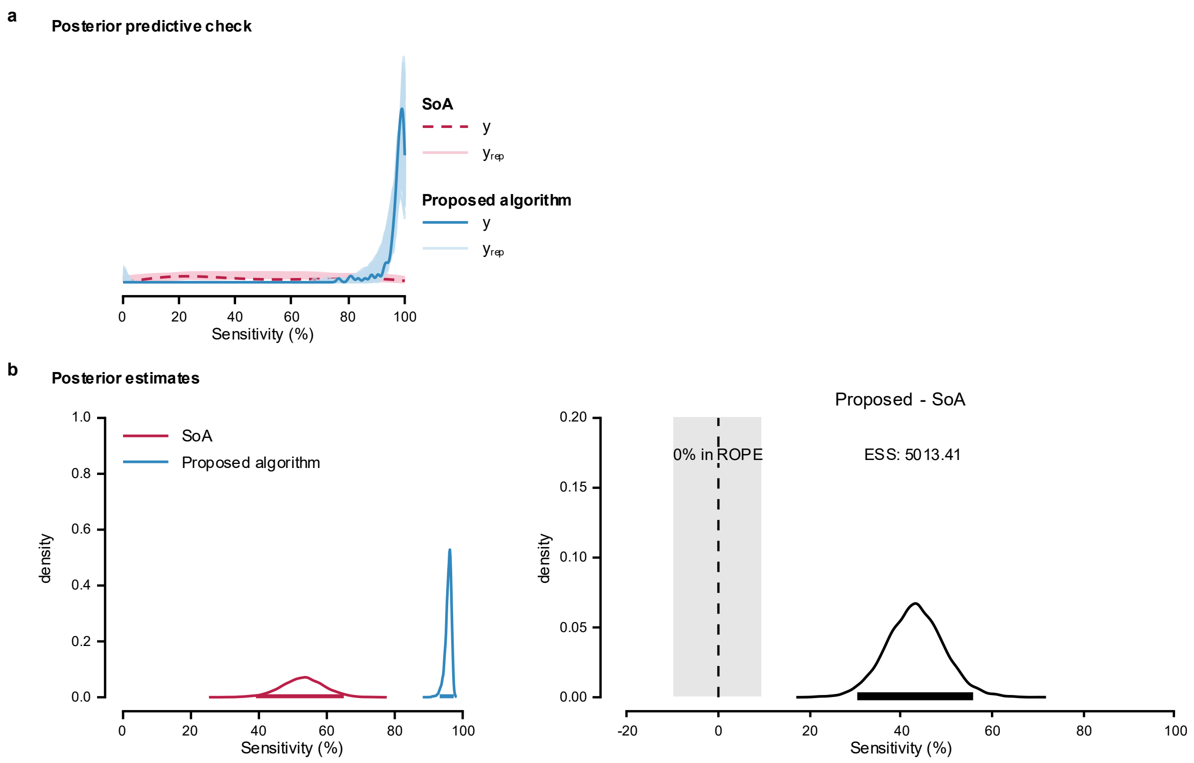


**Supplementary Fig. 45: Posterior predictive check and posterior estimates for Model 2 of firing identification sensitivity in tracked MG MUs across testing visits.**

**a**, Posterior predictive check of the model; **b**, Posterior distributions and the difference in means from Bayesian Model 2, used to compare the performance of the proposed algorithm and SoA method.

<u>Model 3</u>

The maximum PSRF across all parameters was 1.006, indicating good convergence, while the minimum ESS was 719.030, indicating stable estimates of the bounds of the credible intervals (Supplementary Table 65). Posterior summaries are reported as medians with 95% ETIs and 95% HDIs (Supplementary Table 66). Group-level correlations are reported on the correlation scale.

Posterior predictive checks were performed by overlaying model-generated predictions on the observed data (Supplementary Fig. 46a). The posterior distributions (Supplementary Fig.

46b) indicated that the proposed algorithm achieved a higher sensitivity (96.1%, [93.7-97.6%]) than the SoA algorithm (52.7%, [38.9-65.8%]). The posterior distribution of the difference excluded zero, and less than 2.5% of its distribution fell within ROPE.

**Supplementary Table 65: Posterior MCMC diagnostics for Model 3 of firing identification sensitivity in tracked MG MUs across testing visits.**

| | PSRF Pt | PSRF UpCI | ESS |
|---|---|---|---|
| $\alpha_{\mu}$ | 1.000 | 1.001 | 5,036.184 |
| $\alpha_{\phi}$ | 1.000 | 1.001 | 2,728.908 |
| $\alpha_{\zeta}$ | 1.000 | 1.001 | 1,4121.714 |
| $\alpha_{\kappa}$ | 1.000 | 1.000 | 2,1999.444 |
| $\beta_{\mu,\text{Proposed}}$ | 1.000 | 1.001 | 5,364.949 |
| $\beta_{\phi,\text{Proposed}}$ | 1.003 | 1.006 | 719.030 |
| $\beta_{\zeta,\text{Proposed}}$ | 1.000 | 1.001 | 1,4775.563 |
| $\beta_{\kappa,\text{Proposed}}$ | 1.000 | 1.000 | 2,2125.638 |
| $\tau_{b0}$ | 1.000 | 1.000 | 5,943.786 |
| $\tau_{b\text{Proposed}}$ | 1.000 | 1.000 | 6,944.338 |
| $\tau_{c}$ | 1.002 | 1.005 | 748.043 |
| $R_{12}$ | 1.001 | 1.002 | 9,714.394 |

**Supplementary Table 66: Posterior summaries for Model 3 of firing identification sensitivity in tracked MG MUs across testing visits.**

| | **50%** | **95% ETI** | | **Mode** | **95% HDI** | |
|---|---|---|---|---|---|---|
| | | **2.5%** | **97.5%** | | **low** | **high** |
| $\alpha_{\mu}$ | 0.532 | 0.393 | 0.660 | 0.534 | 0.394 | 0.661 |
| $\alpha_{\phi}$ | 3.266 | 2.417 | 4.705 | 3.118 | 2.294 | 4.514 |
| $\alpha_{\zeta}$ | 0.014 | 0.003 | 0.044 | 0.010 | 0.001 | 0.037 |
| $\alpha_{\kappa}$ | 0.238 | 0.011 | 0.897 | 0.054 | 0.001 | 0.839 |
| $\beta_{\mu,\text{Proposed}}$ | 24.620 | 13.294 | 46.154 | 23.050 | 11.575 | 42.878 |
| $\beta_{\phi,\text{Proposed}}$ | 11.111 | 7.067 | 34.400 | 10.098 | 5.757 | 25.570 |
| $\beta_{\zeta,\text{Proposed}}$ | 11.844 | 3.212 | 65.983 | 6.545 | 1.439 | 48.714 |
| $\beta_{\kappa,\text{Proposed}}$ | 67.131 | 1.159 | 5,480.474 | -74.184 | 0.045 | 2,587.436 |
| $\tau_{b0}$ | 0.755 | 0.447 | 1.293 | 0.693 | 0.416 | 1.231 |
| $\tau_{b\text{Proposed}}$ | 0.788 | 0.430 | 1.362 | 0.766 | 0.382 | 1.284 |
| $\tau_{c}$ | 0.217 | 0.009 | 0.761 | 0.052 | 0.000 | 0.697 |
| $R_{12}$ | -0.869 | -0.984 | -0.393 | -0.942 | -0.998 | -0.500 |

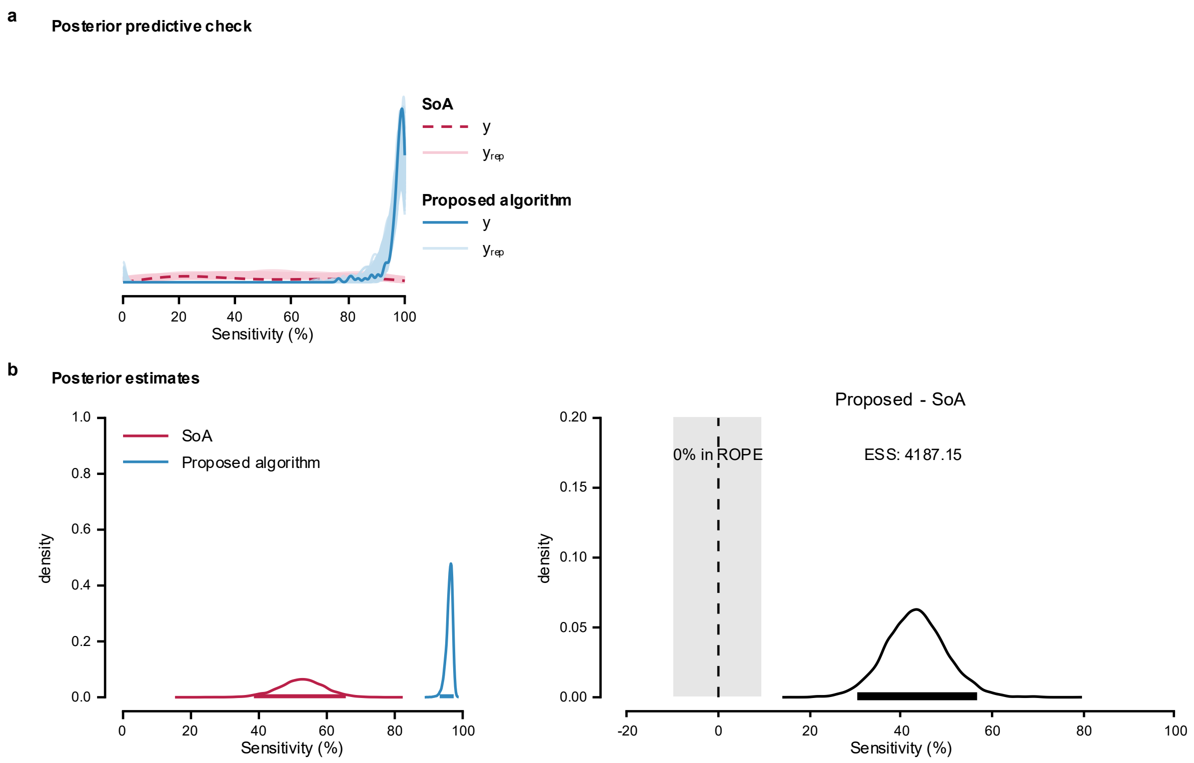


**Supplementary Fig. 46: Posterior predictive check and posterior estimates for Model 3 of firing identification sensitivity in tracked MG MUs across testing visits.**

**a**, Posterior predictive check of the model; **b**, Posterior distributions and the difference in means from Bayesian Model 3, used to compare the performance of the proposed algorithm and SoA method.

### MODEL COMPARISON

Model 2 showed the greatest expected predictive accuracy. ELPD diff relative to Model 2 were −1.9 (SE = 2.5) for Model 3 and −15.4 (SE = 4.7) for Model 1, suggesting similar predictive performance for Models 2 and 3 and reduced performance for Model 1.

**Supplementary Note 17. Comparison of firing identification precision in tracked MG MUs across testing visits.**

Model implementation, likelihood function, and priors are detailed in Supplementary Note 5. The full dataset contains firing identification precision values for 113 tracked MG MUs (hereafter referred to as 'precision'). The MUs firings were obtained by applying the MU filters calculated using the SoA algorithm[5] and the proposed algorithm to the HDsEMG signals obtained in the second testing visit. Distribution of the data is shown in Supplementary Fig. 47.

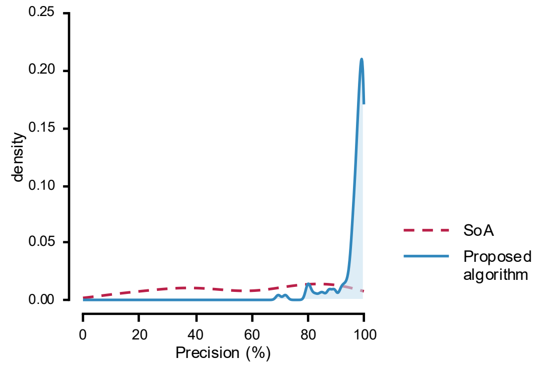


**Supplementary Fig. 47: Distribution of firing identification precision in tracked MG MUs across testing visits.**

Precision values were calculated from firing identification obtained by applying MU filters calculated using SoA (dashed line) and the proposed algorithms (solid line) to the HDsEMG signals generated with synthetic grid displacement.

In all models, ZOIB parameters were back transformed into their interpretable component scales: *mu*, *zoi*, and *coi* to probabilities, and *phi* to the positive precision scale. Group-level SDs are reported on their native model/link scale.

Model 1

The maximum PSRF across all parameters was 1.002, indicating good convergence, while the minimum ESS was 6,125.286, indicating stable estimates of the bounds of the credible intervals (Supplementary Table 67). Posterior summaries are reported in Supplementary Table 68 as medians with 95% ETIs and 95% HDIs. Group-level correlations are reported on the correlation scale.

Posterior predictive checks were performed by overlaying model-generated predictions on the observed data (Supplementary Fig. 48a). The posterior distributions (Supplementary Fig. 48b) indicated that the proposed algorithm achieved a higher precision (95.2%, [92.8-96.8 %]) than the SoA algorithm (57.9%, [50.4-64.7%]). The posterior distribution of the difference excluded zero, and less than 2.5% of its distribution fell within ROPE.

**Supplementary Table 67: Posterior MCMC diagnostics for Model 1 of firing identification precision in tracked MG MUs across testing visits.**

| | PSRF Pt | PSRF UpCI | ESS |
|---|---|---|---|
| $\alpha_{\mu}$ | 1.000 | 1.001 | 1,0126.198 |
| $\alpha_{\phi}$ | 1.000 | 1.001 | 2,1207.749 |
| $\alpha_{\zeta}$ | 1.000 | 1.000 | 1,5749.186 |
| $\alpha_{\kappa}$ | 1.000 | 1.000 | 2,4702.617 |
| $\beta_{\mu,\text{Proposed}}$ | 1 | 1.001 | 2,2631.534 |
| $\beta_{\phi,\text{Proposed}}$ | 1.000 | 1.001 | 2,0799.107 |
| $\beta_{\zeta,\text{Proposed}}$ | 1.000 | 1.000 | 1,6660.903 |
| $\beta_{\kappa,\text{Proposed}}$ | 1.000 | 1.000 | 2,5294.913 |
| $\tau_b$ | 1.001 | 1.002 | 5,537.584 |

**Supplementary Table 68: Posterior summaries for Model 1 of firing identification precision in tracked MG MUs across testing visits.**

| | 50% | 95% ETI | | Mode | 95% HDI | |
|---|---|---|---|---|---|---|
| | | 2.5% | 97.5% | | low | high |
| $\alpha_{\mu}$ | 0.581 | 0.505 | 0.649 | 0.585 | 0.508 | 0.651 |
| $\alpha_{\phi}$ | 2.789 | 2.158 | 3.555 | 2.747 | 2.119 | 3.501 |
| $\alpha_{\zeta}$ | 0.021 | 0.005 | 0.056 | 0.016 | 0.002 | 0.050 |
| $\alpha_{\kappa}$ | 0.537 | 0.070 | 0.950 | 0.613 | 0.081 | 0.958 |
| $\beta_{\mu,\text{Proposed}}$ | 15.539 | 11.579 | 20.600 | 15.279 | 11.388 | 20.304 |
| $\beta_{\phi,\text{Proposed}}$ | 6.868 | 4.512 | 10.240 | 6.595 | 4.380 | 10.013 |
| $\beta_{\zeta,\text{Proposed}}$ | 9.389 | 2.984 | 42.956 | 6.121 | 1.544 | 32.650 |
| $\beta_{\kappa,\text{Proposed}}$ | 17.654 | 0.580 | 700.510 | -14.805 | 0.018 | 351.051 |
| $\tau_b$ | 0.308 | 0.082 | 0.673 | 0.278 | 0.047 | 0.621 |

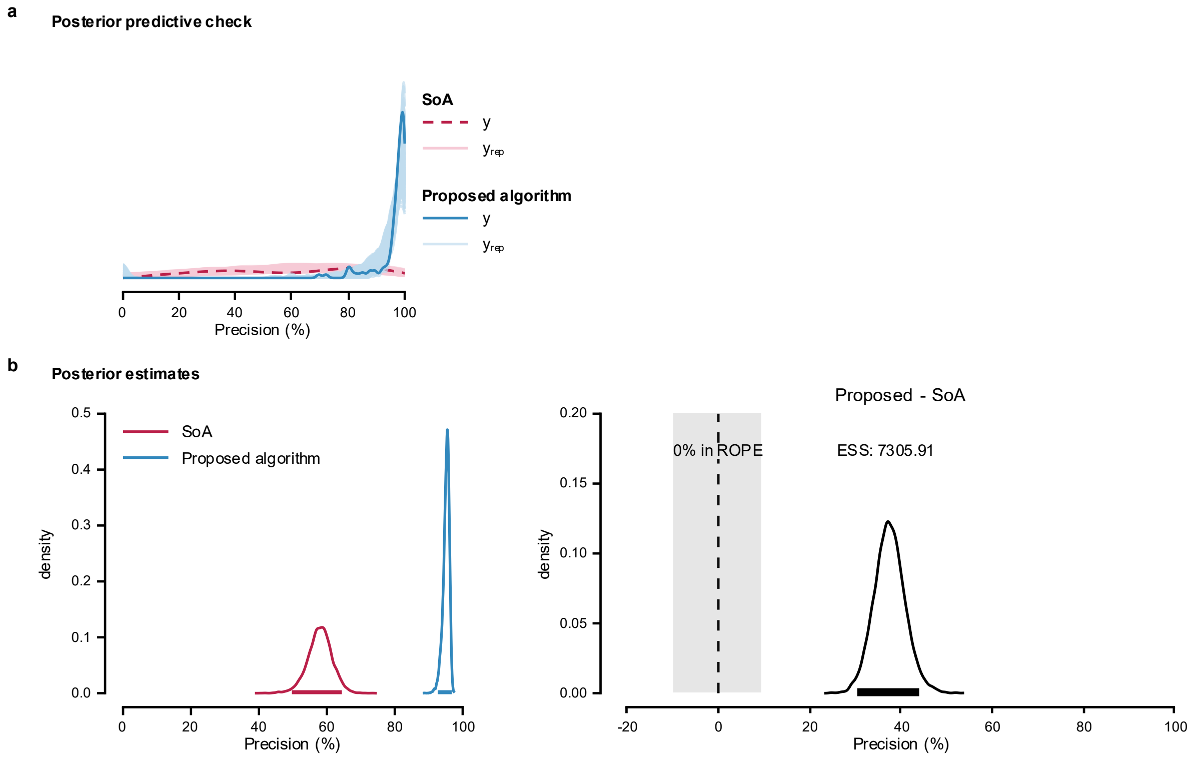


**Supplementary Fig. 48: Posterior predictive check and posterior estimates for Model 1 of firing identification precision in tracked MG MUs across testing visits.**
**a**, Posterior predictive check of the model; **b**, Posterior distributions and the difference between means estimated using Bayesian Model 1 to compare the performance of the proposed algorithm and SoA method.

<u>Model 2</u>

The maximum PSRF across all parameters was 1.002, indicating good convergence, while the minimum ESS was 6,162.999, indicating stable estimates of the bounds of the credible intervals (Supplementary Table 51). Posterior summaries are reported as medians with 95% ETIs and 95% HDIs (Supplementary Table 70). Group-level correlations are reported on the correlation scale.

Posterior predictive checks were performed by overlaying model-generated predictions on the observed data (Supplementary Fig. 49a). The posterior distributions (Supplementary Fig.

49b) indicated that the proposed algorithm achieved a higher precision (95.1%, [92.6-96.8%]) than the SoA algorithm (58.3%, [49.5-66.9%]). The posterior distribution of the difference excluded zero, and less than 2.5% of its distribution fell within ROPE.

**Supplementary Table 69: Posterior MCMC diagnostics for Model 2 of firing identification precision in tracked MG MUs across testing visits.**

| | PSRF Pt | PSRF UpCI | ESS |
|---|---|---|---|
| $\alpha_{\mu}$ | 1.000 | 1.001 | 9,898.987 |
| $\alpha_{\phi}$ | 1.001 | 1.001 | 1,5328.884 |
| $\alpha_{\zeta}$ | 1.000 | 1.001 | 1,8374.223 |
| $\alpha_{\kappa}$ | 1.000 | 1.000 | 2,5391.269 |
| $\beta_{\mu,\text{Proposed}}$ | 1.000 | 1.001 | 1,3683.989 |
| $\beta_{\phi,\text{Proposed}}$ | 1.000 | 1.001 | 2,0644.275 |
| $\beta_{\zeta,\text{Proposed}}$ | 1.000 | 1.001 | 1,9311.802 |
| $\beta_{\kappa,\text{Proposed}}$ | 1.000 | 1.000 | 2,5785.492 |
| $\tau_{b0}$ | 1.001 | 1.002 | 6,538.517 |
| $\tau_{b\text{Proposed}}$ | 1.001 | 1.002 | 6,162.999 |
| $R_{12}$ | 1.000 | 1.001 | 1,0492.527 |

**Supplementary Table 70: Posterior summaries for Model 2 of firing identification precision in tracked MG MUs across testing visits.**

| | 50% | 95% ETI | | Mode | 95% HDI | |
|---|---|---|---|---|---|---|
| | | 2.5% | 97.5% | | low | high |
| $\alpha_{\mu}$ | 0.585 | 0.494 | 0.671 | 0.582 | 0.494 | 0.670 |
| $\alpha_{\phi}$ | 2.983 | 2.283 | 3.856 | 2.946 | 2.236 | 3.788 |
| $\alpha_{\zeta}$ | 0.021 | 0.005 | 0.054 | 0.016 | 0.003 | 0.049 |
| $\alpha_{\kappa}$ | 0.536 | 0.069 | 0.948 | 0.645 | 0.090 | 0.966 |
| $\beta_{\mu,\text{Proposed}}$ | 15.079 | 9.975 | 21.620 | 14.671 | 9.698 | 21.270 |
| $\beta_{\phi,\text{Proposed}}$ | 6.488 | 4.250 | 9.735 | 6.421 | 4.033 | 9.392 |
| $\beta_{\zeta,\text{Proposed}}$ | 9.530 | 3.022 | 41.290 | 6.192 | 1.469 | 31.235 |
| $\beta_{\kappa,\text{Proposed}}$ | 17.023 | 0.569 | 707.389 | -14.814 | 0.023 | 369.925 |
| $\tau_{b0}$ | 0.443 | 0.159 | 0.880 | 0.420 | 0.136 | 0.840 |
| $\tau_{b\text{Proposed}}$ | 0.316 | 0.019 | 0.773 | 0.289 | 0.000 | 0.692 |
| $R_{12}$ | -0.601 | -0.951 | 0.467 | -0.825 | -0.992 | 0.288 |

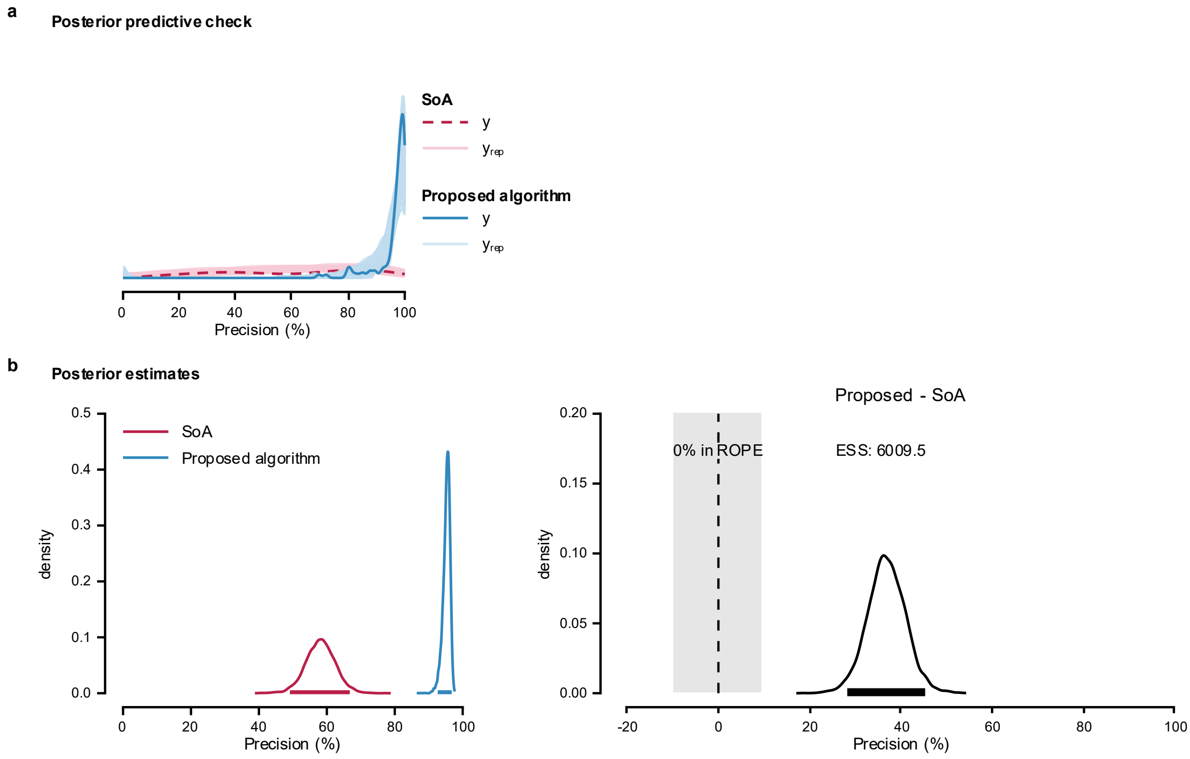


**Supplementary Fig. 49: Posterior predictive check and posterior estimates for Model 2 of firing identification precision in tracked MG MUs across testing visits.**

**a**, Posterior predictive check of the model; **b**, Posterior distributions and the difference between means estimated using Bayesian Model 2 to compare the performance of the proposed algorithm and SoA method.

<u>Model 3</u>

The maximum PSRF across all parameters was 1.002, indicating good convergence, while the minimum ESS was 4,792.538, indicating stable estimates of the bounds of the credible intervals (Supplementary Table 71). Posterior summaries are reported in Supplementary Table 72 as medians with 95% ETIs and 95% HDIs. Group-level correlations are reported on the correlation scale.

Posterior predictive checks were performed by overlaying model-generated predictions on the observed data (Supplementary Fig. 50a). The posterior distributions (Supplementary Fig.

50b) indicated that the proposed algorithm achieved a higher precision (95.2%, [92.6-96.8%]) than the SoA algorithm (58.3%, [49.2-66.8%]). The posterior distribution of the difference excluded zero, and less than 2.5% of its distribution fell within ROPE.

**Supplementary Table 71: Posterior MCMC diagnostics for Model 3 of firing identification precision in tracked MG MUs across testing visits.**

| | PSRF Pt | PSRF UpCI | ESS |
|---|---|---|---|
| $\alpha_{\mu}$ | 1.000 | 1.000 | 1,0768.934 |
| $\alpha_{\phi}$ | 1.000 | 1.001 | 1,2734.244 |
| $\alpha_{\zeta}$ | 1.000 | 1.000 | 1,9212.704 |
| $\alpha_{\kappa}$ | 1.000 | 1.000 | 2,8112.325 |
| $\beta_{\mu,\text{Proposed}}$ | 1.000 | 1.001 | 1,4713.291 |
| $\beta_{\phi,\text{Proposed}}$ | 1.000 | 1.000 | 2,2458.397 |
| $\beta_{\zeta,\text{Proposed}}$ | 1.000 | 1.000 | 2,0158.275 |
| $\beta_{\kappa,\text{Proposed}}$ | 1.000 | 1.000 | 2,7228.644 |
| $\tau_{b0}$ | 1.001 | 1.002 | 6,214.410 |
| $\tau_{b\text{Proposed}}$ | 1.000 | 1.001 | 6,278.603 |
| $\tau_{c}$ | 1.001 | 1.002 | 4,792.538 |
| $R_{12}$ | 1.000 | 1.001 | 1,1848.773 |

**Supplementary Table 72: Posterior summaries for Model 3 of firing identification precision in tracked MG MUs across testing visits.**

| | 50% | 95% ETI | | Mode | 95% HDI | |
|---|---|---|---|---|---|---|
| | | 2.5% | 97.5% | | low | high |
| $\alpha_{\mu}$ | 0.586 | 0.491 | 0.671 | 0.589 | 0.495 | 0.673 |
| $\alpha_{\phi}$ | 3.059 | 2.312 | 4.000 | 2.999 | 2.269 | 3.938 |
| $\alpha_{\zeta}$ | 0.021 | 0.005 | 0.055 | 0.016 | 0.003 | 0.049 |
| $\alpha_{\kappa}$ | 0.535 | 0.074 | 0.947 | 0.658 | 0.078 | 0.950 |
| $\beta_{\mu,\text{Proposed}}$ | 15.207 | 10.086 | 22.128 | 14.864 | 9.685 | 21.579 |
| $\beta_{\phi,\text{Proposed}}$ | 6.508 | 4.238 | 9.882 | 6.256 | 4.043 | 9.542 |
| $\beta_{\zeta,\text{Proposed}}$ | 9.504 | 3.002 | 41.608 | 6.014 | 1.587 | 31.326 |
| $\beta_{\kappa,\text{Proposed}}$ | 17.422 | 0.599 | 680.230 | 75.864 | 0.031 | 357.144 |
| $\tau_{b0}$ | 0.446 | 0.156 | 0.876 | 0.415 | 0.142 | 0.856 |
| $\tau_{b\text{Proposed}}$ | 0.318 | 0.020 | 0.786 | 0.327 | 0.000 | 0.699 |
| $\tau_{c}$ | 0.110 | 0.005 | 0.392 | 0.039 | 0.000 | 0.338 |
| $R_{12}$ | -0.595 | -0.952 | 0.467 | -0.816 | -0.991 | 0.288 |

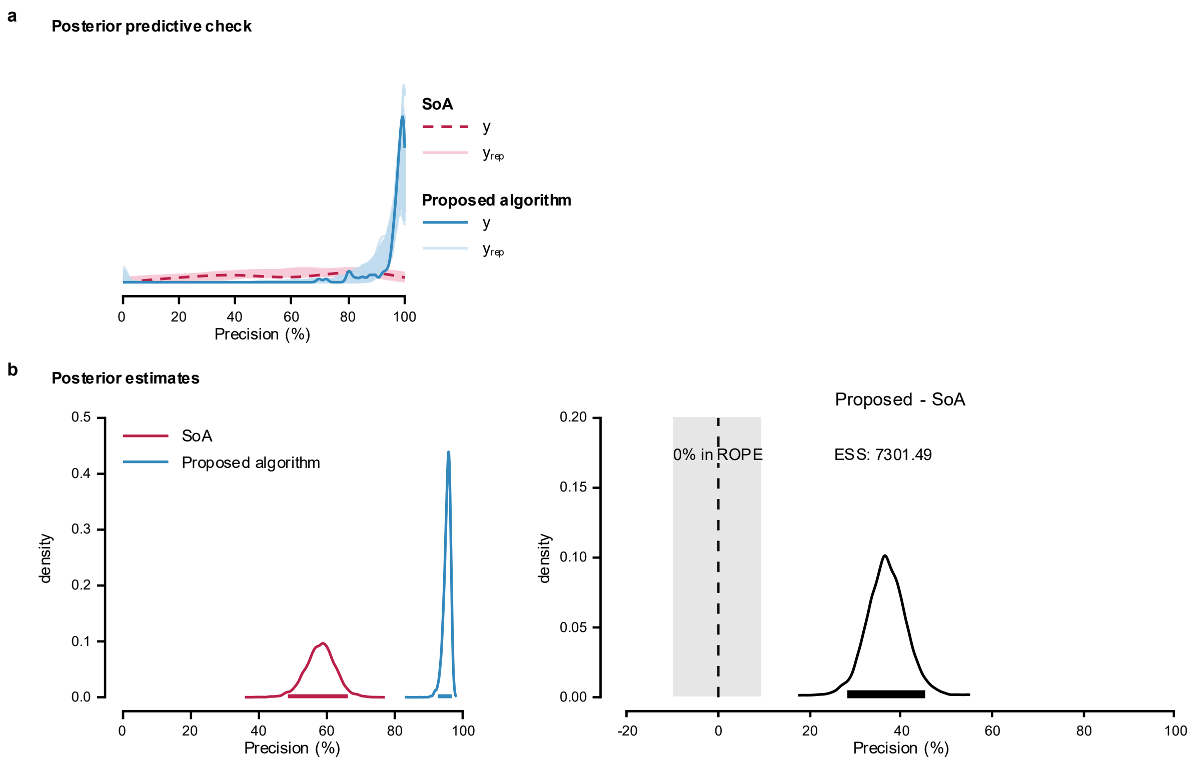


**Supplementary Fig. 50: Posterior predictive check and posterior estimates for Model 3 of firing identification precision in tracked MG MUs across testing visits.**

**a**, Posterior predictive check of the model; **b**, Posterior distributions and the difference between means estimated using Bayesian Model 1 to compare the performance of the proposed algorithm and SoA method.

MODEL COMPARISON

Model 2 showed the greatest expected predictive accuracy. ELPD diff relative to Model 2 were −0.7 (SE = 0.3) for Model 3 and −2.2 (SE = 1.5) for Model 1, suggesting similar predictive performance for all models.

**Supplementary Note 18. Bayesian analysis R session info**

```
R version 4.5.3 (2,026-03-11 ucrt)
Platform: x86_64-w64-mingw32/x64
Running under: Windows 10 x64 (build 1,9045)

Matrix products: default
  LAPACK version 3.12.1

locale:
[1] LC_COLLATE=English_United States.utf8
[2] LC_CTYPE=English_United States.utf8
[3] LC_MONETARY=English_United States.utf8
[4] LC_NUMERIC=C
[5] LC_TIME=English_United States.utf8

time zone: America/Chicago
tzcode source: internal

attached base packages:
[1] stats     graphics  grDevices utils     datasets  methods   base

other attached packages:
 [1] rstudioapi_0.18.0 future_1.70.0     tidyr_1.3.2       tibble_3.3.1      car_3.1-5
 [6] HDInterval_0.2.4  posterior_1.7.0   coda_0.19-4.1     readxl_1.5.0      loo_2.9.0
[11] carData_3.0-6     gridExtra_2.3     ggplot2_4.0.3     dplyr_1.2.1       brms_2.23.0
```

```
[16] bayesplot_1.15.0   Rcpp_1.1.1

loaded via a namespace (and not attached):
 [1] tidyselect_1.2.1     viridisLite_0.4.3    farver_2.1.2          S7_0.2.1
 [5] fastmap_1.2.0        lazyeval_0.2.3       TH.data_1.1-5         tensorA_0.36.2.1
 [9] digest_0.6.39        estimability_1.5.1   lifecycle_1.0.5       StanHeaders_2.32.10
[13] survival_3.8-6       processx_3.9.0       magrittr_2.0.5        compiler_4.5.3
[17] rlang_1.2.0          tools_4.5.3          utf8_1.2.6            data.table_1.18.2.1
[21] knitr_1.51           labeling_0.4.3       bridgesampling_1.2-1  htmlwidgets_1.6.4
[25] pkgbuild_1.4.8       plyr_1.8.9           RColorBrewer_1.1-3    multcomp_1.4-30
[29] abind_1.4-8          withr_3.0.2          purrr_1.2.2           future.apply_1.20.2
[33] stats4_4.5.3         inline_0.3.21        globals_0.19.1        emmeans_2.0.3
[37] scales_1.4.0         MASS_7.3-65          cli_3.6.6             mvtnorm_1.3-7
[41] crayon_1.5.3         generics_0.1.4       otel_0.2.0            RcppParallel_5.1.11-2
[45] grid_4.5.3           reshape2_1.4.5       httr_1.4.8            rstan_2.32.7
[49] stringr_1.6.0        splines_4.5.3        parallel_4.5.3        cellranger_1.1.0
[53] matrixStats_1.5.0    vctrs_0.7.3          Matrix_1.7-4          sandwich_3.1-1
[57] jsonlite_2.0.0       callr_3.7.6          Formula_1.2-5         listenv_0.10.1
[61] plotly_4.12.0        parallelly_1.47.0    glue_1.8.0            codetools_0.2-20
[65] gtable_0.3.6         QuickJSR_1.9.2       distributional_0.7.0  stringi_1.8.7
[69] pillar_1.11.1        htmltools_0.5.9      Brobdingnag_1.2-9     R6_2.6.1
[73] evaluate_1.0.5       lattice_0.22-9       backports_1.5.1       openxlsx_4.2.8.1
[77] rstantools_2.6.0     zip_2.3.3            nlme_3.1-168          checkmate_2.3.4
[81] xfun_0.57            zoo_1.8-15           pkgconfig_2.0.3
```

**Supplementary Note 19. Linear mixed effect model for the rate and recruitment threshold of tracked MUs across follicular phase**

The data contained 189 observations of MU firing rates (FR) and recruitment thresholds (RT) from 9 participants. The MUs were tracked across three visits during the follicular phase (EF1, EF2, and OV). FR and RT were analyzed separately. MUs that were only partially active within the 5 second window of stable torque production or were active outside that window, were not excluded from the analysis. The model was specified as follows

$$Y \sim \text{Visit} + (1 \mid \text{Subject:MUID}) \tag{S50}$$

where *Y* denotes the predicted variable (FR or RT) and MUID denotes the MU identification number. Model diagnostics were assessed visually using scaled Pearson residuals. Residual-versus-fitted plot and residuals by Visit were examined for systematic patterns and heteroscedasticity. Q-Q plots assessed the normality of residuals and MU-within-Subject conditional random effects, with 95 % pointwise confidence envelopes.

<u>Linear mixed effect model for the MU firing rates</u>

Standardized residuals were broadly centered around zero across fitted values, with no pronounced nonlinear pattern or evidence of heteroscedasticity, despite a slight upward trend (Supplementary Fig. 51). Residuals were similarly centered and dispersed across Visits, suggesting that this trend did not indicate substantial model misspecification. The Q–Q plot supported approximate residual normality: most points followed the theoretical quantiles and fell within the 95% pointwise confidence envelope, with only minor tail deviations. The model had a marginal $R^2$ of 0.009 and a conditional $R^2$ of 0.498. FR did not differ significantly across the follicular phase ($p = 0.180$, Supplementary Table 73). Response-scale estimated marginal means (EMM) for each Visit and model-scale pairwise comparison are provided in Supplementary Table 74 and Supplementary Table 75, respectively.

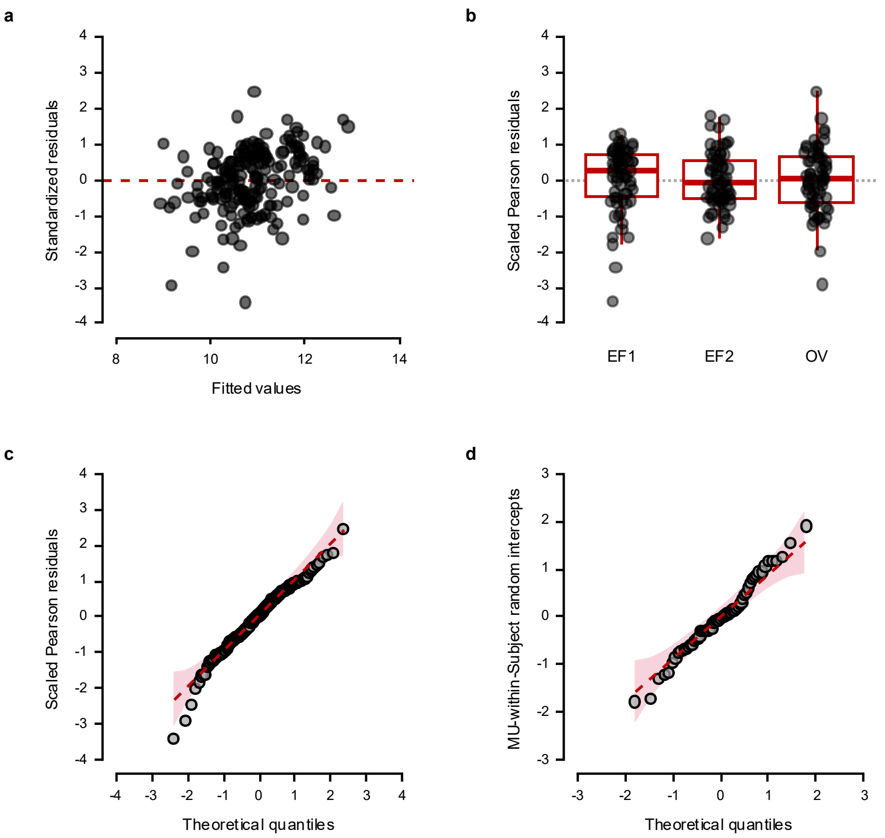


**Supplementary Fig. 51: LMM of tracked MUs FR diagnostics plots.**

**a**, Pearson residuals vs fitted values, **b**, residual by visit, **c**, Q-Q plot of scaled Pearson residuals, and **d**, Q-Q plot of random effects.

**Supplementary Table 73: Type III Analysis of Variance with Satterthwaite's method of the LMM of tracked MUs FR.**

| | Sum Sq | Mean Sq | NumDF | DenDF | F value | Pr(>F) |
|---|---|---|---|---|---|---|
| Visit | 2.745 | 1.372 | 2 | 124 | 1.738 | 0.18 |

**Supplementary Table 74: EMM in response scale for each visit of the LMM of tracked MUs FR.**

| Visit | EMM | SE | 95% CI |
|---|---|---|---|
| EF1 | 10.7 | 0.157 | 10.4 – 11.0 |
| EF2 | 11.0 | 0.157 | 10.7 – 11.3 |
| OV | 10.9 | 0.157 | 10.6 – 11.2 |

**Supplementary Table 75: Tukey-adjusted pairwise comparisons on the model scale of the LMM of tracked MUs FR.**

| contrast | estimate | SE | df | t ratio | p-value |
|---|---|---|---|---|---|
| EF1 – EF2 | -0.292 | 0.158 | 124 | -1.843 | 0.160 |
| EF1 – OV | -0.185 | 0.158 | 124 | -1.167 | 0.475 |
| EF2 – OV | 0.107 | 0.158 | 124 | 0.676 | 0.778 |

<u>Linear mixed effect model for the MU recruitment threshold</u>

A square root transformation was applied to the recruitment threshold because it improved the residual diagnostics (Supplementary Fig. 52). Standardized residuals were broadly centered around zero, with a slight upward trend. However, residuals by Visits showed no evidence of heteroscedasticity. The Q-Q plot indicated approximate normality: most scaled Pearson residuals closely followed the theoretical quantiles and fell within the 95% pointwise confidence envelope, with minor tail deviations. The model had a marginal $R^2$ of 0.059 and a conditional $R^2$ of 0.565.

Unlike the FR, RT showed a significant overall Visit effect in the Type III Analysis of Variance (ANOVA) using Satterthwaite's method (Supplementary Table 76), supported by pairwise comparisons ($p < 0.001$, Supplementary Table 77). Response-scale EMMs are provided in Supplementary Table 78.

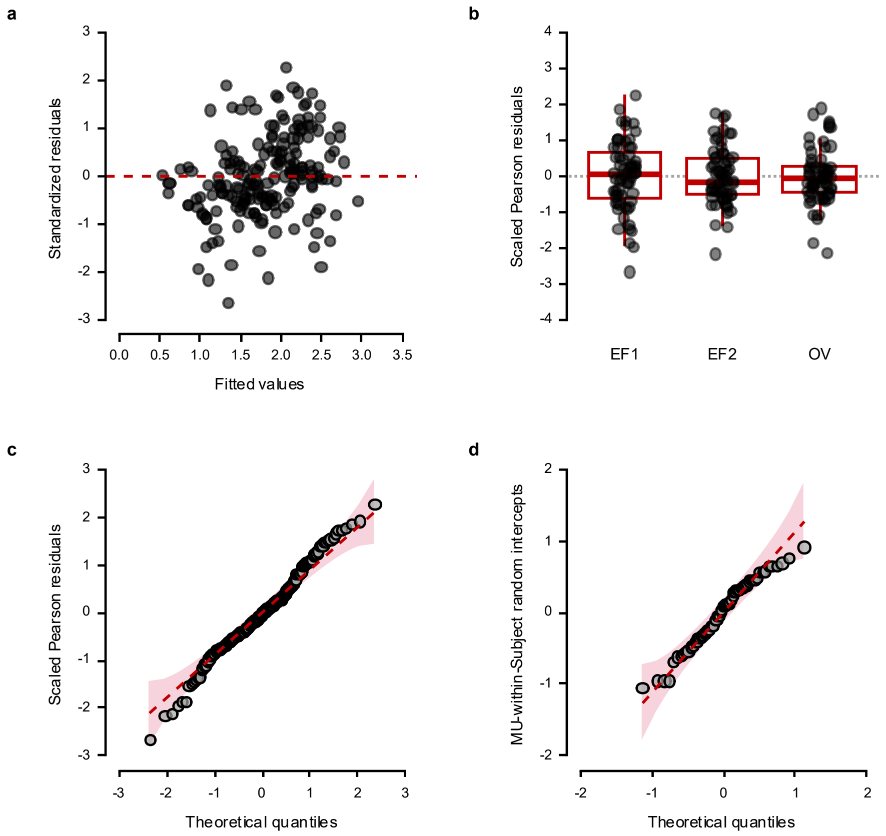


**Supplementary Fig. 52: LMM of tracked MUs RT diagnostics plots.**

**a**, Pearson residuals vs fitted values, **b**, residual by visit, **c**, Q-Q plot of scaled Pearson residuals, and **d**, Q-Q plot of random effects.

**Supplementary Table 76: Type III Analysis of Variance with Satterthwaite's method of the LMM of tracked MUs RT.**

| | Sum Sq | Mean Sq | NumDF | DenDF | F value | Pr(>F) |
|---|---|---|---|---|---|---|
| Visit | 6.400 | 3.200 | 2 | 124 | 12.684 | < 0.001* |

**Supplementary Table 77: Tukey-adjusted pairwise comparisons on the model scale of the LMM of tracked MUs RT.**

| contrast | estimate | SE | df | t ratio | p-value |
|---|---|---|---|---|---|
| EF1 – EF2 | 0.451 | 0.089 | 124 | 5.036 | < 0.001* |
| EF1 – OV | 0.219 | 0.089 | 124 | 2.448 | 0.041* |
| EF2 – OV | -0.232 | 0.089 | 124 | -2.588 | 0.029* |

**Supplementary Table 78: EMM in response scale for each visit of the LMM of tracked MUs RT.**

| Visit | EMM | SE | 95% CI |
|---|---|---|---|
| EF1 | 4.13 | 0.378 | 3.41 – 4.91 |
| EF2 | 2.50 | 0.294 | 1.95 – 3.12 |
| OV | 3.29 | 0.338 | 2.65 – 3.99 |

**Supplementary Note 20. Linear mixed effect model of firing rate and recruitment threshold of all identified motor units across follicular phase**

The data contained 393 observations of MU FR and RT from 9 participants. The MUs were tracked across three visits during the follicular phase (EF1, EF2, and OV). FR and RT were analyzed separately. MUs that were only partially active within the 5 second window of stable torque production or were active outside that window, were not excluded from the analysis. The model was specified as follows

$$Y \sim \text{Visit} + (1 \mid \text{Subject}) \qquad \text{(S51)}$$

where $Y$ is the predicted variable (FR or RT). Because MUs were not tracked across visits in all-observation analysis, MU-level repeated measure could not be specified as in the tracked MU case (Supplementary Note 19). This limited our ability to model within-MU correlation or MU-specific baseline differences across visits. Therefore, the model included Subject as the only random intercept, accounting for clustering of observations within Subject.

Model diagnostics were assessed visually using scaled Pearson residuals. Residual-versus-fitted plot and residuals by Visit were examined for systematic patterns and heteroscedasticity. Q-Q plots assessed the normality of residuals and within-Subject conditional random effects, with 95 % pointwise confidence envelopes.

<u>Linear mixed effect model for the MU firing rates</u>

Standardized residuals were broadly centered around zero across the range of fitted values, with no strong evidence of heteroscedasticity (Supplementary Fig. 53). Residuals showed similar spread across Visit, and the vertical bands were expected given the categorical predictor. The Q-Q plot supported approximate normality: most scaled Pearson residuals collowed the theoretical quantiles and fell within the 95% pointwise confidence envelope, with only minor tail deviations. The model had a marginal $R^2$ of 0.012 and a conditional $R^2$ of

0.067. The results suggest that the FR was not significantly different across the follicular phase (Supplementary Table 79). Response-scale EMMs and model scale pairwise are provided in Supplementary Table 80 and Supplementary Table 81, respectively.

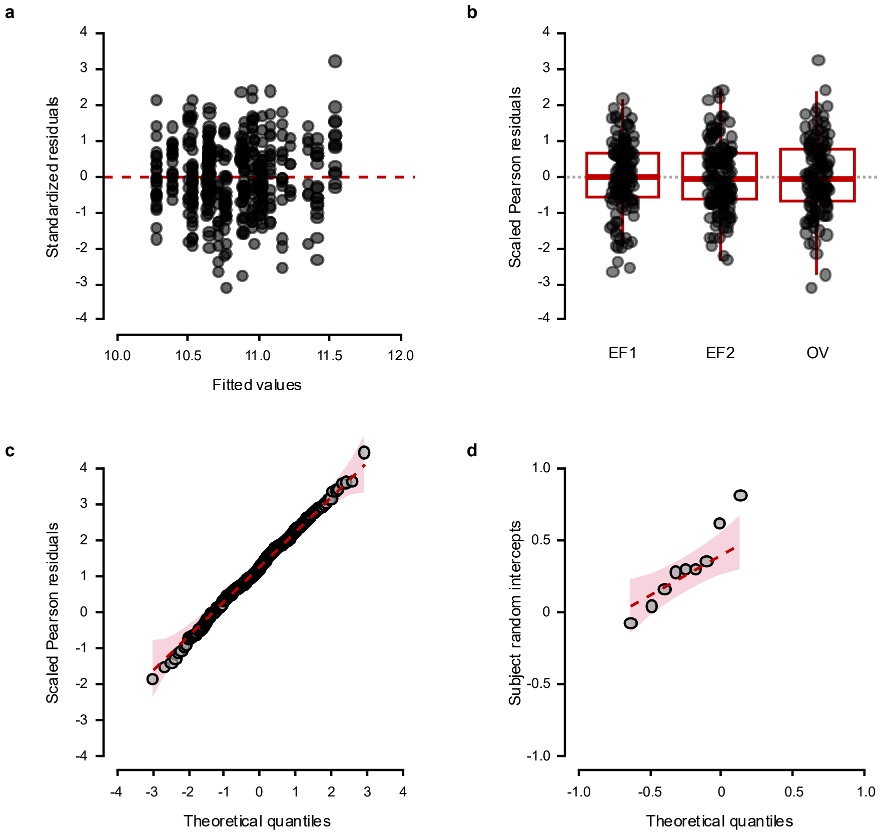


**Supplementary Fig. 53: LMM of all identified MUs FR diagnostics plots.**

**a**, Pearson residuals vs fitted values, **b**, residual by visit, **c**, Q-Q plot of scaled Pearson residuals, and **d**, Q-Q plot of random effects.

**Supplementary Table 79: Type III Analysis of Variance with Satterthwaite's method of the LMM of all identified MUs FR.**

| | Sum Sq | Mean Sq | NumDF | DenDF | F value | Pr(>F) |
|---|---|---|---|---|---|---|
| Visit | 9.081 | 4.540 | 2 | 384.29 | 2.494 | 0.084 |

**Supplementary Table 80: EMM in response scale for each visit of the LMM of all identified MUs FR.**

| Visit | EMM | SE | 95% CI |
|---|---|---|---|
| EF1 | 10.7 | 0.162 | 10.3 – 11.0 |
| EF2 | 10.9 | 0.159 | 10.6 – 11.2 |
| OV | 11.0 | 0.165 | 10.7 – 11.4 |

**Supplementary Table 81: Tukey-adjusted pairwise comparisons on the model scale of the LMM of all identified MUs FR.**

| contrast | estimate | SE | df | t ratio | p-value |
|---|---|---|---|---|---|
| EF1 – EF2 | -0.247 | 0.164 | 384 | -1.500 | 0.292 |
| EF1 – OV | -0.373 | 0.171 | 384 | -2.184 | 0.075 |
| EF2 – OV | -0.127 | 0.167 | 385 | -0.757 | 0.729 |

Linear mixed effect model for the MU recruitment threshold

A square root transformation was applied to the recruitment threshold because it improved the residual diagnostics (Supplementary Fig. 54). Standardized residuals were broadly centered around zero with similar spread across Visit. The Q-Q plot indicated generally reasonable normality, with mild upper-tail deviation. The model had a marginal $R^2$ of 0.020 and a conditional $R^2$ of 0.106.

Type III ANOVA with Satterthwaite's method revealed a significant overall Visit effect on RT (Supplementary Table 82) which was confirmed with the pairwise comparison (Supplementary Table 83). The EMM in response scale is provided in Supplementary Table 84.

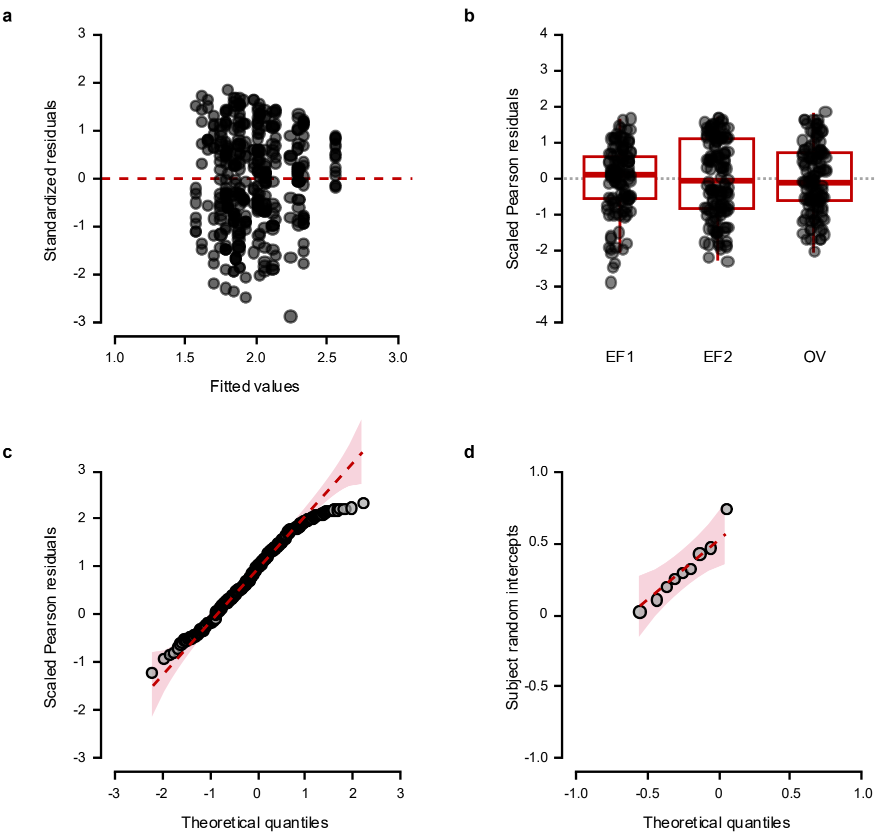


**Supplementary Fig. 54: LMM of all identified MUs RT diagnostics plots.**

**a**, Pearson residuals vs fitted values, **b**, residual by visit, **c**, Q-Q plot of scaled Pearson residuals, and **d**, Q-Q plot of random effects.

**Supplementary Table 82: Type III Analysis of Variance with Satterthwaite's method of the LMM of all identified MUs RT.**

| | Sum Sq | Mean Sq | NumDF | DenDF | F value | Pr(>F) |
|---|---|---|---|---|---|---|
| Visit | 5.256 | 2.628 | 2 | 383.28 | 4.367 | 0.013* |

**Supplementary Table 83: Tukey-adjusted pairwise comparisons on the model scale of the LMM of all identified MUs RT.**

| contrast | estimate | SE | df | t ratio | p-value |
|---|---|---|---|---|---|
| EF1 – EF2 | 0.228 | 0.095 | 383 | 2.413 | 0.043* |
| EF1 – OV | 0.264 | 0.098 | 383 | 2.688 | 0.020* |
| EF2 – OV | 0.036 | 0.096 | 384 | 0.374 | 0.926 |

**Supplementary Table 84: EMM in response scale for each visit of the LMM of all identified MUs RT.**

| Visit | EMM | SE | 95% CI |
|---|---|---|---|
| EF1 | 4.55 | 0.453 | 3.64 – 5.57 |
| EF2 | 3.63 | 0.399 | 2.82 – 4.54 |
| OV | 3.49 | 0.401 | 2.69 – 4.40 |
